# Metabolic capabilities are highly conserved among human nasal-associated *Corynebacterium* species in pangenomic analyses

**DOI:** 10.1101/2023.06.05.543719

**Authors:** Tommy H. Tran, Isabel F. Escapa, Ari Q. Roberts, Wei Gao, Abiola C. Obawemimo, Julia A. Segre, Heidi H. Kong, Sean Conlan, Matthew S. Kelly, Katherine P. Lemon

**Author notes:** Address correspondence to Katherine P. Lemon. Equal contribution (Isabel F. Escapa, Ari Q. Roberts). W.G. is now at Illumina Inc. (of San Diego, CA, USA) in Melbourne, Australia.

## Abstract

*Corynebact*e*rium* species are globally ubiquitous in human nasal microbiota across the lifespan. Moreover, nasal microbiota profiles typified by higher relative abundances of *Corynebacterium* are often positively associated with health. Among the most common human nasal *Corynebacterium* species are *C. propinquum*, *C. pseudodiphtheriticum, C. accolens*, and *C. tuberculostearicum*. To gain insight into the functions of these four species, we identified genomic, phylogenomic, and pangenomic properties and estimated the metabolic capabilities of 87 distinct human nasal *Corynebacterium* strain genomes: 31 from Botswana and 56 from the USA. *C. pseudodiphtheriticum* had geographically distinct clades consistent with localized strain circulation, whereas some strains from the other species had wide geographic distribution spanning Africa and North America. All species had similar genomic and pangenomic structures. Gene clusters assigned to all COG metabolic categories were overrepresented in the persistent versus accessory genome of each species indicating limited strain-level variability in metabolic capacity. Based on prevalence data, at least two *Corynebacterium* species likely coexist in the nasal microbiota of 82% of adults. So, it was surprising that core metabolic capabilities were highly conserved among the four species indicating limited species-level metabolic variation. Strikingly, strains in the USA clade of *C. pseudodiphtheriticum* lacked genes for assimilatory sulfate reduction present in most of the strains in the Botswana clade and in the other studied species, indicating a recent, geographically related loss of assimilatory sulfate reduction. Overall, the minimal species and strain variability in metabolic capacity implies coexisting strains might have limited ability to occupy distinct metabolic niches.

**IMPORTANCE:** Pangenomic analysis with estimation of functional capabilities facilitates our understanding of the full biologic diversity of bacterial species. We performed systematic genomic, phylogenomic, and pangenomic analyses with qualitative estimation of the metabolic capabilities of four common human nasal *Corynebacterium* species, along with focused experimental validations, generating a foundational resource. The prevalence of each species in human nasal microbiota is consistent with the common coexistence of at least two species. We identified a notably high level of metabolic conservation within and among species indicating limited options for species to occupy distinct metabolic niches, highlighting the importance of investigating interactions among nasal *Corynebacterium* species. Comparing strains from two continents, *C. pseudodiphtheriticum* had restricted geographic strain distribution characterized by an evolutionarily recent loss of assimilatory sulfate reduction in USA strains. Our findings contribute to understanding the functions of *Corynebacterium* within human nasal microbiota and to evaluating their potential for future use as biotherapeutics.

## INTRODUCTION

Nasal *Corynebacterium* species are frequently associated with health in compositional studies of human nasal microbiota. *Corynebacterium* are gram-positive bacteria in the phylum Actinobacteria (Actinomycetota). Based on studies from five continents, *Corynebacterium* species begin colonizing the human nasal passages before two months of age (1-13). *Corynebacterium* colonize both the skin-coated surface of the nasal vestibule (aka nostrils/nares) and the mucus-producing nasal respiratory epithelium coating the nasal passages posterior of the limen nasi through the nasopharynx (14-19). The bacterial microbiota of the human nasal passages from the nostrils through the nasopharynx is highly similar, and we refer to it herein as the human nasal microbiota.

Pediatric nasal microbiota profiles characterized by a high relative abundance of *Corynebacterium* are often associated with health rather than a specific disease or disease-risk state in children (1, 2, 4, 7, 11-13, 20-30). In young children, the genus *Corynebacterium* (alone or with the genus *Dolosigranulum*) is negatively associated with *Streptococcus pneumoniae* nasal colonization, which is important because *S*. *pneumoniae* colonization is a necessary precursor to invasive pneumococcal disease (13, 20, 22, 25, 28). For example, in young children in Botswana, the genus *Corynebacterium* is negatively associated with *S. pneumoniae* colonization both in a cross-sectional study of children younger than two years (25) and in a longitudinal study of infants followed from birth to one year of age (13). In contrast to these genus-level associations, little is known about species-level prevalence and relative abundance of nasal *Corynebacterium* in children. However, in a cultivation-based study, *C. pseudodiphtheriticum* is positively associated with ear and nasal health in young Indigenous Australian children (age 2-7 years), as is *D. pigrum* (29).

In adult nasal microbiota, the prevalence of the genus *Corynebacterium* is as high as 98.6%, with highly prevalent species including *C. accolens* (prevalence of 82%), *C. tuberculostearicum* (93%), *C. propinquum* (18%), and *C. pseudodiphtheriticum* (20%), based on 16S rRNA V1-V3 sequences (Tables S4A-B and S7 in (31)). In these data, 82% of the adult nostril samples contained ≥ 2 of these 4 *Corynebacterium* species, 30% contained ≥ 3, and 2.4% contained all 4 species. Thus, there is a high probability of coexistence of these *Corynebacterium* species in nasal microbial communities. Like children, some adults have nasal microbiota profiles characterized by a high relative abundance of *Corynebacterium* (18, 32). At least 23 validly published species of *Corynebacterium* can be cultivated from the adult nasal passages (17, 33). However, among these, *C. accolens* and *C. tuberculostearicum* (and/or other members of the *C. tuberculostearicum* species complex (34)) followed by *C. propinquum* and *C. pseudodiphtheriticum* are the most common in human nasal microbiota, in terms of both prevalence and relative abundance (15, 17, 31, 33). Indeed, the human nasal passages appear to be a primary habitat for *C. accolens, C. propinquum*, and *C. pseudodiphtheriticum,* whereas *C. tuberculostearicum* is also prevalent, often at high relative abundances, at other human skin sites (31, 34-37).

Nasal *Corynebacterium* interact with other common commensal/mutualistic nasal microbionts (38, 39) and with nasal pathobionts (22, 40-42). Some studies of healthy adult nasal microbiota report a negative association between *Staphylococcus aureus* and either the genus *Corynebacterium* or specific species of *Corynebacterium* (14, 31, 32, 43-47), although others do not (which might reflect strain-level variation and/or differences in populations studied). Furthermore, several small human studies support the potential use of *Corynebacterium* species to inhibit or eradicate pathobiont colonization of the human nasal passages (43, 48). This is of particular interest for *S. aureus* in the absence of an effective vaccine, since *S. aureus* nasal colonization increases the risk of invasive infection at distant body sites and the infection isolate matches the colonizing isolate in ∼80% of cases (49-52). Studies in mouse models further support potential benefits of nasal *Corynebacterium* for the prevention/treatment of respiratory syncytial virus and pneumococcal infections (53, 54). Inhibition of *S. pneumoniae* or *S. aureus in vitro* by nasal *Corynebacterium* species displays strain-level variation, highlighting the importance of sequencing the genomes of multiple strains per species (13, 14, 28, 55). An increasing number of these inhibitory interactions are characterized (22, 40, 41, 54).

Some interactions between *Corynebacterium* species and other nasal microbionts are related to metabolism/metabolites. For example, *C. accolens* strains secrete the triacylglycerol lipase LipS1 to hydrolyze host-surface triacylglycerols releasing nutritionally required free fatty acids that also inhibit *S. pneumoniae in vitro* (22, 54). Additionally, a positive metabolic interaction, such as cross-feeding, might be how *C. accolens*, *C. pseudodiphtheriticum,* and *C. propinquum* enhance the growth yield of the candidate mutualist *D. pigrum in vitro* (38). These examples highlight the importance of sequencing the genomes of multiple strains of each species to elucidate their metabolic capabilities, and the variation, that might influence interspecies interactions and contribute to promoting health-associated nasal microbiome compositions.

Overall, their ubiquity, frequent positive associations with health, and potential therapeutic use raise fundamental questions about the role of *Corynebacterium* species in human nasal microbiota. To increase genomic and metabolic knowledge of these, we performed systematic phylogenomic and pangenomic analyses of four common human nasal-associated *Corynebacterium* species. To increase the generalizability of our findings, we analyzed genomes of 87 nasal strains collected across two continents from both children and adults. Nasal strains of *C. pseudodiphtheriticum* overwhelmingly partitioned into clades by country of origin, consistent with geographically restricted strain circulation. Comparison of the core versus accessory genome of each of these four *Corynebacterium* species demonstrated that all COG categories associated with metabolism were enriched in the core genome, indicating limited strain-level metabolic variation within each species. To provide broader context, we compare the predicted metabolic abilities of nasal *Corynebacterium* species to two well-studied *Corynebacterium* species, *C. diphtheriae* and *C. glutamicum*, and to common nasal species from other bacterial genera. Metabolic estimation revealed that these four species share the majority of KEGG modules with few species-specific, or even clade-specific, metabolic abilities. However, we found that the clade of *C. pseudodiphtheriticum* dominated by strains from the USA lacked the module for assimilatory sulfate reduction, which is key for biosynthesis of sulfur-containing amino acids, and two representative USA strains were unable to grow under conditions requiring sulfate assimilation. We also validated the predictions that *C. tuberculostearicum,* alone of the four nasal species, accumulated intracellular glycogen and that *C. tuberculostearicum* and *C. accolens* generated all 20 amino acids since both grew in their absence.

## RESULTS

### *Corynebacterium pseudodiphtheriticum* displays geographically restricted strain circulation

To compare the genomic content and phylogenomic relationships among and within four *Corynebacterium* species commonly found in human nasal microbiota, we isolated strains from the nasal vestibule (nostrils) of generally healthy children and adults in the USA and from nasopharyngeal swabs of mother-infant pairs in Botswana. We compared 87 distinct nasal strain genomes to publicly available genomes of the type strain plus several other reference strains of each species, for a total of 20 reference genomes (39, 56-60) (**Table S1A**).

To confidently assign each new nasal isolate to a species, we first generated a maximum-likelihood phylogenomic tree based on the 632 single-copy core gene clusters (GCs) shared by the 107 strain genomes (**Fig. S1A**) and determined that each new nasal isolate was in a clade with the type strain of one of the nasal species (**Fig. S1B**; type strains in bold). *C. macginleyi* is the closest relative of *C. accolens* and these two species are challenging to distinguish by partial 16S rRNA gene sequences. Therefore, we included three *C. macginleyi* genomes in this phylogenomic analysis to confidently assign candidate *C. accolens* strains to a species. This five-species phylogenomic tree contained two major clades (**Fig. S1B**) confirming that *C. propinquum* and *C. pseudodiphtheriticum* are more closely related to each other, whereas *C. macginleyi*, *C. accolens*, and *C. tuberculostearicum* are more closely related to each other, with *C. macginleyi* closest to, yet distinct from *C. accolens.* Furthermore, these two major clades are distinctly separate from each other in a broader phylogenomic representation of the genus *Corynebacterium* (**Fig. S1C**, **Table S1C**). Next, we confirmed that each strain had a pairwise average nucleotide identity (ANI) of ≥ 95% for core GCs compared to the type strain of its assigned species (**Fig. S2A**). For each species, the pairwise ANIs for core GCs were very similar to those for all shared CoDing Sequences (CDS) (**Fig. S2B**).

To assess the evolutionary relationships between nasal isolates from both the USA and Botswana, we produced individual maximum-likelihood phylogenomic trees for each species (**Fig. 1**) based on its conservative core genome (**Fig. S2C**). These species-specific phylogenies provided a refined view of the relationships between strains based on the larger number of shared single-copy core GCs within each species (ranging from 1345 to 1788). To better approximate the root of each species-specific tree (**Fig. 1**), we used the type strain of the most closely related species in the multispecies phylogenomic trees (**Fig. S1**) as the outgroup (**Fig. S2D**). With a relatively even representation of Botswana and USA strains (40% vs. 58%), the phylogenomic tree for *C. pseudodiphtheriticum* had two large, well-supported clades dominated respectively by nasal strains from Botswana (15/15) or from the USA (20/22), indicating a restricted distribution of strains by country (**Fig. 1B**). We avoided calculating geographic proportions within major clades for *C. propinquum* (**Fig. 1A**) and *C. accolens* (**Fig. 1C**) because of the disproportionately high representation of USA strains (80%) and for *C. tuberculostearicum* (**Fig. 1D**) because there were only 6 nasal strains, 5 of which were from Botswana. Within these limitations, the phylogenomic analysis of these three species revealed some remarkably similar strains present in samples collected in the USA and Botswana based on their residing together in terminal clades. This raises the possibility that a subset of strains from each of these three species might have a wide geographic distribution, spanning from at least Botswana to Massachusetts.

**Figure 1.**
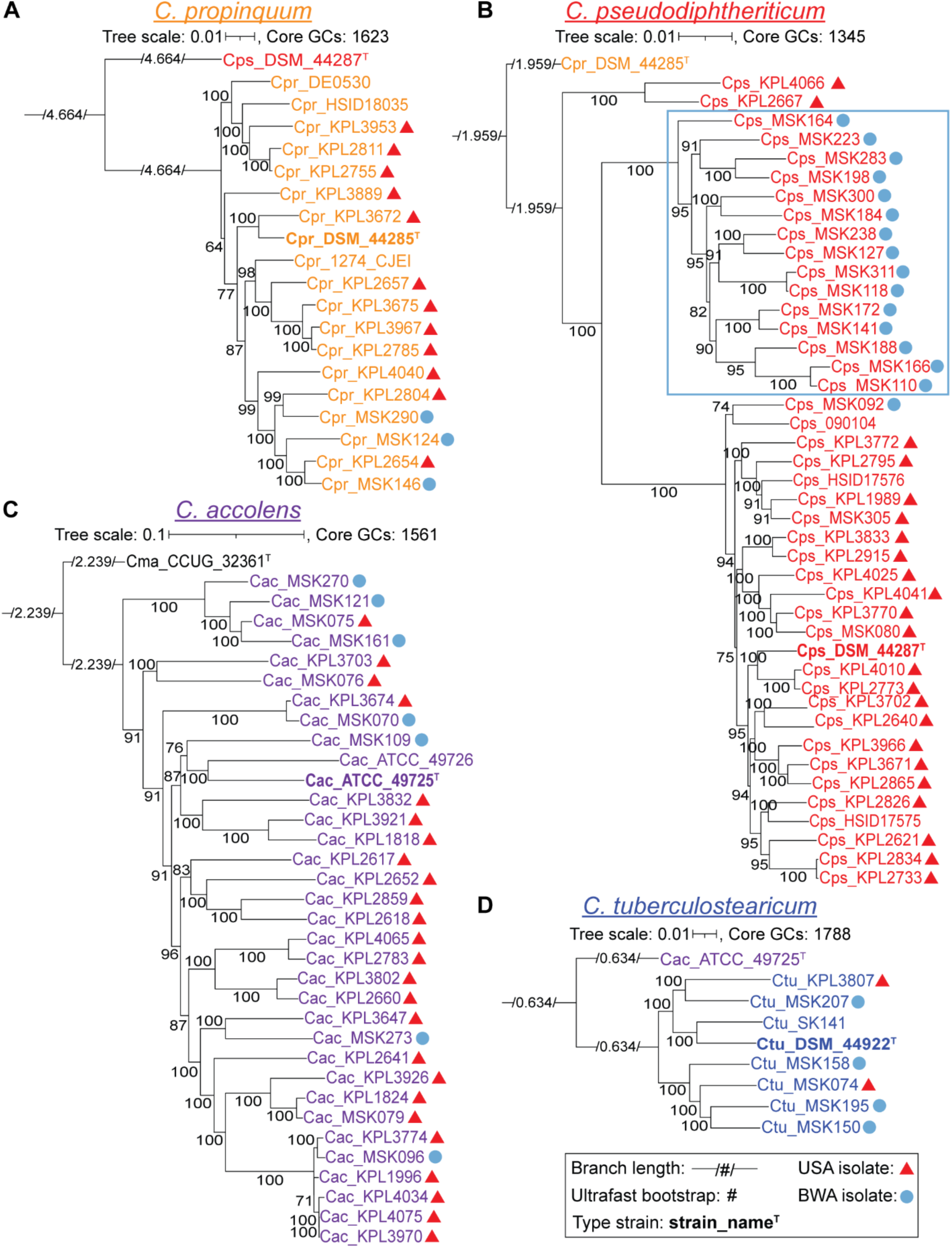
Species-specific phylogenomic trees show a geographic pattern of clades for *C. pseudodiphtheriticum*. Each panel shows a core-genome-based maximum-likelihood species-specific phylogeny. The majority (86%) of the MSK-named strains are from Botswana (blue circles), whereas all KPL-named strains are from the USA (red triangles). (**A**) Phylogeny of 19 *C. propinquum* strains based on 1,623 core GCs shows two major clades (BIC value 9762417.2123). (**B**) Phylogeny of 43 *C. pseudodiphtheriticum* strains based on 1,345 core GCs shows three major clades, one of which is entirely composed of strains from Botswana (15/15, outlined in light blue), whereas the other two have a majority of the USA nasal strains (red triangles), with 2/2 and 20/22, respectively (BIC value 10177769.6675). The branching pattern separating the Botswana and USA clades was well-supported with ultrafast bootstrap values ≥ 95% (61). (**C**) Phylogeny of 34 *C. accolens* strains based on 1,561 core GCs with the majority collected in the USA shows most Botswanan strains dispersed throughout (BIC value 10700765.2332). (**D**) Phylogeny of eight *C. tuberculostearicum* strains based on 1,788 core GCs with 6 nasal isolates from Botswana and the USA (BIC value 10452720.3067). For each species-specific phylogeny, the type strain from the most closely related species (**Fig. S1B**) serves as the outgroup. Each phylogeny was made from all shared conservative core GCs for a given species (**Fig. S2C**), including the subset of GCs that were absent in the corresponding outgroup (**Fig. S2D**), to provide the highest possible resolution among the strains within each species. A large majority of the branches have highly supported ultrafast bootstrap values with the lowest at 64 on an ancestral branch in the *C. propinquum* phylogeny. Type strains are indicated in bold with a superscript T. Ancestral branch lengths are indicated numerically within a visually shortened branch to fit on the page. Phylogenies were generated with IQ-Tree v2.1.3 using model finder, edge-linked-proportional partition model, and 1,000 ultrafast rapid bootstraps.

### The sizes of the core genomes of four common nasal *Corynebacterium* species have leveled off

Based on rarefaction analysis, the core genomes of *C. propinquum, C. pseudodiphtheriticum, C. accolens,* and *C. tuberculostearicum* have reached a stable size and are unlikely to decrease much further with the sequencing of additional strains (**Fig. 2Ai-Aiv**). Based on the respective Tettelin curves (red line), the *C. tuberculostearicum* core genome stabilized first at ∼7 genomes; however, with the fewest genomes at 8 this might be an upper bound that will continue to decrease with additional strain genomes. In comparison, *C. pseudodiphtheriticum* had the largest number of strain genomes at 43, with pairwise ANIs of ≥ 96.2% (**Fig. S2Aii**), and its core genome stabilized last at ∼37 genomes.

**Figure 2.**
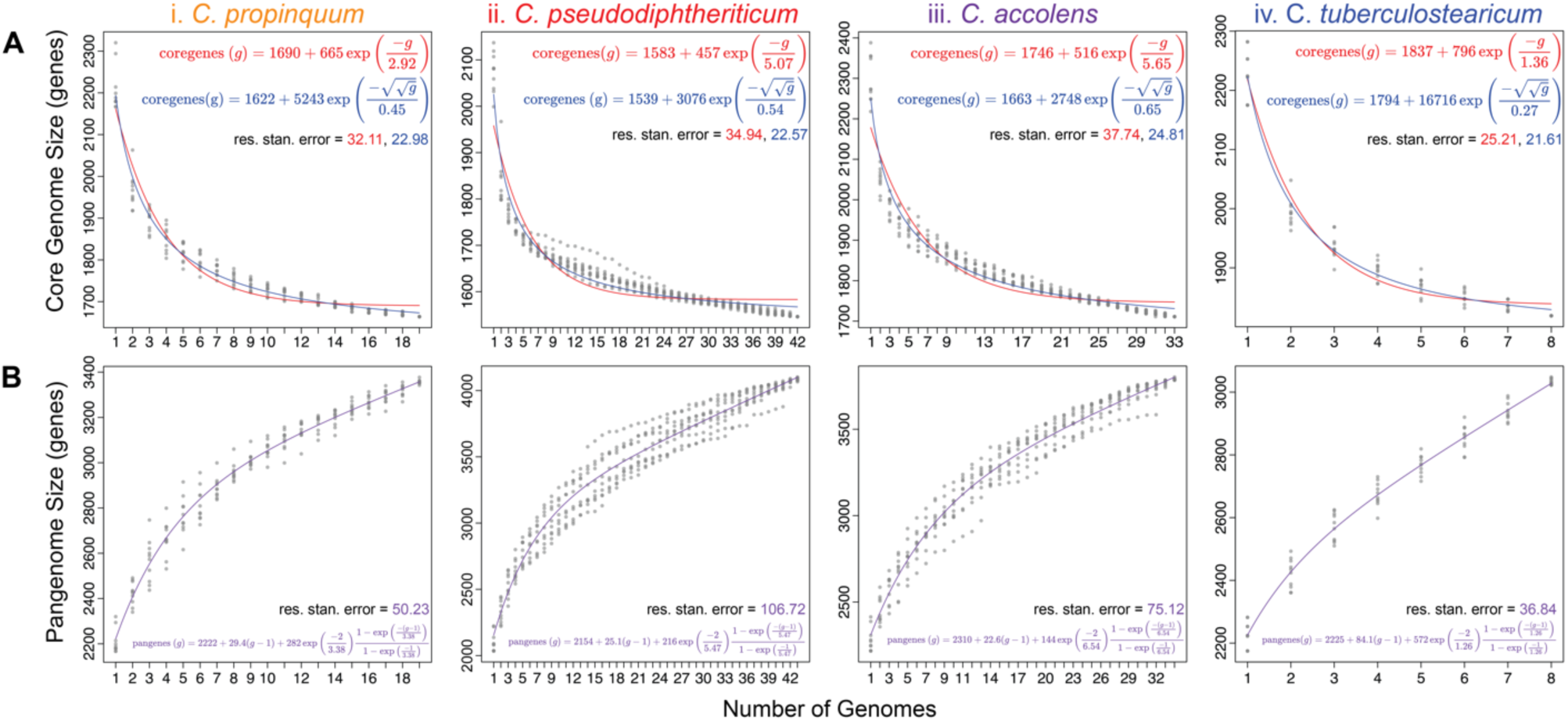
The four nasal *Corynebacterium* species have core genomes that have leveled off and pangenomes that remain open. **(A)** All four species have a core genome that has leveled off using a Tettelin curve fit model. (Ai) The *C. propinquum* core genome (n = 19) leveled off at ∼12 genomes. (Aii) The *C. pseudodiphtheriticum* core genome (n = 42) leveled off at ∼19 genomes. (Aiii) The *C. accolens* core genome (n = 33) leveled off at ∼ 21 genomes. (Aiv) The *C. tuberculostearicum* core genome (n = 8) leveled off at ∼7 genomes. Two best fit curve line models are shown for the core genome: Tettelin (red) and Willenbrock (blue). (B) The pangenomes for the four *Corynebacterium* species (i-vi) remain open as indicated by the continuous steep slope of the best fit line shown in purple. Core and pangenome size estimations were calculated from 10 random genome samplings (represented by gray dots) using the OMCL algorithm predicted GCs with GET_HOMOLOGUES v24082022.

The proportion of an individual genome of each of these nasal *Corynebacterium* species devoted to conservative core GCs ranged from 72% for *C. pseudodiphtheriticum* (1517/2105) to 79% for *C. tuberculostearicum* (1788/2250). This is based on the average number of CDS per genome (**Table 1**). The average / median genome size for each species ranged from ∼2.33 / 2.33 Mb for *C. pseudodiphtheriticum* to ∼2.51 / 2.52 Mb for *C. propinquum* with the average / median predicted CDS per genome ranging from 2105 / 2096 for *C. pseudodiphtheriticum* to 2265 / 2272 for *C. propinquum* (**Table 1**). These sizes and proportions are consistent with the reduced genome size and the GCs per genome of host-associated compared to environment-associated *Corynebacterium* species. For example, Swaney et al. report that environmental *Corynebacterium* species have a larger median genome size of 3.03 Mb and more GCs per genome (with an average of 2664) compared to host-associated *Corynebacterium* species (61).

**Table 1.** Basic genomic information for four common human nasal-associated *Corynebacterium* species.

| *Corynebacterium species* | # Strain Genomes (# Nasal isolates) | Average (median) genome size (Mb) | Average (median) CDS/genome | Average (median) G+C% | Conservative Core GCs/species<sup>a</sup> | Pangenome GCs/species<sup>b</sup> | Country #USA/# BWA | Age range (years) of nasal strains |
| --- | --- | --- | --- | --- | --- | --- | --- | --- |
| *C. propinquum* | 19 (15) | 2.51 (2.52) | 2265 (2272) | 56.47 (56.48) | 1623 | 3777 | 12/3 | 0.3-72 |
| *C. pseudodiphtheriticum*<sup>c</sup> | 43 (39) | 2.33 (2.33) | 2105 (2096) | 55.29 (55.29) | 1345 | 4590<sup>c</sup> | 22/17 | 0.2-55 |
| *C. accolens*<sup>d</sup> | 34<br>(32) | 2.50<br>(2.49) | 2304<br>(2294) | 59.46<br>(59.44) | 1561 | 4220<sup>d</sup> | 25/7 | 0.1-62 |
| *C. tuberculo*<br>*stearicu*<br>*m* | 8<br>(6) | 2.39<br>(2.39) | 2250<br>(2253) | 59.86<br>(59.88) | 1788 | 3232 | 2/4 | 0.1-39 |

<sup>a</sup>GET\_HOMOLOGUES conservative core GCs predicted from the consensus of BDBH, OMCL, and COGS algorithms.
<sup>b</sup>GET\_HOMOLOGUES pangenome GCs predicted from the consensus of OMCL, and COGS algorithms.
<sup>c</sup>Cps\_090104 removed from dataset for this analysis due to false % core lower bound.
<sup>d</sup>Cac\_ATCC\_49756 removed from dataset for this analysis due to false % core lower bound.

### The pangenomes of these four human nasal-associated *Corynebacterium* species remain open

With the number of strain genomes analyzed (**Table 1**), the pangenome of each of the four species continued to increase with each additional new genome, indicating that all are open (**Fig. 2Bi-Biv**). Parameters used to generate a pangenome via rarefaction yielded an overly conservative estimate of its size in GCs. Therefore, we used two other approaches to estimate the number of GCs in the pangenome for each species. These pangenome composition estimates are a lower bound for each species and will increase with sequencing of additional strain genomes. Starting with GET_HOMOLOGUES, we estimated pangenome size using the COG triangle and OMCL clustering algorithms. The pangenome size and its proportion contributed by core versus accessory GCs for each species ranged from 3232 GCs with 56% core and 44% accessory for *C. tuberculostearicum* to 4590 GCs with 33% core and 67% accessory for *C. pseudodiphtheriticum* (**Table 2**). The 56% core percentage for *C. tuberculostearicum* is likely an overestimate since this pangenome is based on only 8 genomes. This range of 33% to 56% for core genes per pangenome is similar to estimates for other human upper respiratory tract microbionts, such as *D. pigrum* (31%) (62), *S. aureus* (36%), and *Streptococcus pyogenes* (37%) (63).

**Table 2.** Pangenomic estimation of human nasal-associated *Corynebacterium* species based on three different platforms.

| Platform<br>Species | Pangenome size<br>(GCs) | % Core<br>GCs/pangenome | % Accessory<br>GCs/pangenome |
| --- | --- | --- | --- |
| **GET_HOMOLOGUES<sup>a</sup>** |  |  |  |
| *C. propinquum* | 3777 | 43% | 57% |
| *C. pseudodiphtheriticum<sup>c</sup>* | 4590 | 33% | 67% |
| *C. accolens<sup>d</sup>* | 4220 | 40% | 60% |
| *C. tuberculostearicum* | 3232 | 56% | 44% |
| **anvi'o** |  |  |  |
| *C. propinquum* | 3108 | 59% | 40% |
| *C. pseudodiphtheriticum* | 3590 | 48% | 51% |
| *C. accolens* | 3427 | 57% | 42% |
| *C. tuberculostearicum* | 2907 | 66% | 32% |
| **PPanGGOLiN** |  | % Persistent |  |
| *C. propinquum* | <sup>b</sup> | 63% | 37% |
| *C. pseudodiphtheriticum* | <sup>b</sup> | 49% | 51% |
| *C. accolens* | <sup>b</sup> | 59% | 41% |
| *C. tuberculostearicum* | <sup>b</sup> | 69% | 31% |

<sup>a</sup>GET\_HOMOLOGUES pangenome size, % core, and % accessory are from the consensus of OMCL and COGS algorithms.
<sup>b</sup>Pangenome size was estimated in anvi'o then the GCs imported into PPanGGOLiN to estimate persistent vs. accessory genome percentages.
<sup>c</sup>Cps\_090104 was removed from dataset only for this analysis due to an aberrant % core lower bound.
<sup>d</sup>Cac\_ATCC\_49756 was removed from dataset only for this analysis due to an aberrant % core lower bound.

Next, we used anvi’o to estimate the core- and pangenomes (64). The number of GCs in the core genome of each species estimated with GET_HOMOLOGUES was within 6-13% of those estimated with anvi’o; however, the GET_HOMOLOGUES estimated pangenome sizes were 11-24% larger (**Fig. S3A, File S1**). Consistent with this, the estimated single-copy core as a proportion of the pangenome using anvi’o was higher for each species ranging from 41% to 64% (**Fig. S3B**).

We also used anvi’o to visualize the strain-level variation in gene presence and absence within the four human nasal-associated *Corynebacterium* species (**Fig. S3B**). Manually arraying the genomes in anvi’o to correspond with their species-specific phylogenomic tree (**Fig. 1**) showed that some blocks of gene presence/absence correlated with the core-genome-based phylogenetic relationships among strains, but others did not (**Fig. S3B**). This is consistent with gene gain and loss playing a role in strain diversification with some of this due to mobile genetic elements and horizontal gene transfer (65, 66).

### Gene clusters assigned to the COG categories associated with metabolism are highly enriched in the core genomes of common nasal *Corynebacterium* species

To predict and compare functions based on the pangenomes of each species, we assigned GCs to COG categories and used PPanGGOLiN to define the persistent versus the accessory genome (**Table S2**) (67). We used the PPanGGOLiN estimation of the persistent genome rather than the traditionally defined core genome for this analysis because PPanGGOLin limits the effect of technical artifacts and/or strain-level gene loss events on the assessment of persistently shared gene clusters by considering the genomic context via examining genetic contiguity. As is common in bacteria, only about 63-65% of the GCs in the persistent genome and 26-36% of the GCs in the accessory genome of each species had an informative assignment to a definitive COG category (**Figs. 3Ai-iv & S4Ai-iv**). There was also variability in the size of the accessory genome among strains within each species (**Fig. S4A-B**). We next generated functional enrichment plots for COG categories in the persistent versus the accessory genome of each species (**Fig. 3Bi-vi**). GCs assigned to “Mobilome: prophages, transposons” (mobile genetic elements (MGEs); orange bar **Fig. 3B**) were overrepresented in the accessory genome of each species with the ratio of GCs in the accessory/persistent genome ranging from 4.2 (*C. tuberculostearicum*) to 36.1 (*C. pseudodiphtheriticum*). GCs assigned to “defense mechanisms” (purple bar **Fig. 3B**), which protect bacteria from MGEs, were more evenly distributed with the ratio of GCs in the accessory/persistent genome ranging from 1 (*C. tuberculostearicum*) to 2.9 (*C. pseudodiphtheriticum*). These findings are consistent with pangenomic analyses of other bacterial species, including our prior analysis of the candidate beneficial nasal bacterium *D. pigrum* (62).

**Figure 3.**
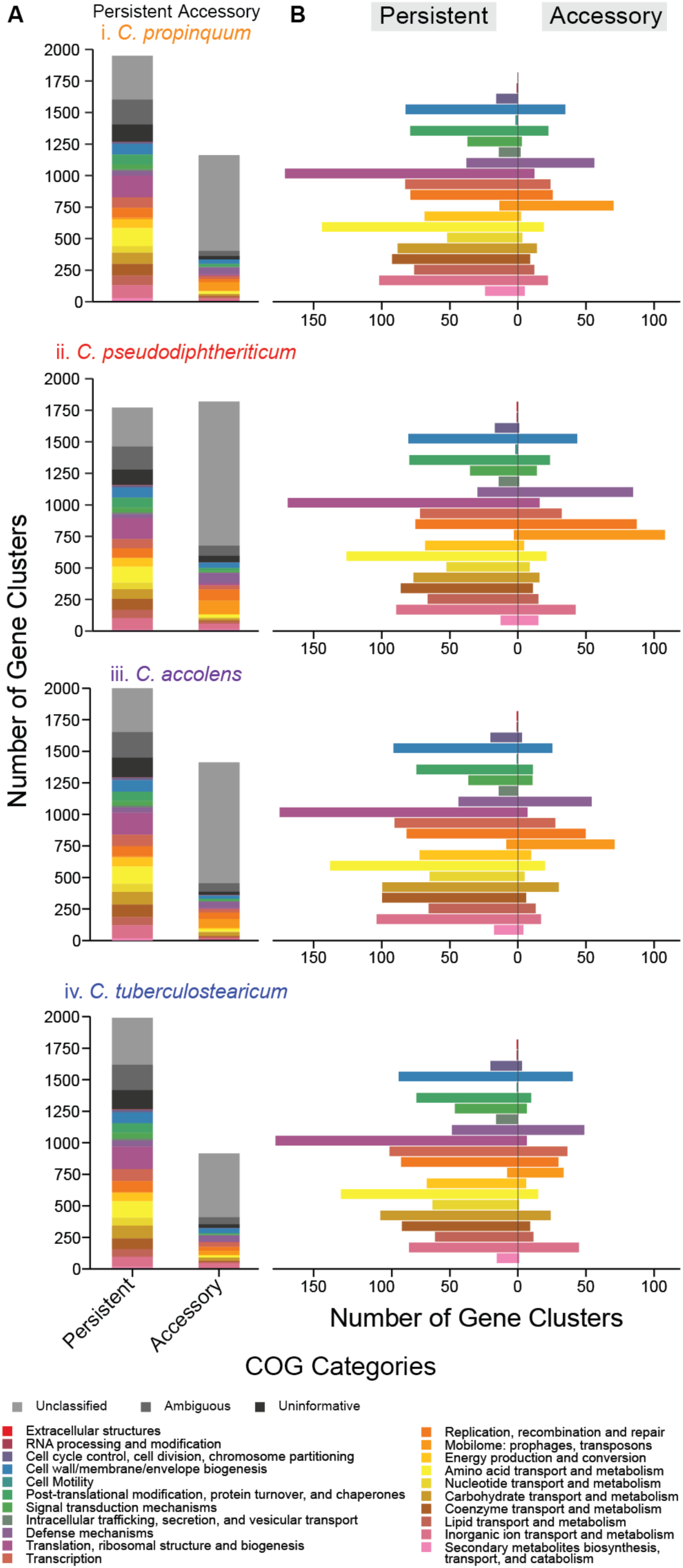
GCs assigned to COG metabolism categories are overrepresented in the persistent compared to the accessory genomes of each species indicating limited strain-specific metabolism. We identified the COG functional annotations for GCs using anvi’o and then used PPanGGOLiN to assign GCs to the persistent vs. accessory genome. **(A)** Over one-third of the GCs in each species (i-iv) were assigned as uninformative (black), ambiguous (dark gray), or unclassified (gray) across both the persistent and accessory genome. The combined percentage of each of these categories out of all the genes per species was 38.1% *Cpr*, 37.9% *Cps*, 37.1% *Cac*, and 38.3% *Ctu*. For each species, the percentage of GCs with an informative COG assignment was higher in the persistent genome, 64.9% *Cpr* (1262), 65.3% *Cps* (1156), 64.7% *Cac* (1300), and 63.5% *Ctu* (1264), than in the accessory genome, with 28.9% *Cpr* (336), 29.9% *Cps* (543), 25.7% *Cac* (363), and 35.6% Ctu (326). (B) Functional enrichment of GCs in the persistent vs. the accessory genome for the different COG categories. Metabolic COG categories, e.g., those involved in energy production (pale orange), or in amino acid (yellow), nucleotide (gold), carbohydrate (khaki), and lipid metabolism (dark salmon), were enriched in the persistent genome of each species. In contrast, mobilome (bright orange) and to a lesser extent defense mechanisms were enriched in the accessory genomes. Each *Corynebacterium* species shared similar COG functional enrichment ratios of GCs in its persistent vs. its accessory genome.

Our COG-enrichment analysis also showed that all the COG categories associated with metabolism, from “energy production and conversion” (pale orange) through “secondary metabolites biosynthesis, transport, and catabolism” (pink) in **Fig. 3B**, were highly overrepresented in the persistent (or core) genome of each species with ratios of accessory/persistent ranging from 0.02 to 0.56 (median of 0.16). The exception was an accessory/persistent GC ratio of 1.2 for “secondary metabolites biosynthesis, transport, and catabolism” in *C. pseudodiphtheriticum*. The overrepresentation of metabolic categories in the persistent genome of each species points to limited strain-level variation in metabolic capabilities, such as carbohydrate or amino acid metabolism. This contrasts with our previous analysis of *D. pigrum* in which GCs assigned to the COG category “carbohydrate transport and metabolism” are enriched in the accessory genome (ratio 1.66) (62).

### Common human nasal-associated *Corynebacterium* species have a largely shared metabolic capacity

Based on 16S rRNA V1-V3 sequences, 82% of adults are colonized with ≥ 2 of these 4 *Corynebacterium* species (see Tables S4A-B and S7 in (31)). This, combined with the enrichment of GCs assigned to metabolism COG categories in each persistent genome, led us to hypothesize that there would be much species-specific variation in core metabolic capabilities enabling the different nasal *Corynebacterium* species to occupy distinct metabolic niches within human nasal microbiota. To test our hypothesis, we used anvi’o to assign genes to Kyoto Encyclopedia of Genes and Genomes (KEGG) Orthology family (KO) annotations (**Table S3A**) and to estimate KEGG module completeness (**Table S4A-B**). In contrast to our hypothesis, we learned that *C. propinquum*, *C. pseudodiphtheriticum*, *C. accolens,* and *C. tuberculostearicum* encode highly conserved core metabolic capabilities sharing 43 of 58 (74%) detected fully complete KEGG modules (**Table 3**, **Fig. 4**). Next, we identified modules enriched in one or more of the four species, with various combinations of three of the four species sharing eleven additional modules (19%) and combinations of two of the four species sharing three additional modules (**Table S4D**). There were a few differences between the *C. propinquum-C. pseudodiphtheriticum* clade and the *C. accolens-C. tuberculostearicum* clade (**Fig. S1B**), with one and eight clade-specific KEGG modules, respectively. Only *C. tuberculostearicum*, which is broadly distributed across human skin sites as well as in the nasal passages (31, 34-37), was predicted to encode one complete KEGG module that was absent in the other three nasal species, as discussed in detail later.

**Figure 4.**
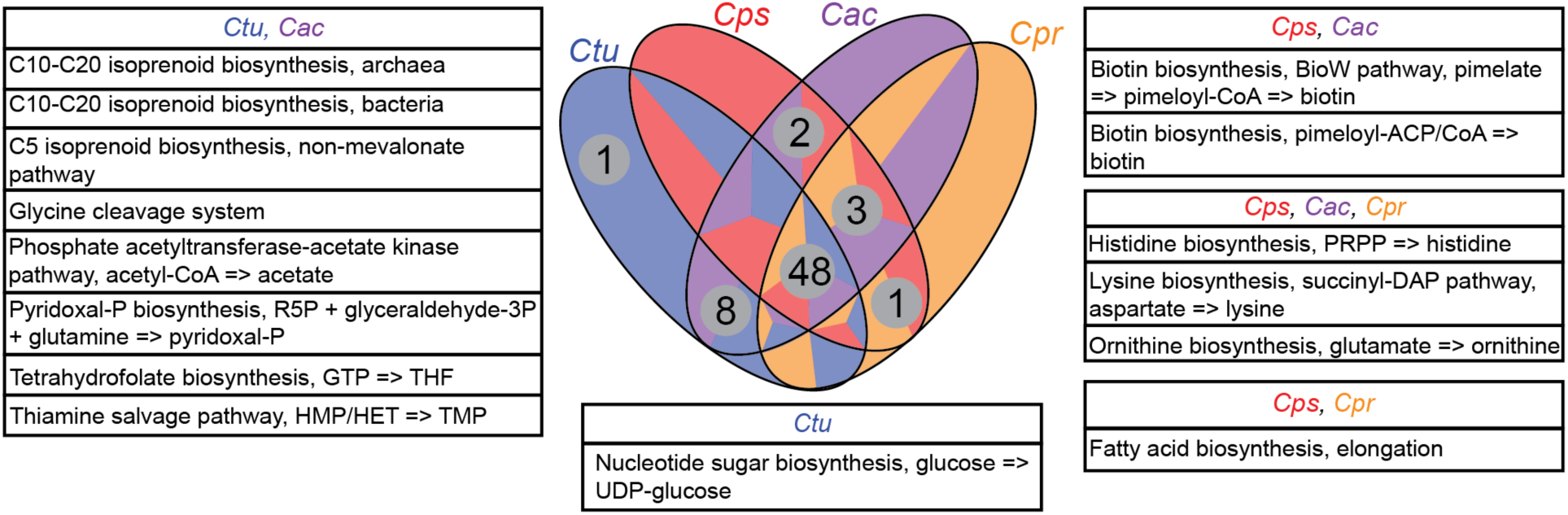
These four common human-nasal-associated *Corynebacterium* species have a largely shared metabolic capacity. The Venn diagram summarizes the results of an enrichment analysis of complete KEGG modules (stepwise completion score = 1) by species using anvi’o. Modules that both had an adjusted q-value > 1e-9 and were complete in at least 87% of the analyzed genomes were categorized as shared between the four *Corynebacterium*. Modules with an adjusted q-value < 1e-9 were considered enriched in their associated group of ≤ 3 species and are shown in boxes surrounding the Venn diagram. Species labels: *C*. *propinquum* (*Cpr*), *C. pseudodiphtheriticum (Cps), C. accolens (Cac)*, and *C. tuberculostearicum (Ctu*).

**Table 3.**
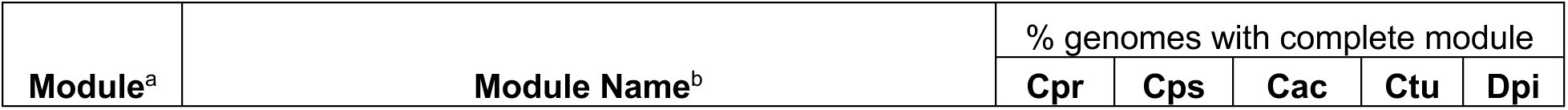
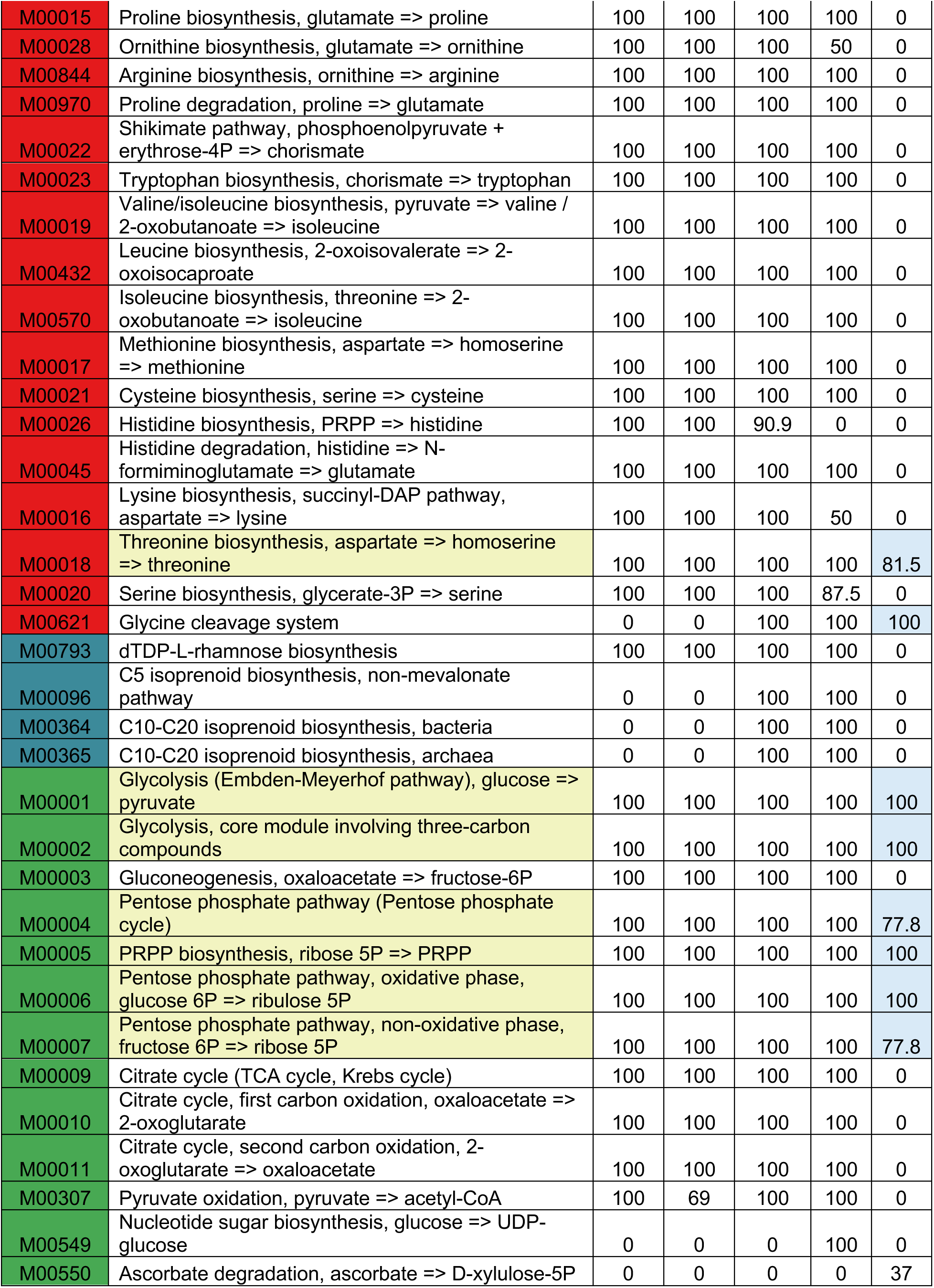
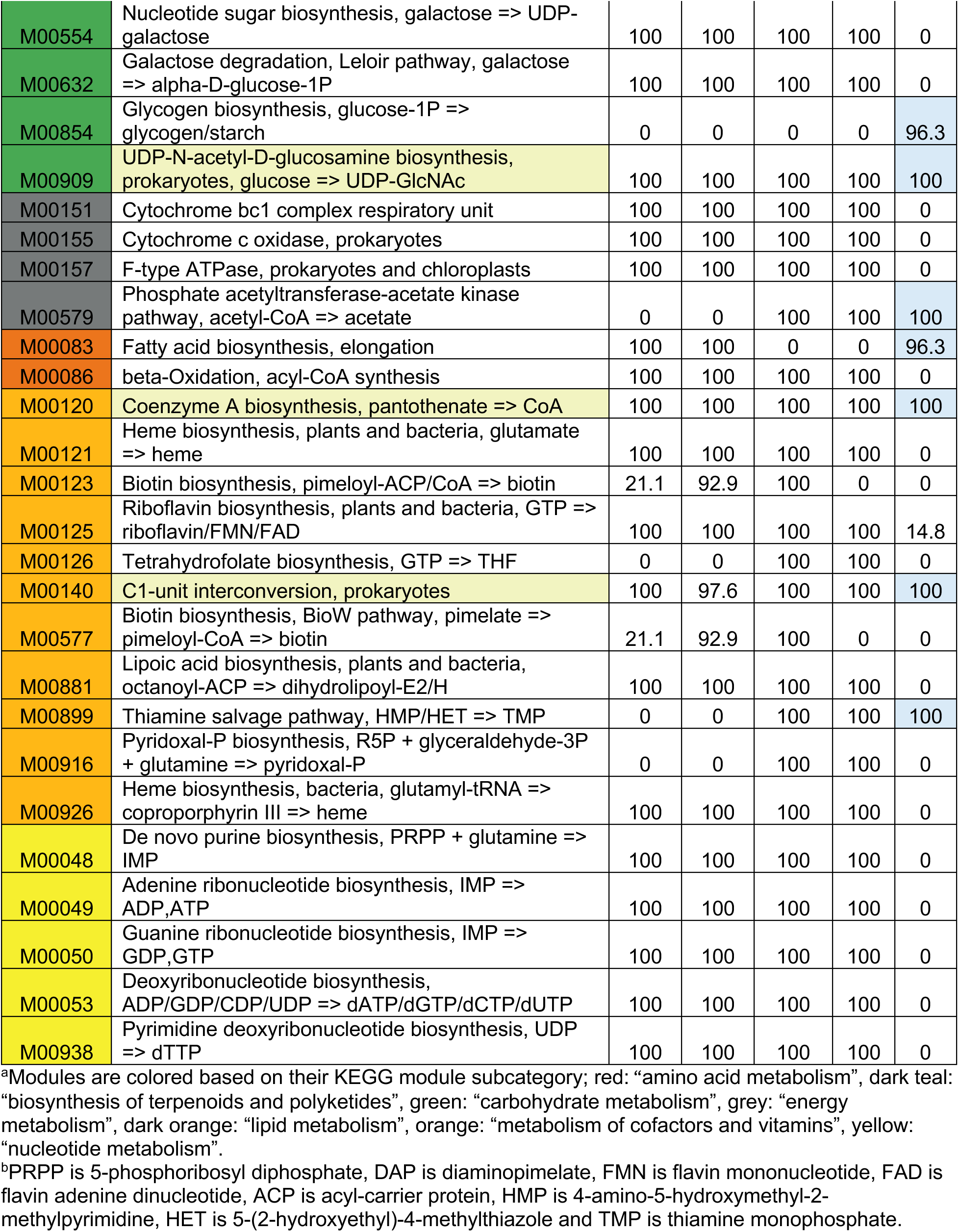
List of estimated complete KEGG modules.

### Nasal *Corynebacterium* species encode for central carbohydrate metabolism

To contextualize our findings within the genus *Corynebacterium*, we included in our anvi’o analyses the genomes of the type strains of two well-studied *Corynebacterium* species: *C. glutamicum* ATCC 13032 (*C. glutamicum*^T^) (68) and *C. diphtheriae* NCTC11397 (*C. diphtheriae*^T^) (**Table S1D**). The metabolism of the soil bacterium *C. glutamicum* is the best studied of *Corynebacterium* species (69). *C. diphtheriae* colonizes the human pharynx and toxigenic strains cause the human disease diphtheria (70). Along with *C. diphtheriae*^T^ and *C. glutamicum*^T^, the four human nasal *Corynebacterium* species all encoded complete modules for glycolysis and gluconeogenesis, the pentose phosphate pathway and phosphoribosyl pyrophosphate, and the tricarboxylic acid (TCA) cycle as part of central carbon metabolism (**Table 3**). In terms of other carbohydrate metabolism, all four nasal species, plus *C. diphtheriae*^T^ and *C. glutamicum*^T^, also encoded a complete module for UDP-N-acetyl-D-glucosamine biosynthesis (M00909), a precursor of cell wall peptidoglycan (71). All four species plus *C. diphtheriae*^T^, but not *C. glutamicum*^T^, also encoded the module for the Leloir pathway for galactose degradation.

### Nasal *Corynebacterium* species encode for synthesis of key biosynthetic cofactors, vitamins, and electron transport chain components

The strain genomes of all four species contained complete modules for biosynthesis of several cofactors and vitamins required for synthesis of essential biomolecules and central metabolism (**Table 3**). These include coenzyme A (M00120), required for the TCA cycle; lipoic acid (M00881), an organosulfur cofactor required in central metabolism (72); and C1-unit interconversion (M00140), which is connected to biosynthesis of tetrahydrofolate. Consistent with an intact TCA cycle, all four also had complete modules for the biosynthesis of key compounds involved in the electron transport chain, including heme (M00121, M00926), and riboflavin (M00125). *C. accolens* and *C. tuberculostearicum* also encoded biosynthesis of pyridoxal 5’-phosphate (M00916), which is a coenzyme in many transamination reactions (73); and tetrahydrofolate biosynthesis (M00126), which acts as a carrier for single carbon groups (**Table 3**). We also detected the modules for coenzyme A, lipoic acid, heme, riboflavin, and pyridoxal 5’-phosphate in both *C. glutamicum*^T^ and *C*. *diphtheriae*^T^. Of note, modules for the biosynthesis of cobalamin/vitamin B12 (M00925, M00924, M00122) were incomplete or absent in all four nasal *Corynebacterium* species. However, the nasal *Corynebacterium* species also encoded for the version of enzymes that are expected to be cobalamin independent, e.g., *metE* rather than the B12-dependent *metH*, so are unlikely to require cobalamin, consistent with predictions from Shelton and colleagues (74).

### Nasal *Corynebacterium* species share necessary modules for nucleotide synthesis and energy generation

All four of these common nasal *Corynebacterium* species had five complete modules related to nucleotide metabolism (yellow in **Table 3**) and three complete modules involved in ATP synthesis (M00151, M00155, M00157). Lastly, all four had complete modules for dTDP-L-rhamnose biosynthesis (M00793), a precursor to rhamnose cell wall polysaccharides. Rhamnose is part of the polysaccharide linker between peptidoglycan and arabinogalactan in members of *Mycobacteriales*, including *Corynebacterium* (71). Many of these 9 KEGG modules are also present in other common nasal microbionts with 8/9 in *Cutibacterium acnes* KPA171202 (75), 6/9 in *S. pneumoniae* TIGR4 (76) and *S. aureus* USA300_FPR3757 (77), 5/9 in *Staphylococcus epidermidis* RP62A (78), as well as 9/9 in *C. diphtheriae*^T^, and *C. glutamicum*^T^. However, all 9 of these modules were incomplete or absent across all 27 *D. pigrum* strains (62). (**Table S4C**).

### Nasal *Corynebacterium* species encode for biosynthesis of UDP-glucose via UDP-galactose

The UDP-glucose biosynthesis module (M00549) was fully detected in *C. tuberculostearicum* but was missing a step in the other nasal *Corynebacterium* and in *D. pigrum* (and present in all the other species analyzed (**Table S4C**)). UDP-glucose is a key part of central metabolism. It is the activated form of glucose that serves as a precursor for other activated carbohydrates and is used by most organisms for glucosyl transfers. Phosphoglucomutase (*pgm*) performs the second step in its three-step biosynthesis module. We identified a GC that included *pgm* from *C. glutamicum*, *C. tuberculostearicum,* and the skin-associated *Corynebacterium* species but lacked sequences from the other nasal *Corynebacterium* species. However, all four nasal *Corynebacterium* species encoded the module for UDP-galactose biosynthesis (M00554) plus a UDP-glucose 4-epimerase (K01784) to covert UDP-galactose to UDP-glucose. In contrast, the *D. pigrum* genomes encoded a *pgm* but lacked the third step, a UDP-sugar pyrophosphorylase or a UTP--glucose-1-phosphate uridylyltransferase (*galU*), for addition of glucose to UDP and also lacked the comparable step for addition of galactose to UDP suggesting the genes encoding for these steps are not yet included in the KEGG orthology.

### *C. tuberculostearicum* performs glycogen metabolism, unlike the other three nasal-associated *Corynebacterium* species

Based on anvi’o KEGG module reconstruction (**Table S4A-B**), three of the nasal *Corynebacterium* strains had a stepwise completeness score of zero for the modules for glycogen synthesis and degradation (M00854 and M00855), whereas *C. tuberculostearicum* had scores of 0.5 and 0.67, respectively, for these two modules. Likewise, *C. glutamicum*^T^ had completeness scores of 0.5 and 0.67 for these modules and was missing the same steps as *C. tuberculostearicum*. However, published data indicate *C. glutamicum* both synthesizes and degrades glycogen pointing to the use of glycogen for energy storage (79, 80). It encodes four genes for glycogen synthesis (*glgC*, *glgA*, *glgB, glgE*) and three genes for glycogen degradation (*glgX* and two copies of *glgP*). After identifying the predicted enzymatic function (KO number) that anvi’o assigned to the product of each of these six genes, we noted that all of these KOs are enriched in *C. tuberculostearicum* compared to the other three nasal *Corynebacterium* species (**Table S3B**). However, the KEGG module definitions for M00854 and M00855 lack the KOs corresponding to *glgA* (K16148), *glgE* (K16147), and *glgX* (K01214) in *C. glutamicum*^T^ and *C. tuberculostearicum*, and likely other *Corynebacterium* species. In fact, when we included these three KOs in the glycogen metabolism predictions for representative strains of other *Corynebacterium* species and non-*Corynebacterium* human nasal bacterial species, we identified complete glycogen synthesis pathways in *Corynebacterium diphtheriae* NCTC11397, the three common skin *Corynebacterium* species *Corynebacterium simulans* PES1 (81)*, Corynebacterium kroppenstedtii* DSM 44395 (82), and *Corynebacterium amycolatum* FDAARGOS_1108 (83) as well as in strains of other common nasal bacteria: *C. acnes* KPA171202 (75), *S. pneumoniae* TIGR4 (76), and *D. pigrum* (26/27 strain pangenome) (62). Glycogen degradation was complete in all of those strains except *C. kroppenstedtii* and *D. pigrum*, which were missing enzymatic activities for the first or second steps, respectively, of the module.

Based on this analysis, we predicted that *C. tuberculostearicum* can synthesize and degrade glycogen like *C. glutamicum*. To test for this, we measured intracellular concentrations of glycogen in the four nasal *Corynebacterium* species, using *C. glutamicum*^T^ as a positive control. We grew each in a liquid chemically defined medium (CDM) supplemented with all 20 amino acids and 5% glucose (**Fig. 5A**). We used a standardized estimated number of cells, based on OD_600_, to assay for intracellular glycogen. Under these conditions, *C. tuberculostearicum* harbored increased intracellular glycogen compared to the basal level observed in the other nasal strains (**Fig. 5B**).

**Figure 5.**
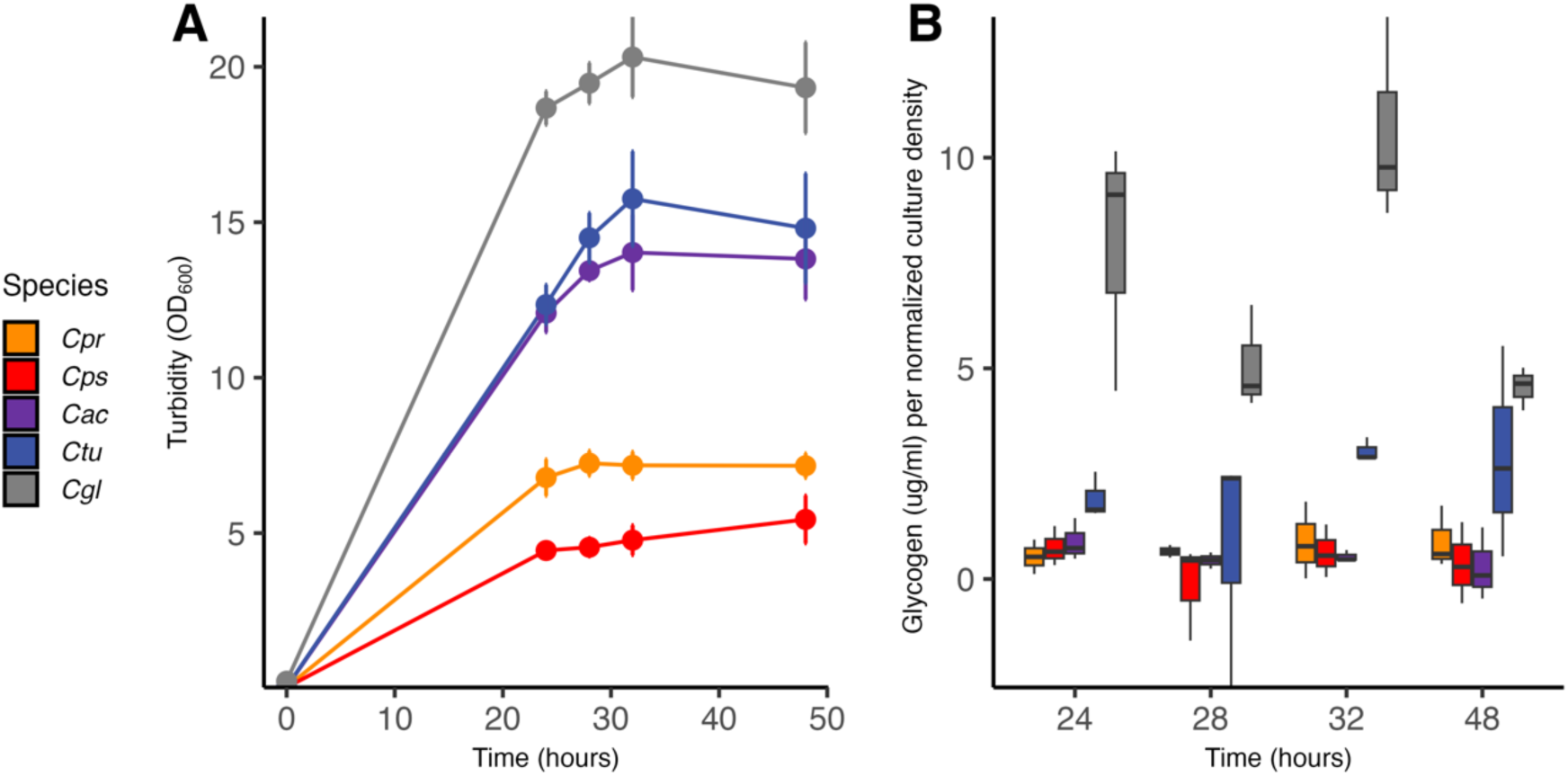
*C. tuberculostearicum* accumulates intracellular glycogen. (**A**) Growth curves for each of the nasal *Corynebacterium* species at 34°C in a MOPs-buffered CDM with all 20 amino acids and 5% glucose, with *C. glutamicum* as a positive control for the assay. *C. propinquum* and *C. pseudodiphtheriticum* reached stationary phase at a lower optical density at wavelength 600 nm (OD_600_) than did *C. tuberculostearicum* and *C. accolens*, with *C. glutamicum*^T^ reaching the highest. (**B**) Overall, *C. tuberculostearicum* accumulated more intracellular glycogen than the other three nasal species (*p*-value range 0.033 to 0.009), and less than *C. glutamicum* (*p* = 6.64 x 10^-10^). Intracellular glycogen content was quantified in µg/ml at timepoints between 24 and 48 hours. Statistics were done using a linear mixed model with species as a fixed effect and time point as a random effect. Boxplots show the median with first and third quartiles. Data are from *n* = 3 independent experiments. Species labels: *C. propinquum* (*Cpr*), *C. pseudodiphtheriticum (Cps), C. accolens (Cac), C. tuberculostearicum (Ctu*), and *C. glutamicum* (*Cgl*).

### Nasal *Corynebacterium* species can synthesize proteinogenic amino acids

Our overall analysis estimated that the vast majority of the analyzed strains of these four common human nasal *Corynebacterium* species can synthesize all 20 standard amino acids. In most of the genomes, we detected complete biosynthetic modules for 11 amino acids, including the hydrophobic amino acids (isoleucine, leucine, methionine, valine, and tryptophan); the polar uncharged amino acids (serine and threonine); the charged amino acids (arginine and lysine); cysteine; and proline (**Table 3**).

Based on anvi’o KEGG module reconstruction (**Table S4C**), the four nasal *Corynebacterium* species and *C. glutamicum*^T^ ATCC13032 have incomplete modules for phenylalanine (M00024) and tyrosine (M00025) due to lack of the corresponding aromatic aminotransferase KO. However, published experimental data confirms that the protein encoded by the *aroT* gene (NCgl0215 or CGL_RS01140) in *C. glutamicum*^T^ acts as an aromatic aminotransferase on substrates phenylpyruvate (O-Phe) and 4-hydroxyphenylpyruvate (O-Tyr) (84). Using anvi’o to cluster all of the genomes for the four nasal *Corynebacterium* species plus *C. glutamicum*^T^, we identified a GC that includes *C. glutamicum aroT* (annotated as K00817) and has matches in all of the analyzed strains. Based on this, we predicted that all four nasal *Corynebacterium* species should indeed be able to synthesize phenylalanine and tyrosine in the same way as *C. glutamicum*.

Substrate specificity of aminotransferase enzymes is notoriously difficult to predict from sequencing data due to the high degree of amino acid similarity within this enzyme family. Moreover, substrate promiscuity often leads to nonobservable phenotypes for deletion mutants of many aminotransferase genes (84-86). Both in *E.coli* (87) and *C. glutamicum* (84) this substrate overlap is especially significant for hydrophobic amino acids, including phenylalanine and tyrosine, which might explain why the initial anvi’o KEGG predictions were inaccurate for *aroT*. In fact, the identified *aroT* gene had an anvi’o KOfam annotation of histidinol-phosphate aminotransferase (K00817) and in *C. glutamicum*^T^ this motif is predicted in both *aroT* and *hisC* (named cg0267 and cg2304 respectively in (85)). In that study, *aroT* had higher similarity to a set of histidinol-phosphate aminotransferases with broader substrate specificity, whereas *hisC* was most similar to an aminotransferase specific for histidine biosynthesis. The *hisC* gene is also located in an operon with other genes involved in histidine biosynthesis. In our genomes of interest we also identified another GC including the *C. glutamicum*^T^ *hisC*.

The majority (91/102) of the genomes had a complete KEGG module for histidine biosynthesis (M00026), except that all *C. tuberculostearicum* and 3 *C. accolens* genomes were missing K01693. However, the gene cluster analysis revealed matches to this KO, corresponding to *C. glutamicum*^T^ *hisB,* in all 11 of these genomes, leading us to predict that all of the strains of interest can biosynthesize histidine.

### The *C. pseudodiphtheriticum* phylogeny revealed a recent and geographically localized loss of assimilatory sulfate reduction

In addition to the anabolic modules for biosynthesis, production of cysteine (M00021) and methionine (M00017) (**Table 3**) requires assimilatory sulfate reduction, which takes environmental sulfate and converts it to a usable form in the cell (88). Anvi’o analysis detected stepwise completion scores of only 0.5 for the assimilatory sulfate reduction module (M00176) for *C. glutamicum*^T^, all *C. accolens,* all *C. tuberculostearicum,* most *C. propinquum* (17/19), and 13 of 42 *C. pseudodiphtheriticum* genomes. However, *C. glutamicum*^T^ has proven assimilatory sulfate reduction capabilities via the *fpr2-cysIXHDNYZ* locus (89). This operon includes genes for two subunits of a sulphate adenylyltransferase, *cysD* (K00957) and *cysN* (K00956); an APS reductase, *cysH* (K00390); and a sulphite reductase, *cysI* (K00392). Of note, in *E. coli,* two enzymatic steps are required for the release of sulfite from APS: APS kinase (CysC) and PAPS reductase (CysH). Whereas, in *C. glutamicum*, *M. tuberculosis*, and *B. subtilis,* CysH is an APS reductase that directly converts APS to sulfite (89-91). However, the KEGG definition for M00176 currently fails to account for the experimental evidence that a complete assimilatory sulfate reduction module in *C. glutamicum*^T^ does not require an APS kinase. All the strains in our analysis with a stepwise completion score of 0.5 for this module have the same four KOs in the module as *C. glutamicum*^T^ and are, therefore, predicted to perform assimilatory sulfate reduction. However, this module was completely absent in all the *C. pseudodiphtheriticum* strains from the USA and 4 from Botswana (**Fig. 6A**), plus 2 *C. propinquum* strains, and *C. diphtheriae*^T^. The complete absence of the module for assimilatory sulfate reduction predicts that this subset of *C. pseudodiphtheriticum* strains cannot synthesize methionine or cysteine when sulfate is the only exogenous source of sulfur. To test this, we generated a chemically defined agarose medium supplemented with 18 amino acids (lacking cysteine and methionine) in which 75 mM sodium sulfate was the only exogenous source of sulfur. As predicted *C. pseudodiphtheriticum* MSK311, isolated in Botswana, showed growth under conditions requiring assimilatory sulfate reduction, whereas the USA isolates *C. pseudodiphtheriticum* KPL1989 and KPL4025 did not (**Figs. 6B** and **S5A**). These findings indicate a recent and geographically localized complete loss of the assimilatory sulfate reduction module M00176 within the *C. pseudodiphtheriticum* phylogeny.

**Figure 6.**
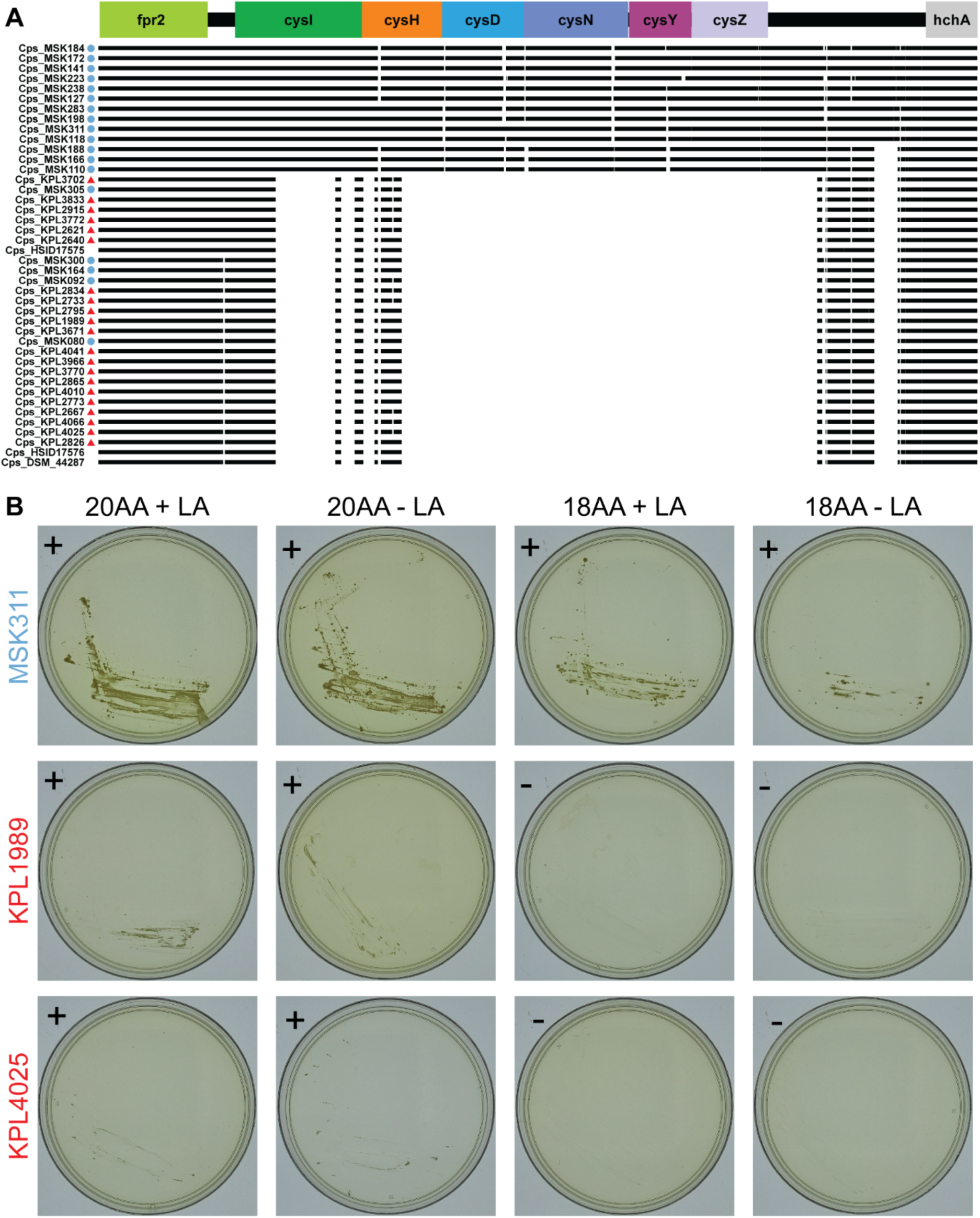
*C. pseudodiphtheriticum* strains from the Botswanan clade in Figure 1B can perform assimilatory sulfate reduction. (**A**) Most of the Botswanan *C. pseudodiphtheriticum* strains encode genes with KO annotations equivalent to those in the *C. glutamicum*^T^ *fpr2-cysIXHDNYZ* operon, which is required for assimilatory sulfate reduction. Figure represents the MAFFT multiple sequence alignment with strain genomes from Botswana (blue circles) and from USA (red triangles). (**B**) The Botswanan *C. pseudodiphtheriticum* strain MSK311, which encodes the *fpr2cysIXHDNYZ* operon, grew in chemically defined medium with sulfate as the only source of sulfur (KS-CDM 18AA - LA) indicating ability to assimilate sulfate, whereas the USA strains KPL1989 and KPL4025, which lack this operon, did not. Representative strains from the Botswanan (blue font) and USA (red font) clades in Figure 1B were grown on KS-CDM agarose with 2% glucose, supplemented with either 20 amino acids (20AA), including 2mM cysteine and 4mM methionine, or 18 amino acids (18AA), excluding cysteine and methionine; both with and without lipoic acid (LA, an additional source of sulfur). Images were captured after 8 days of growth at 34°C with 5% CO_2_ and humidification. Data are from four independent experiments with images from one experiment here and three others in Figure S5A. (+) = growth, (-) = no growth.

### Nasal *Corynebacterium* species encode for glycine and amino acids involved in nitrogen assimilation

KEGG lacks module definitions for biosynthesis of six of the standard amino acids: glycine, glutamate, glutamine, aspartate, asparagine, and alanine. Based on gene cluster analysis and the known capabilities of *C. glutamicum^T^*, we predicted that all four nasal *Corynebacterium* species can generate these six amino acids. In *C. glutamicum*^T^, the gene *glyA* encodes the enzyme serine hydroxymethyltransferase (SHMT) that generates glycine from L-serine (92). This essential gene corresponds with K00600, which was present in all the analyzed *Corynebacterium* genomes. Production of the remaining five amino acids is intertwined with nitrogen assimilation, which is well studied in *C. glutamicum*.

We predicted that the four nasal *Corynebacterium* species can assimilate ammonium and synthetize glutamate and glutamine using glutamate dehydrogenase and glutamine synthetase based on genomic comparisons to *C. glutamicum*^T^. Tesch et al. propose that glutamine amidotransferase reactions are critical for the flux of NH_4_^+^ into biomass in *C. glutamicum* based on^15^N-ammonium flux measurements showing that *C. glutamicum*^T^ assimilates 72% of the NH_4_^+^ into glutamate using glutamate dehydrogenase and 28% into glutamine via glutamine synthetase (93). Surprisingly, 2-oxoglutarate aminotransferase (GOGAT) does not actively contribute to NH_4_^+^ assimilation in *C. glutamicum*^T^ (93, 94). Moreover, *C. glutamicum*^T^ GOGAT mutations display only a slight increase in doubling time in minimal medium with limiting amounts of ammonium or urea as the nitrogen source, indicating GOGAT is nonessential for ammonium assimilation in *C. glutamicum*^T^ (94). In fact, the genes encoding for the two subunits of the GOGAT in *C. glutamicum*^T^ (*gltBD*) were absent in all of the nasal *Corynebacterium* strains, as were matches to the corresponding KOs (K00265 and K00266). In contrast, using GC analysis, we detected *C. glutamicum*^T^ *gdhA,* encoding glutamate dehydrogenase (K00262), and *glnA,* encoding glutamine synthetase, in all of the nasal strains (except for three *C. propinquum* strains that had a truncated *ghdA*), predicting that the four nasal *Corynebacterium* species can assimilate ammonium and synthetize glutamate and glutamine.

We predicted that the four nasal *Corynebacterium* species can all generate aspartate because *C. glutamicum aspT* (which encodes for an aspartate aminotransferase that interchangeably converts glutamate and oxaloacetate to 2-oxoglutarate and aspartate) (84), clustered with genes from all of the nasal *Corynebacterium* strains. The *C. glutamicum aspT* (aka *aspAT*) was recently recognized as part of the new subgroup 1c of class I aspartate aminotransferases (95), which likely accounts for the lack of KEGG annotation even though the COG and Pfam annotations are consistent with an aspartate aminotransferase domain (COG1167; PF12897.11).

We predicted that the four nasal *Corynebacterium* species do not require exogenous asparagine. We identified a GC including the *C. glutamicum*^T^ *asnB* gene plus genes from all of the analyzed *Corynebacterium* genomes annotated as an asparagine synthase (K01953). However, Hirasawa et al. report that the *C. glutamicum asnB* gene (aka *ltsA*, GenBank: AB029550.1) fails to complement an *E. coli asnA asnB* double mutant suggesting that it lacks asparagine synthetase activity, which led them to propose that it encodes for a glutamine-dependent amidotransferase that modifies cell wall component(s) involved in lysozyme and temperature sensitivity (96). The putative *asnB* genes of *Mycobacterium tuberculosis* and *Bacillus subtilis* are more similar to *C. glutamicum asnB* than to *E. coli asnB*, suggesting their gene products also lack an asparagine synthase function (96). This agrees with the observation that *M. tuberculosis* relies on the amidation of aspartate-tRNA to incorporate asparagine into proteins (97, 98). Based on these data and that *C. glutamicum^T^* grows in defined medium in the absence of amino acids, we predicted the four nasal *Corynebacterium* species also do not require asparagine.

We predicted that the four nasal *Corynebacterium* species can synthesize alanine since they all encoded *alaT* (K14260), an aminotransferase that converts pyruvate and glutamate to alanine and 2-oxoglutarate. *C. glutamicum*^T^ encodes two genes, *alaT* and *avtA*, that can both generate alanine via an amino transferase reaction, and mutants with a deletion of *alaT* have an L-alanine requirement under specific growth conditions (99).

To test our predictions, we assayed for growth on a base MOPS-buffered CDM agarose supplemented with 2% glucose with *C. glutamicum*^T^ as a positive control. All four nasal *Corynebacterium* species grew on MOPS-CDM agarose supplemented with all 20 proteogenic amino acids (**Figs. 7** and **S5B**). Along with *C. glutamicum*^T^, *C. accolens* and *C. tuberculostearicum* both grew in the absence of all 20 amino acids confirming the metabolic predictions, whereas *C. pseudodiphtheriticum* and *C. propinquum* did not. We hypothesized that MOPS-CDM lacking amino acids failed to support the growth in these two species due to nitrogen limitation. Therefore, we tested whether urea, glutamine (File S1), or asparagine, which are all easily bioavailable sources of nitrogen, would restore growth; however, none did so (**Figs. 7** and **S5B**). This leaves open the possibility that both *C. pseudodiphtheriticum* and *C. propinquum* are auxotrophic for at least one amino acid due to either a misannotation or a point mutation(s) resulting in a loss of function, or, alternatively, that under these specific growth conditions both fail to produce at least one of the required amino acids due to regulatory issues.

**Figure 7.**
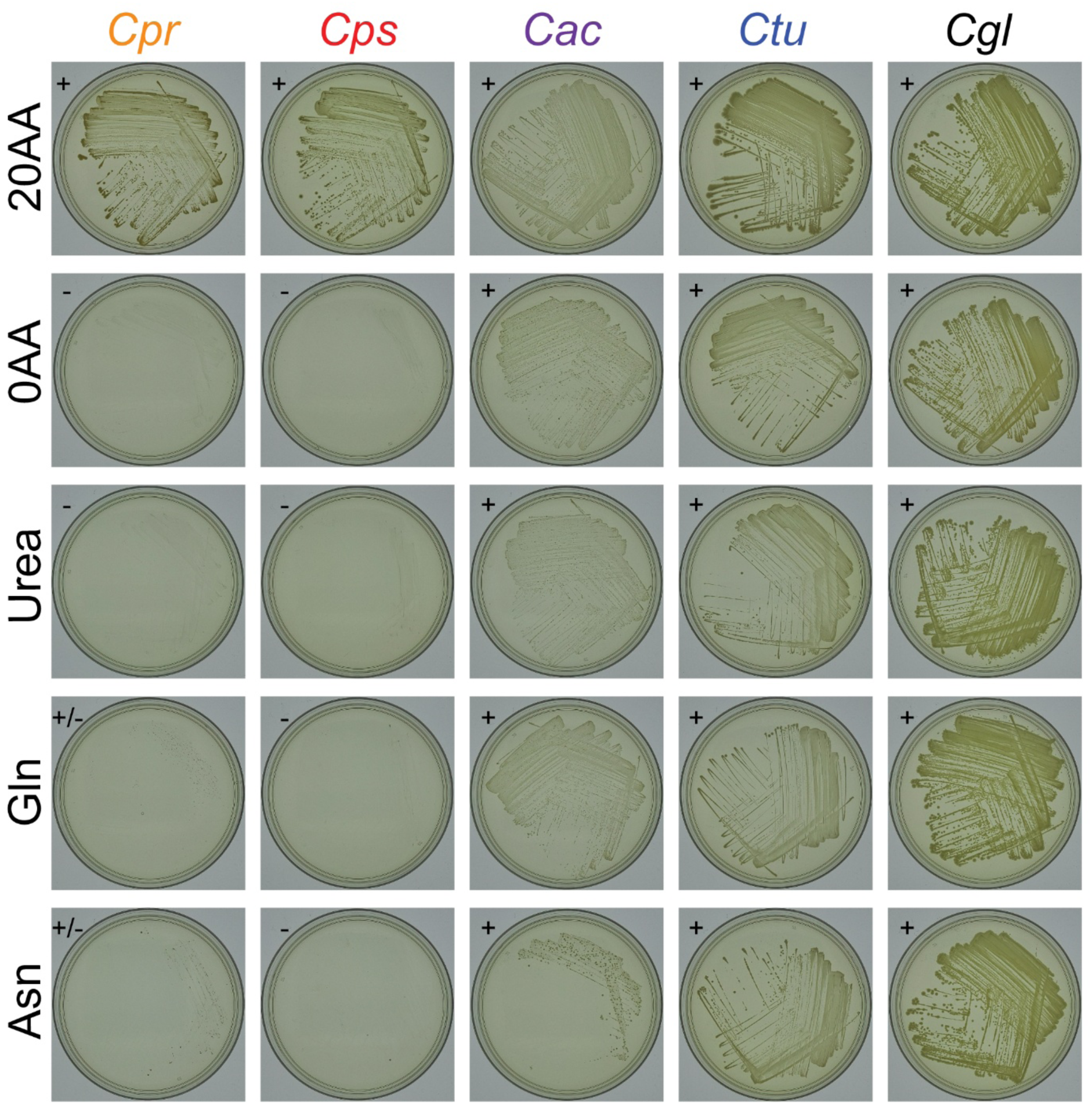
*C. accolens* and *C. tuberculostearicum* grow in the absence of all 20 amino acids on MOPS-buffered CDM agarose. Strains were grown on MOPS-buffered base CDM agarose with 2% glucose supplemented with 20 amino acids (20AA) vs. 0 amino acids (0 AA), and also in the absence of amino acids supplemented with either 5 mM Urea, 10 mM glutamine (Gln), or 10 mM asparagine (Asn) in *n* = 4 independent experiments. Shown are images from 1 experiment photographed after 8 days of growth at 34°C with 5% CO_2_ and humidification; remaining data are in Figure S5B. (+) = growth, (-) = no growth. Species labels: *C. propinquum* (*Cpr*), *C. pseudodiphtheriticum (Cps), C. accolens (Cac), C. tuberculostearicum (Ctu*), and *C. glutamicum* (*Cgl*).

### Human nasal *Corynebacterium* species have a broader metabolic capacity for biosynthesis of amino acids and cofactors/vitamins than *Dolosigranulum pigrum*

Many compositional studies of human nasal microbiota show a positive association at the genus level between *Corynebacterium* and *Dolosigranulum,* e.g., (1, 2, 4, 7, 24, 26, 38, 100). Nasal *Corynebacterium* species can enhance *in vitro* growth yields of *D. pigrum*, a lactic acid producing bacterium (38). Together with other prior analyses (101), these data indicate *D. pigrum* must access nutrients from its host and its microbial neighbors. As hypothesized, the nasal *Corynebacterium* species with their larger genome sizes (2.3 to 2.6 Mb) had a greater number of complete KEGG modules per genome than *D. pigrum* (1.9 Mb) (**Fig. 8**). Using anvi’o, we identified 15 complete modules shared by the majority (≥ 78%) of the 27 *D. pigrum* strain genomes (**Table 3**, highlighted in light blue). These are for the metabolism of amino acids (2), carbohydrates (8), energy (1), lipids (1), and cofactors and vitamins (3). This is approximately 30% of the number of complete modules found in the majority of each of the four *Corynebacterium* species’ genomes (range 47 to 56). The module for glycogen biosynthesis (M00854) was the only one found in *D. pigrum* and incomplete in all four *Corynebacterium* species (**Table 3**); however, as described above, *C. tuberculostearicum* produces glycogen (**Fig. 5**). Of note, each *D. pigrum* genome had at least one CDS annotated as a sialidase (K01186), which can release sialic acid from sialylated glycans found in mucus providing bacteria with carbon and nitrogen (102), whereas there were none in the nasal *Corynebacterium* genomes (**Table S3A**). Only 10 complete KEGG modules were shared by *D. pigrum* and all 4 *Corynebacterium* species (**Table 3**, highlighted in pale yellow). In contrast, the 4 *Corynebacterium* species shared 12 modules for amino acid metabolism and 4 modules for cofactor/vitamin metabolism in the majority of their genomes that were absent/incomplete in *D. pigrum* (**Table 3**).

**Figure 8.**
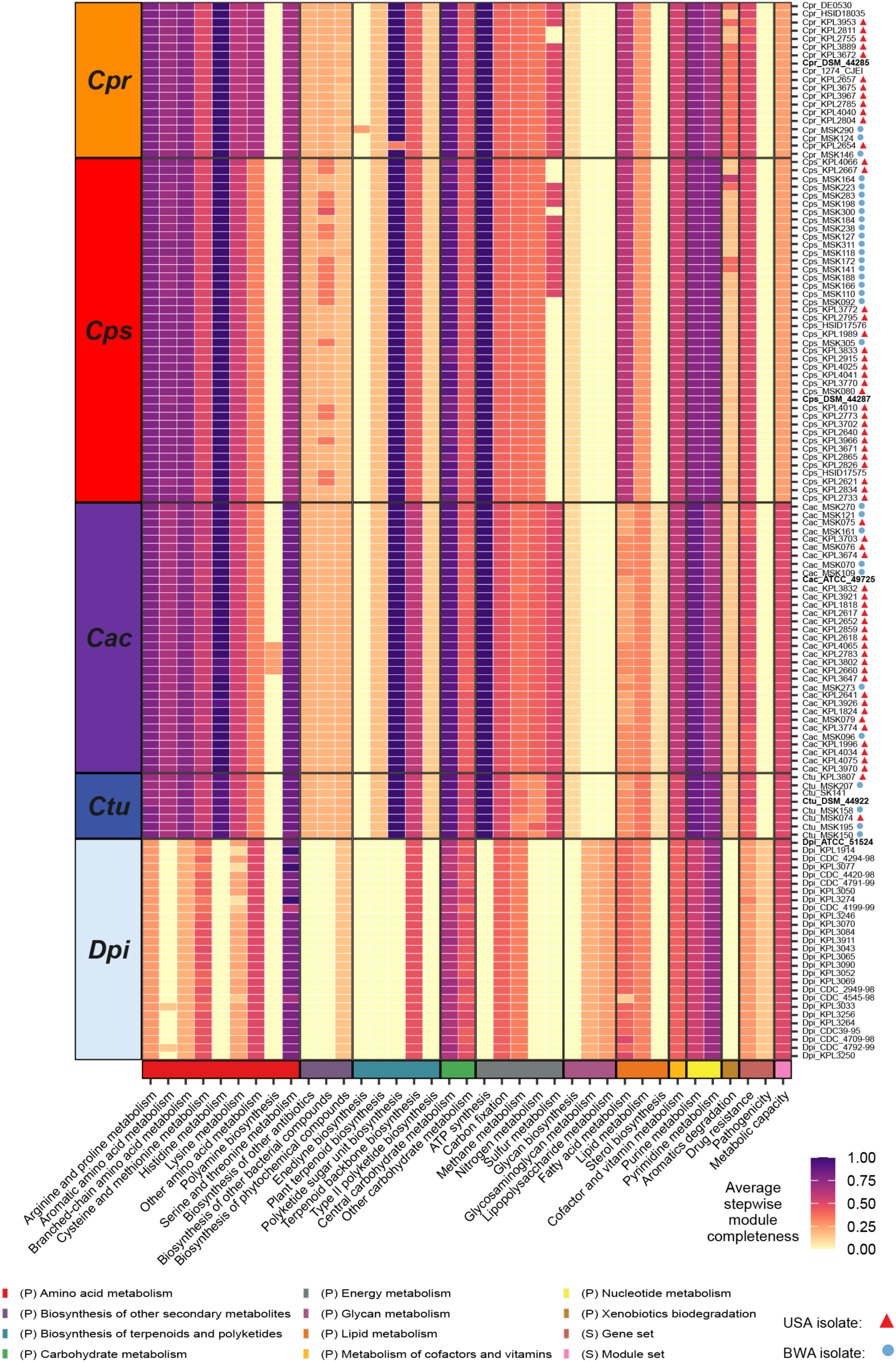
Each of the four nasal *Corynebacterium* species encode for an increased metabolic capacity compared to *D. pigrum*. This heatmap shows stepwise module completeness averaged by KEGG module subcategory (y axis labels) for each of four nasal *Corynebacterium* species (avg. genome size 2.3 - 2.5 Mb) and for *D. pigrum* (avg. genome size 1.9 Mb). Average stepwise completion scores were calculated including only modules detected in at least one of the analyzed genomes. Color legend on the bottom represents KEGG module categories and corresponds with the colors in Table 3. (P) and (S) indicate Pathway and Signature modules respectively. Species labels: *C. propinquum* (*Cpr*), *C. pseudodiphtheriticum (Cps), C. accolens (Cac), C. tuberculostearicum (Ctu*), and *D. pigrum* (*Dpi*).

## DISCUSSION

Here, we analyzed strain genomes of four common *Corynebacterium* species including those of 87 distinct human nasal isolates collected in Africa and North America across the human lifespan. Phylogenomic analysis showed *C. pseudodiphtheriticum* displays geographically restricted strain circulation. This corresponded with a recent geographically restricted loss of the KEGG module for assimilatory sulfate reduction in strains isolated in the USA, since this module was present in the other three species and in most *C. pseudodiphtheriticum* strains from Botswana. We confirmed the absence of the *cysIXHDNYZ* operon in the USA *C. pseudodiphtheriticum* strains and experimentally demonstrated that representative USA strains failed to grow under conditions requiring assimilatory sulfate reduction, whereas a strain from Botswana grew (**Fig. 6**). Across the four species, genomic analysis revealed average genome sizes of 2.3 to 2.5 Mb, with the average CDS per genome ranging from 2105 to 2265 and with 72-79% of each individual genome encoding GCs of the shared conservative core genome of the respective species. For each species, the core genome size had leveled off while the pangenome remained open. An informative assignment to a definitive COG category was possible only for approximately 65% of the GCs in the persistent genome and 26-36% of the GCs in the accessory genome of each species, which points to the need for ongoing experimental research to identify the function of many bacterial GCs. GCs assigned to the COG categories for metabolism were overrepresented in the persistent genome of each species and all four species shared the majority (43 of 58) of complete KEGG modules identified, which implies limited strain- and species-level metabolic variation restricting the possibilities for strains to occupy distinct metabolic niches during the common occurrence of cocolonization of the human nasal passages. We experimentally validated the genome-based metabolic prediction that of the four species only *C. tuberculoste*a*ricum* accumulates intracellular glycogen (**Fig. 5**). *Corynebacterium* species are often positively associated with *Dolosigranulum* in human nasal microbiota. We found human nasal *Corynebacterium* species have a broader metabolic capacity for biosynthesis of amino acids and cofactors/vitamins than *D. pigrum,* supporting the possibility that *Corynebacterium* might cross-feed or serve as a source of nutrients for *D. pigrum,* and possibly other microbionts, in human nasal microbiota. We experimentally validated that *C. accolens* and *C. tuberculostearicum* both grow on defined medium lacking amino acids. Our findings combined with data showing that the majority of adults likely host at least two *Corynebacterium* species in their nasal passages points to the importance of future investigation into how *Corynebacterium* species interact with each other and with other microbionts in human nasal microbiota.

By analyzing the extent and distribution of cobamide production in 11,000 bacterial species, Shelton et al. report 86% of bacteria in their dataset have at least 1 of the 15 cobamide-dependent enzyme families, but only 37% are predicted to synthesize their own cobamide (74) pointing to widespread interspecies cobamide sharing among bacteria. Swaney et al. find d*e novo* cobamide biosynthesis is enriched in host-associated compared to environment-associated *Corynebacterium* species, with several human skin-associated *Corynebacterium* species encoding complete biosynthesis pathways (61). In contrast, all four nasal *Corynebacterium* species had nearly absent modules for production of the corrin ring (M00925 or M00924) and the nucleotide loop (M00122) (**Table S4A-B**). They also lacked cobamide-dependent enzymes. Furthermore, based on Shelton and colleagues’ analysis of representative genomes, the majority of the most common human nasal bacteria are both likely nonproducers and lack B12-dependent enzymes and B12 pfam binding domains (74), the exception being *C. acnes*, which is a highly prevalent and abundant member of human nasal microbiota, especially in adults (31), and produces the cobamide vitamin B12 (cobalamin) (103).

Limitations of this study include the uneven representation of strains from the USA and Botswana for *C. propinquum* and *C. accolens*; the limited number of *C. tuberculostearicum* strains; the inherent limitations of KEGG annotations; and the predictive nature of genome-based metabolic estimations, some of which still require future experimental validation. To our knowledge, this analysis includes the largest number of strain genomes for *C. pseudodiphtheriticum, C. propinquum*, and *C. accolens* to date, with the aforementioned smaller number for *C. tuberculostearicum*. However, compared to the thousands of strain genomes that have been analyzed for nasal pathobionts, e.g., *S. aureus* (104), there are still a limited number of available genomes of nasal *Corynebacterium* species. This highlights the need to build large strain collections of human-associated *Corynebacterium* species to better assess the potential use of these strains for the promotion of human health. Similarly, although we included genomes for strains isolated from two continents and from a range of ages, the geographic sampling was limited compared to the distribution of human populations globally and there has yet to be a systematic large-scale sampling of nasal microbiota across the human lifespan.

Qualitatively, we isolated fewer *C. tuberculostearicum* from nasal swabs than expected based on its prevalence and relative abundance estimated in our earlier reanalysis of 16S rRNA gene V1-V3 sequences from human nasal samples (31). In contrast to using a single gene, here, we assigned isolates to *C. tuberculostearicum* based on WGS with an ANI of ≥ 95% to the type strain *C. tuberculostearicum* DSM 44922. Only a subset of our isolates from the USA and Botswana with partial 16S rRNA gene Sanger sequences (approximately V1-V3) matching to *C. tuberculostearicum* met this criterion after WGS. This points to the existence of another common nasal *Corynebacterium* species that is closely related to *C. tuberculostearicum.* Recent human skin metagenomic analyses by Salamzade et al. identify metagenome-assembled genomes and the strain genome LK1134 with ANI ≥ 95% to the genome called “*Corynebacterium kefirresidentii”* (105), which is not a validly published species, and show via phylogenomic analysis these are closely related to *C. tuberculostearicum* (34). Furthermore, using metagenomic analyses, they show sequences matching the “*C. kefirresidentii”* genome are more prevalent on nasal and nearby facial skin sites, whereas *C. tuberculostearicum* is more prevalent and at higher relative abundance on foot-associated skin sites (34). Future work to validly name the species currently identified by the genome called “*C. kefirresidentii*” with designation and deposition of a type strain in publicly accessible stock centers is critical for experiments seeking to identify the function of this species in human nasal microbiota. Isolation and whole genome sequencing of multiple strains for microbial species commonly detected in the human microbiome, such as this one, is an ongoing effort across multiple body sites. Collections of genome-sequenced strains from the microbiota are a critical resource for experimentally testing hypotheses generated from metagenomic and metatranscriptomic studies to identify the functions of human microbionts and mechanisms by which they persist in the microbiome. The Human Oral Microbiome Database (<u>eHOMD</u>) is an early and ongoing example of a body-site focused resource for the human microbiome based on a combination of culture-dependent and -independent data (106). Originally focused on the oral cavity, eHOMD now serves the full human respiratory tract (31). More recently, Saheb Kashaf et al. established the Skin Microbial Genome Collection (SMGC), a combined cultivation- and metagenomic-based resource for the human skin microbiome (107). These well-curated, body-site focused databases serve a critical role in advancing microbiome research, including their importance in shedding light on so-called microbial and metagenomic “dark matter.” The data we presented here serves as a foundational resource for future genomic, metagenomic, phenotypic, metabolic, functional, and mechanistic research on the role of nasal *Corynebacterium* species in human development and health.

## MATERIALS AND METHODS

### Collecting new nasal *Corynebacterium* sp. isolates

The USA *Corynebacterium* strains with KPL in their name were collected in Massachusetts, USA under a protocol approved by the Forsyth Institutional Review Board (FIRB #17-02) as described previously (62). In brief, adults and children participating in scientific outreach events in April 2017 and 2018 performed supervised self-sampling of their nostrils (nasal vestibule) with sterile swabs. They then inoculated their swab onto up to two types of agar medium: 1) brain heart infusion with 1% Tween80 (BHIT) and 25 microgram/ml fosfomycin (BHITF25) to enrich for *Corynebacterium* sp. and/or 2) BBL Columbia Colistin-Nalidixic acid agar with 5% sheep’s blood (CNA BAP). These were incubated at 37°C for 48 hrs under either atmospheric (BHITF25) or 5% CO_2_-enriched (CNA BAP) conditions. We selected colonies with a morphology typical of nasal *Corynebacterium* species and passed each two to three times for purification on BHIT with 100 ug/ml fosfomycin (BHITF100) at 37°C prior to storage in medium with 15-20% glycerol at -80°C. (Isolates from 2017 were picked from growth on BHITF100 at 37°C under atmospheric conditions that had been inoculated from sweeps of the original mixed growth on agar medium and stored at -80°C.)

The majority of the *Corynebacterium* strains with MSK in their name were cultured from nasopharyngeal swab samples collected from mothers and infants in a birth cohort study conducted in Botswana, as previously described (13), with a small number also collected from mid-turbinate nasal swab samples from patients cared for within the Duke University Health System (MSK074, MSK075, MSK076, MSK079, and MSK080). This work was reviewed and considered exempt by the Duke Health Institutional Review Board (Pro00102629). Bacteria were cultivated and isolated as previously described (13).

### Selection of nasal *Corynebacterium* isolates for Illumina sequencing

For each KPL-named new isolate, Sanger sequencing (Macrogen, USA) was performed using primer 27F on a V1-V3 16S rRNA gene colony-PCR amplicon (GoTaq Green, Promega) of primers 27F and 519R. We assigned each initial isolate to a genus and a putative species based on blastn of each sequence against eHOMDv15.1 (31). We then selected a subset of these isolates for whole genome sequencing (WGS). For MSK-named new isolates, all isolates preliminarily assigned to *Corynebacterium* based on MALDI and/or Sanger sequencing of V1-V3 16S rRNA gene underwent WGS.

### Genomic DNA extraction

We extracted genomic DNA (gDNA) from the KPL-named USA strains using the MasterPure Gram Positive Genomic DNA Extraction Kit with the following modifications to the manufacturer’s protocol: 10 mg/mL lysozyme treatment at 37°C for 10 min and 2x 30 sec bead beat in Lysing Matrix B tubes (MP Biomedicals) at setting 6 on a FastPrep24 (MP Bio) with 1-minute interval on ice. To assess gDNA quality, we performed electrophoresis on 0.5% TBE agarose gel, used a NanoDrop spectrophotometer to quantify 260/280 and 260/230 ratios, and used a Qubit Fluorometric Quantification (Invitrogen) to measure concentration. We extracted gDNA from the MSK-named strains collected in Botswana and North Carolina using the Powersoil Pro extraction kit (Qiagen) following the manufacturer’s instructions. DNA concentrations were determined using Qubit dsDNA high-sensitivity assay kits (Thermo Fisher Scientific).

### Whole genome sequencing and assembly

For the KPL-named USA isolates, Nextera XT (Illumina) libraries were generated from gDNA. Each isolate was sequenced using a paired-end 151-base dual index run on an Illumina Novaseq6000 at the NIH Intramural Sequencing Center. The reads were subsampled to achieve 80x coverage and then assembled with SPAdes (version 3.13.0) (108) and polished using Pilon (version 1.22) (109). For the MSK-named isolates, which are mostly from Botswana, library preparation was performed using DNA Prep Kits (Illumina) and these libraries were sequenced on a NovaSeq 6000 instrument (Illumina) configured for 150 base pair paired-end reads. Adapter removal and read trimming were performed using Trimmomatic version 0.39 (110) to a Phred score of 30 across a 4-bp sliding window. Surviving reads shorter than 70 bp were discarded. The final quality of reads was assessed using FastQC version 0.11.9. Assembly was performed using SPAdes version 3.15.3 (111). The completeness of the genomes was evaluated with checkM version 1.1.3 (112) and all genomes with a completeness less than 95% were discarded. Genomic data are deposited under BioProjects PRJNA842433 for the KPL-named isolates (which are a subset of 94 *Corynebacterium* isolated in MA, USA) and PRJNA804245 for the MSK-named isolates (which are a subset of 71 genomes isolated from Botswana and the Duke University Health System). **Table S1A** includes NCBI accession IDs.

### Selection of strain genomes for pangenomic analysis

To the 165 assemblies mentioned in the previous section, we added another 16 KPL-named *Corynebacterium* sp. nasal-isolate genomes originally sequenced as part of the HMP and deposited by the Broad at NCBI to consider for analysis (113). Furthermore, 31 reference assemblies for relevant *Corynebacterium* species, including the genome of the type strain of *C. propinquum*, *C. pseudodiphtheriticum*, *C. accolens,* and *C. tuberculostearicum* plus 3 strain genomes of *C. macginleyi,* were downloaded from NCBI using the PanACoTA v1.4.1 (114) ‵prepare -s‵ subcommand. We used default parameters such that genomes with MASH distances to the type strain outside of the range 1e-4 to 0.06 were discarded to avoid redundant pairs or mislabeled assemblies and low-quality assemblies based on L90 ≤ 100 and number of contigs ≤ 999 were filtered out. The collected 212 assemblies were filtered using the ‵prepare --norefseq‵ subcommand as above to select higher quality assemblies (L90 ≤ 100 and number of contigs ≤ 999) and to eliminate redundant genomes defined by a MASH distance < 10^-4^ keeping the genome with the highest quality score from each redundant set. Finally, we confirmed the species-level assignment of our nasal isolates, and the nontype reference strains, based on an ANIb (nucleotide) of ≥ 95% for all shared CDS regions compared to the respective type strain of each species using GET_HOMOLOGUES (see below). For each species, this resulted in a set of distinct strain genomes (including the type strain) that we used for subsequent analyses, which totaled to 104 genomes: 19 *C. propinquum* genomes, 43 *C. pseudodiphtheriticum* genomes, 34 *C. accolens* genomes, and 8 *C. tuberculostearicum* genomes. **Table S1A** contains a list of these 104 strain genomes selected for further analysis plus 3 *C. macginleyi* reference strains, and **Table S1B** the all-by-all MASH distance analysis result of the PanACoTA analysis for all 107 genomes.

### Determination of the conservative core genome

We annotated all bacterial genomes with Prokka version 1.14.6 (115) with default parameters, including gene recognition and translation initiation site identification using Prodigal (116). Then, we used the ‵./get_homologues.pl‵ command from GET_HOMOLOGUES version 24082022 (117, 118) to determine a conservative core genome for the selected *Corynebacterium* genomes based on the consensus of three algorithms: bidirectional best-hits (BDBH), cluster of orthologs triangles (COGS) v2.1 (119), and Markov Cluster Algorithm OrthoMCL (OMCL) v2.4 (120) (**Figs. S1A** and **S2C**). Each of the three algorithms reported clustering at the protein level using blastp from NCBI BLAST v2.2 (121) with ‵-C 90‵ (min % coverage in BLAST pairwise alignments). The data output created from running the three different clustering algorithms was used to identify the intersection of the core GCs with the command ‵./compare_clusters.pl‵ with ‵-t # of genomes‵. We ran this last command twice, with and without the -n flag to generate both nucleotide and protein outputs. Additional methods for genome annotations (https://klemonlab.github.io/CorPGA_Pangenomics/SupplementalMethods_Prokka_Annotations.html) and for the determination of the conservative core genome are available online (https://klemonlab.github.io/CorPGA_Pangenomics/SupplementalMethods_GET_HOMOLOGUES.html).

### Determination of core, soft core, shell, and cloud genomes

For this analysis to identify the core genomes, we dropped the strains Cps_090104 and Cac_ATCC_49726 from their respective species because, although these both passed CheckM analysis, their inclusion in the initial rarefaction analyses caused a splitting of data points towards the lower end resulting in an aberrant lower bound. We separately analyzed the genomes of each *Corynebacterium* species using GET_HOMOLOGUES with the command ‵./get_homologues.pl‵ and flag‵-t 0‵ (to get all the possible clusters), first with ‵-M‵ (OrthoMCL) and a second time with ‵-G‵ (COGS). The COGS and OMCL results were then used with ‵./compare_clusters.pl‵ with ‵-m‵ (produce intersection pangenome matrices). Then the command ‵./parse_pangenome_matrix.pl‵ was used with ‵-s‵ (report core, soft core, shell, and cloud clusters), and ‵-x‵ (produce matrix of intersection pangenome clusters) (**Fig. S3Ai-iv**). In-depth code and descriptions to determine the core, soft core, shell, and cloud genomes are available online (https://klemonlab.github.io/CorPGA_Pangenomics/SupplementalMethods_GET_HOMOLOGUES.html).

### Calculating the conservative core genome for each species with the addition of an outgroup

We chose the type strain genome of the most closely related species from the phylogenomic tree in **Fig. S1B** to serve as the outgroup in each species-specific phylogeny with *C. pseudodiphtheriticum* DSM 44287 for the *C*. *propinquum* phylogeny, *C. propinquum* DSM 44285 for the *C. pseudodiphtheriticum* phylogeny*, C. macginleyi* CCUG 32361 for the C. *accolens* phylogeny, and *C. accolens* ATCC 49725 for the *C. tuberculostearicum* phylogeny. Addition of an outgroup resulted in four new datasets with n + 1 total genomes, where n denotes the number of strain genomes for each species. Each species-plus-outgroup dataset of genomes was analyzed as before using the commands ‵. /get_homologues.pl‵ (with BDBH, COG triangles, and OrthoMCL) and ‵./compare_clusters.pl‵, but now with the outgroup genome included (**Fig. S2D**). For construction of single-species trees (**Fig. 1**), we then combined this smaller conservative core of GCs shared between the species and its outgroup (shared intersection of each Venn diagram in Fig. S2D) with the subset of GCs belonging only to the conservative core of each species (which is the intersection of each Venn diagram in Fig. S2C minus the intersection of those in Fig. S2D). The code used to do this is available online (https://klemonlab.github.io/CorPGA_Pangenomics/SupplementalMethods_GET_HOMOLOGUES.html).

### Construction of phylogenomic trees

We used GET_PHYLOMARKERS v.2.2.9.1 to concatenate and align the single copy core GCs for each phylogeny (122). The command ‵run_get_phylomarkers_pipeline.sh‵ was run with the flags: ‵-R 1 -t DNA -k 0.7 -m 0.7‵ on both the protein and nucleotide GET_HOMOLOGUES outputs. The flag ‵-R‵ was used to select optimal markers for phylogenomics; ‵-t‵ for whether the input is DNA or protein; ‵-k‵ for kde stringency; and ‵-m‵ for the minimum average support values for trees to be selected. The codon fasta alignments generated by GET_PHYLOMARKERS were analyzed with IQ-TREE v2.1.3 (123) with: ‵-p‵, (uses edge linked partition model and ModelFinder functions (124-126)), ‵-alrt 1000‵ (perform replicate SH-like approximate likelihood ratio test) and ‵-B 1000‵ (number of ultrafast bootstrap replicates). The phylogenetic tool iTOL v6 (127) was used to visualize, scale, edit, annotate names, and root the tree at the midpoint for each phylogeny. Detailed code and methods to create these phylogenies are available online (https://klemonlab.github.io/CorPGA_Pangenomics/SupplementalMethods_GET_HOMOLOGUES.html).

### Rarefaction analysis and Average Nucleotide Identity (ANI) across strain genomes

For each species, with GET_HOMOLOGUES, we used rarefaction analysis to estimate whether the core genome and pangenome were closed or open (**Fig. 2**). We modified “$MIN_PERSEQID_HOM” and “$MIN_COVERAGE_HOM” values to equal 50 (inside “marfil_homology.pm”) so only protein sequences with identity and coverage ≥ 50% will be called homologues.

Again, we made the decision to eliminate the two genomes, namely Cac_ATCC_49726 and Cps_090104 from this dataset. The reason for this action was that their core genome size resulted in the preliminary graph getting divided while conducting random sampling. This is because their core genome sizes were considerably lower, resulting in a misleading lower bound. The clustering was redone for each species using ‵./get_homologues.pl‵ and the following parameters: ‵-C 90‵, ‵-c‵, (genome composition analysis) and ‵-M‵ for OMCL. The rarefaction curve .tab files were produced from the ‵-c‵ flag. Rarefaction .tab files for each species were plotted into svg files using ‵/plot_pancore_matrix.pl‵ with flags ‵-f core_both‵ (displays both Tettelin and Willenbrock curves for core genome), and ‵-f pan‵ (curve for estimating pangenome size). Blastn instead of blastp was used to report GCs for ANI heat plots. To generate the core and the all shared CDS regions ANI .tab files, we used the command ‵./get_homologues.pl‵ with flags: ‵-d‵, ‵-À, ‵-t‵, ‵-à, ‵-M‵. Furthermore, to plot the ANI heatmaps the ‵./plot_matrix_heatmap.sh‵ command was used.

### Functional analysis of four *Corynebacterium* pangenomes using anvi’o and PPanGGOLiN

The pangenome for each *Corynebacterium* species was analyzed using anvi’o v8 (64, 128). We used a workflow that allowed us to import Prokka annotated genomes into anvi’o (https://klemonlab.github.io/CorPGA_Pangenomics/SupplementalMethods_Prokka_Annotations.html), followed by the addition of functional COG annotations using the ‵anvi-run-ncbi-cogs‵ command with --sensitive flag (runs sensitive version of DIAMOND (129)) and the 2020 updated COG20 database (130, 131). Various annotations were also applied to each genome db file, such as KEGG/KOfam (132, 133), Pfam (134)Mistry, 2021 #3457] and hmm-hits (135). The pangenome for each species was computed with the ‵anvi-pan-genomè program (flags: --mcl-inflation 10, and --use-ncbi-blast) using blastp search (136), muscle alignment (137) ‵minbit heuristic‵ (138) to filter weak hits, and the MCL algorithm (139). A combined pangenome was also computed for 107 strains: the 102 nasal *Corynebacterium* genomes selected for metabolic analysis (Cac_ATCC_49726 & Cps_090104 were excluded) plus 5 *Corynebacterium* listed in **Table S1D**. This high level pangenomic analysis was used for gene cluster level comparisons of the KEGG results. The functional and the geometric homogeneity index, and the rest of the layers shown within the anvi’o pangenome *Corynebacterium* figures were determined from the standard anvi’o pangenomic pipeline (https://merenlab.org/2016/11/08/pangenomics-v2/). The core (genes contained in all genomes), soft core (genes contained in 95% of the genomes), shell (genes contained in several genomes), and cloud (genes present in only a few genomes) assignments from GET_HOMOLOGUES were uploaded into the anvi’o pangenome for each species by creating bins in the anvi’o interactive interface. We also manually rearranged the order of strains in each species-specific analysis to match the order in our species-specific phylogenomic trees and imported our phylogenies into the anvi’o figures. This facilitated visualizing the strain genomes from an evolutionary standpoint and identifying GCs that might be unique to a strain or a group of strains in a clade. The code used to generate the anvi’o pangenomic analysis can be found at https://klemonlab.github.io/CorPGA_Pangenomics/SupplementalMethods_Anvio.html.

We then exported the summary files from the anvi’o pangenomic analyses to synchronize gene cluster identities with PPanGGOLiN v1.2.74 (67) (**Table S2**) (for detailed code see https://klemonlab.github.io/CorPGA_Pangenomics/SupplementalMethods_PPanGGOLiN.html). GCs were defined as persistent or accessory by PPanGGOLiN and then we used an in-house R script (https://github.com/KLemonLab/CorPGA_Pangenomics/blob/main/SupplementalMethods_COGS.Rmd) to clean up and retrieve informative COG20 annotated GCs and to generate the functional enrichment plots shown in **Fig 3** and **Fig S4**.

### Estimation of metabolic capabilities using anvi’o v8

We then used ‵anvi-estimate-metabolism‵ (140) to estimate gene enzymatic functions (**Table S3A**) and the completeness of KEGG modules in each of the four *Corynebacterium* genomes for the selected 102 strains, in each of 27 *D. pigrum* strain genomes (62), in each reference genome of the 9 other species listed in Table S1D, using their original NCBI annotations. The complete KEGG modules estimation output in tabular format is provided in Table S4A, including stepwise and pathwise completeness scores, and, for easier readability, in Table S4B as a matrix summarizing module stepwise completeness by genome. Table S4C includes the module estimations averaged by species. For the data summaries presented in Table 3 and Figure 4, we excluded modules that were complete in < 12.5% of the strains of each species, across all analyzed species. Additional detailed methods and code, including the code for generating Table 3 and Figure 8, are available online (https://klemonlab.github.io/CorPGA_Pangenomics/SupplementalMethods_Anvio.html).

### Determining KEGG module enrichment

We used ‵anvi-display-functions‵ and ‵anvi-compute-metabolic-enrichment‵(141), with a module stepwise completeness score of 1 for the latter, to identify individual KOs (**Table S3B**) and KEGG modules (**Table S4D**) enriched across the four *Corynebacterium* species. Modules with an adjusted q-value < 1e-9 were considered enriched in their associated group. Modules both with an adjusted q-value > 1e-9 and complete in at least 87% of the analyzed genomes were considered shared. Additional detailed methods and code are available online (https://klemonlab.github.io/CorPGA_Pangenomics/SupplementalMethods_Anvio.html).

### Bacterial strains and growth conditions

*C. propinquum* KPL3953, *C. pseudodiphtheriticum* MSK311, KPL1989, and KPL4025, *C. accolens* KPL1818, *C. tuberculostearicum* MSK074, and *C. glutamicum* DSM 20300^T^ were each grown from a - 80 °C freezer stock on Difco Brain Heart Infusion (BHI) agar medium supplemented with 1% Tween80 (BHIT) for 36-48 hours at 34 °C in a 5% CO_2_ CellXpert C170 incubator (Eppendorf, # 6734010015) with a humidification pan. Subsequently, for each strain, we carefully collected 10-20 single colonies from BHIT agar and pooled these for resuspension in individual 5 ml Eppendorf tubes containing 4.5 ml of either phosphate buffered saline (PBS) for the glycogen experiments or Earle’s Balanced Salt Solution (EBSS) for the growth assays with amino acid supplementation.

### MOPS-buffered chemically defined medium base

We used a base chemically defined medium (CDM) buffered with 3-(N-morpholino)propanesulfonic acid (MOPS) into which we supplemented specific concentrations of glucose, varying amino acids, and/or other supplements, as detailed below, to generate sterile chemically defined media. MOPS-buffered base CDM is a 2X solution of a Teknova medium prepared from Teknova 10X MOPS Buffer (M2101), Teknova 10X AGCU solution (M2103), Teknova 100X Vitamin Stock I (V1015), and Teknova 0.132 M Potassium Phosphate Dibasic (M2102) plus 1000X ferric chloride, 400X lipoic acid, and Tween80 with the following final concentrations: 80 mM MOPS, 8 mM tricine, 2.64 mM potassium phosphate dibasic anhydrous, 3 mM potassium hydroxide, 0.398 mM adenine, 0.398 mM cytosine, 0.398 mM uracil, 0.398 mM guanine, 0.02 mM ferrous (Iron II) sulfate heptahydrate, 19 mM ammonium chloride, 0.552 mM potassium sulfate, 0.001 mM calcium chloride dihydrate, 1.05 mM magnesium chloride hexahydrate, 100 mM sodium chloride, 5.84E-07 mM ammonium molybdate, 8.00E-05 mM boric acid, 6.04E-06 mM cobalt chloride hexahydrate, 1.92E-06 mM copper sulfate pentahydrate, 1.62E-05 mM manganese (II) chloride tetrahydrate, 1.95E-06 mM zinc sulfate heptahydrate, 0.2 mM ferric chloride hexahydrate, 0.408 mM choline chloride, 0.406 mM nicotinic acid, 1.46E-02 mM pyridoxine HCl, 3.94E-02 mM calcium pantothenate, 0.3732 mM PABA, 0.1048 mM thiamine HCl, 8.18E-04 mM biotin, 1.48E-05 mM cyanocobalamin, 3.16E-04 mM folic acid, 2.06E-02 mM riboflavin, 0.0192 mM lipoic acid, 0.1% Tween80. The pH of the medium is 7 to 7.5.

### 20 amino acid mix

We prepared a 5X solution mixture of 20 amino acids in MilliQ H_2_O such that the final concentration of each after addition to base CDM was as follows: 1.6 mM L-alanine, 10.4 mM L-arginine, 0.8 mM L-asparagine, 0.8 mM L-aspartic Acid, 0.2 mM L-cysteine, 1.2 mM L-glutamic acid, 1.2 mM L-glutamine, 1.6 mM L-glycine, 0.4 mM L-histidine, 0.8 mM L-isoleucine, 1.6 mM L-leucine, 0.8 mM L-lysine, 0.4 mM L-methionine, 0.8 mM L-phenylalanine, 0.8 mM L-proline, 20 mM L-serine, 0.8 mM L-threonine, 0.2 mM L-tryptophan, 0.4 mM L-tyrosine, 1.2 mM L-valine.

### Determination of intracellular glycogen concentration

Bacteria were resuspended in PBS as described above then washed twice by centrifugation at 16,000 RCF (relative centrifugal force) at 4 °C for 10 minutes using a 5430R centrifuge (Eppendorf # 022620689) and resuspended in 4.5 ml of PBS. Next, we used PBS to adjust the OD_600_ to 2 for each strain and then inoculated 1.25 ml of each as a monoculture into 50 ml base phosphate-buffered CDM supplemented with 5% glucose and the 20 amino acid mix in a 250 ml flask. Cultures were grown in a shaking incubator at 250 RPM and 34 °C. At 24, 28, 32, and 48 hours, we measured the OD_600_ and collected 4 ml aliquots of each strain. Samples were centrifuged, resuspended in PBS, and washed twice as above. After washing, the samples were resuspended in 400 µl PBS and the OD_600_ was measured and adjusted with PBS to ∼ 0.5 to assay an equal biomass per strain. We recorded the OD_600_ values and used these to normalize the assayed glycogen per unit of OD_600_. A 300 µl aliquot of each strain was then combined with 150 µl of HCl in a 2 ml lysis matrix B tube (MP Biomedicals, # 116911050-CF) and heated at 95 °C for 5 minutes. To lyse the bacteria, we bead beat the samples 3 times at 6.5 m/s for 45 seconds each time using a FastPrep-24 5G (MP Biomedicals # 116005500). After mechanical lysis, 150 µl of Tris buffer was added to each tube. The samples were then centrifuged at 16,000 RCF at 4°C for 20 minutes. The supernatant was recovered and 25 µl of each sample was aliquoted into 6 separate wells per strain in a white 96-well plate with clear bottom (Greiner Bio-One # 655074) and assayed for glycogen with the Glycogen-Glo Assay kit (Promega # J5051). Per the manufacturer’s instructions, we prepared a glycogen titration curve with two-fold serial dilutions in PBS from 20 to 0 µg/ml and we added 25 µl of each of these standards to the 96-well plate for each time point: one set of standards for the glucoamylase assay and one as the control. We then added either 25 µl of glucoamylase solution or buffer alone to each sample (three technical replicates each) and standard well. Plates were incubated at room temp for one hour to allow the glucoamylase to digest glycogen into free glucose. Following incubation, 50 µl of glucose detection reagent was added to each well. Luminescence was measured after 70 minutes using a BioTek Synergy H1 microplate reader (Agilent # SH1M-SN) with default settings. The average luminescence for glucoamylase-treated and buffer-only samples was calculated from three technical replicates for each strain at each timepoint for each independent experiment. The average value from the buffer-only samples was subtracted from the glucoamylase-treated samples for each sample to correct for background levels of free glucose. The glycogen titration curve was derived by subtracting the luminescence of the buffer-only negative controls from the glucoamylase positive standards to establish a best-fit regression curve. The average differences in luminescence values from bacterial samples with and without glucoamylase treatment were input into the corresponding best-fit curve equation to determine total glycogen concentration (µg/ml) per sample. These were normalized to µg/ml per OD_600_ of the bacterial suspension measured before mechanical lysis to approximate the glycogen concentration per cell biomass of each strain. Statistics were performed using a linear mixed model with species as the fixed effect and time point as a random effect fit by restricted maximum likelihood with t-tests using Satterthwaite’s method with R packages lme4 (142) and afex. Additional detailed methods and code, including the code for generating Figure 5A-B, are available online https://klemonlab.github.io/CorPGA_Pangenomics/SupplementalMethods_Glycogen.html.

### Sequence alignment for the *fpr2-cysIXHDNYZ* operon

We manually inspected KO assignments for all the enzymes encoded in the *C. glutamicum fpr2-cysIXHDNYZ* operon (89) that absent from the KEGG definition of the assimilatory sulfate reduction module (M00176). Genes annotated as K00528 (*fprA*) were identified in all the *Corynebacterium* genomes. A manual exploration of the gene annotations in the genomic region around the putative *fprA* gene in all *C. pseudodiphtheriticum* strains identified *hchA* as a common gene downstream the *fpr2-cysIXHDNYZ* operon. We used BEDTools (143, 144) to extract the genomic sequence flanked by *fprA* and *hchA* and performed multiple sequence alignment using MAFFT (145, 146). AliView (147) was used for sequence inspection and generation of a reverse-complemented alignment. The final visualization in Figure 6A was generated in R with the package msavisr. Additional detailed methods and code, are available online https://klemonlab.github.io/CorPGA_Pangenomics/SupplementalMethods_Anvio.html

### Assay for growth under conditions requiring assimilatory sulfate reduction

We generated a phosphate-buffered version of base CDM with 75 mM (f.c.) sodium sulfate as the only source of sulfur (KS-CDM) by replacing the Teknova MOPS buffer with a phosphate buffer consisting of potassium phosphate monobasic, potassium phosphate dibasic, and sodium phosphate dibasic and trace minerals (see below) since *C. glutamicum* can reportedly use MOPS as a source of sulfur (148). We supplemented the KS-CDM base with 2% glucose (Teknova 20% Glucose solution G0520) and an 18 amino acid mix, excluding cysteine and methionine, to generate KS-CDM 0.76% agarose. We also assayed for growth on KS-CDM agarose supplemented with a 75X cysteine and methionine solution to final concentrations of 2 mM cysteine and 4 mM methionine. Both conditions were tested with and without lipoic acid, as a potential additional sulfur source. Prior to inoculation for each experiment, each bacterial strain resuspended in EBSS (as described above) was centrifuged at 10,000 RCF for 10 minutes and then subjected to three washes of 4.5 ml EBSS. After the final wash, cells were resuspended in 500 µl EBSS to achieve a visibly turbid solution and 15 µl from the resuspension of each strain was inoculated and struck for single colonies onto KS-CDM 0.76% agarose (SeaKem LE Agarose, # 50004) prepared with the different variations and incubated at 34 °C in 5% CO2 incubator with a humidification pan. We used agarose because of its higher purity compared to agar. The KS-CDM agarose plates were imaged at 8 days to assess bacterial growth with each supplementation. We performed four independent experiments (**Figs. 6** and **S5A**). KS-CDM is a 2X solution of a Teknova medium prepared from Teknova 10X AGCU solution (M2103), and Teknova 100X Vitamin Stock I (V1015), plus 50000X Trace Minerals I and 50X Trace Minerals II (File S1), 1000X ferric chloride, 1000X ferrous chloride, 10X phosphates, 10X NaCl, 100X tricine, 25X sodium sulfate, 400X lipoic acid, and Tween80. Final concentrations in KS-CDM were as follows: 37 mM potassium phosphate monobasic, 109 mM potassium phosphate dibasic, 63 mM sodium phosphate dibasic, 0.398 mM adenine, 0.398 mM cytosine, 0.398 mM uracil, 0.398 mM guanine, 8 mM tricine, 75mM sodium sulfate, 6.04E-06 mM cobalt chloride hexahydrate, 1.92E-06 mM copper chloride dihydrate, 1.62E-05 mM manganese chloride tetrahydrate, 5.84E-07 mM ammonium molybate, 1.95E-06 mM zinc chloride, 0.00008 mM boric acid, 19 mM ammonium chloride, 0.001 mM calcium chloride, 1.05 mM magnesium chloride, 0.2 mM ferric (Iron III) chloride hexahydrate, 0.02 mM ferrous (Iron II) chloride tetrahydrate, 35 mM sodium chloride, 1.6 mM L-alanine, 10.4 mM L-arginine, 0.8 mM L-asparagine, 0.8 mM L-aspartic Acid, 1.2 mM L-glutamic Acid, 1.2 mM L-glutamine, 1.6 mM L-glycine, 0.4 mM L-histidine, 0.8 mM L-isoleucine, 1.6 mM L-leucine, 0.8 mM L-lysine, 0.8 mM L-phenylalanine, 0.8 mM L-proline, 20 mM L-serine, 0.8 mM L-threonine, 0.2 mM L-tryptophan, 0.4 mM L-tyrosine, 1.2 mM L-valine, 0.1% Tween80.

### Amino acid auxotrophy assays

To test for amino acid auxotrophy, we used MOPS-buffered base CDM 0.76% agarose supplemented with 2% glucose and either lacking amino acids (0 AA) or supplemented with one of the following: the 20 amino acid mix, 5 mM Urea, 10 mM glutamine, or 10 mM asparagine. Bacterial inocula were prepared and growth assays were performed as described above in testing for assimilatory sulfate reducing ability. The MOPS-CDM agarose plates were imaged at 8 days to assess bacterial growth with each supplementation. We performed four independent experiments (**Figs. 7** and **S5B**).

## Data and code availability

All genomes are available from NCBI in Bioprojects <u>PRJNA842433</u> and <u>PRJNA804245</u>. Table S1 lists genome accession numbers. Code used is in our GitHub repository https://github.com/KLemonLab/CorPGA_Pangenomics.

## Acknowledgments

Thank you to all the individuals who donated nostril swab samples at a 2017 and 2018 science festival that were used to isolate KPL strains of *Corynebacterium* species. Their contributions continue to expand knowledge about human nasal microbiota. Thank you to lab members and colleagues who contributed to these outreach events, including Javier Fernandez Juarez, Kerry Maguire, Genevieve Holmes, Pallavi Murugkar, Pooja Balani, Sowmya Balasubramanian, Fan Zhu, Andrew Collins, Andy Kempczynski, Brian Klein, and Megan Lambert. For early planning and initial cultivation efforts we thank Silvio D. Brugger, Lindsey Bomar, Stephany Flores Ramos, and Sara M. Eslami for KPL strains; and Jhoanna N. Aquino and Christopher R. Polage for MSK strains. We also thank all of the individuals in the USA and Botswana who participated in the research studies that resulted in cultivation of the MSK strains. Thank you to Clay Deming and the NIH Intramural Sequencing Center for sequencing of the KPL strains. Thank you to Bruno Contreras-Moreira for advice on using GET_HOMOLOGUES; to A. Murat Eren for advice on using anvi’o; and to Ashlee M. Earl for early efforts to genome sequence nasal *Corynebacterium* isolates. Thank you to other members of our labs.

## Authorship contributions

Conceptualization: KPL, WG, IFE. New Strain isolation: WG. Investigation: THT, IFE, AQR, ACO, WG, SC, MSK. Bioinformatic analysis: THT, IFE, AQR, SC. Interpretation of data: THT, IFE, AQR, ACO, WG, JAS, HHK, SC, MSK, KPL. Visualization: THT, IFE, ACO, AQR. Wrote Original Draft: THT, AQR, IFE, ACO, KPL. Editing and review: KPL, MSK, SC, IFE, THT, ACO, AQR, WG, HHK, JAS. All authors read and approved the final manuscript. Supervision: KPL, IFE, MSK, HHK, JAS. Funding Acquisition: KPL, MSK, HHK, JAS.

## SUPPLEMENTAL MATERIALS

**File S1 (text).** Supplemental text

**Table S2 (excel file).** PPanGGOLiN assignment of GCs to the persistent or accessory genome

**Table S3 (excel file).** KO metabolic estimations and enrichment analysis by anvi’o

**Table S4 (excel file).** KEGG module estimations and enrichment analysis by anvi’o

## Supplemental File S1

### Species assignment for *Corynebacterium* sp. KPL# previously submitted to NCBI

The Broad Institute originally sequenced a set of *Corynebacterium* strains we had isolated from human nostril swabs, and we included in our current analyses any of these that were assigned to one of the four species of interest (113). The species assignment for each of these based on phylogenomic analyses (**Figs. 1** and **S1**) and ANI (**Figs. S2A-B**) are as follows: *C. accolens* KPL1818; *C. accolens* KPL1824; *C. accolens* KPL1996 (*C. accolens* KPL1986, KPL1998, and KPL2004); and *C. pseudodiphtheriticum* KPL1989 (*C. pseudodiphtheriticum* KPL1995). Isolates in parentheses are genomes listed in NCBI that have a MASH distance ≤ 10^-4^ to the strain genome listed immediately before the parentheses.

### Comparing core genomes and pangenomes predicted by GET_HOMOLOGUES and anvi’o

The difference in the outputs of gene clusters (GCs) from GET_HOMOLOGUES (consensus of OMCL and COGS algorithms) and anvi’o were measured using the percent difference formula 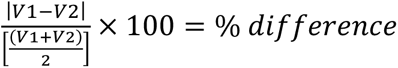. Core and pangenomes sizes for both the pangenomic platforms are listed below for comparison, with their percentage differences.

| Species | Core genome |  |  | Pangenome |  |  |
| --- | --- | --- | --- | --- | --- | --- |
|  | GH | anvi'o | % diff. | GH | anvi'o | % diff. |
| *C. propinquum* | 1624 | 1824 | 11.6% | 3777 | 3108 | 19.4% |
| *C. pseudodiphtheriticum* | 1515 | 1714 | 12.3% | 4590 | 3590 | 24.4% |
| *C. accolens* | 1688 | 1904 | 12.0% | 4220 | 3427 | 20.7% |
| *C. tuberculostearicum* | 1810 | 1915 | 5.6% | 3232 | 2907 | 10.6% |

### L-Glutamine is also an excellent nitrogen source in *C. glutamicum*

Although the GOGAT enzyme is the main pathway involved in glutamine assimilation, it is not required for this, since in a *C. glutamicum* GOGAT deletion mutant, a glutaminase in conjunction with glutamate dehydrogenase appears to contribute to glutamine utilization, although primarily as a carbon and energy source (149, 150) In all the analyzed *Corynebacterium* genomes, except for *C. diphtheriae^T^,* we identified genes annotated as a glutaminase (K01425) corresponding to the *C. glutamicum*^T^ *glsK* gene, which hydrolyzes glutamine producing ammonium and glutamate. Glutamine is also essential to produce carbamoyl phosphate, a precursor for the arginine and pyrimidine synthesis pathways. In *C. glutamicum* the glutamine-dependent carbamoyl phosphate synthetase II (CPS_II) encoded by the *carAB* operon converts HCO_3_^-^, ATP, and glutamine to carbamoyl phosphate (151). We identified K01956 and K01955 (*carA* and *carB*) annotations in all analyzed *Corynebacterium* species. Based on these findings, we predicted that addition of glutamine might improve the growth of nasal *Corynebacterium* species in defined medium lacking amino acids via enzymatic activities that use glutamine as a precursor, including, but probably not limited to, glutaminase, carbamoyl phosphate, and/or glutamine amidotransferase enzymes.

### Trace Minerals Solutions

The Trace Minerals I 50000X stock consists of the following: 0.302 mM cobalt chloride hexahydrate, 0.096 mM copper chloride dihydrate, 0.81 mM manganese chloride tetrahydrate, 0.0292 mM ammonium molybdate, 0.0975 mM zinc chloride, 4 mM boric acid. The Trace Minerals II 50X stock consists of the following: 950 mM ammonium chloride, 0.05 mM calcium chloride, 52.5 mM magnesium chloride.

### Main and Supplemental Figures (in order of appearance in text)

**Figure S1.**
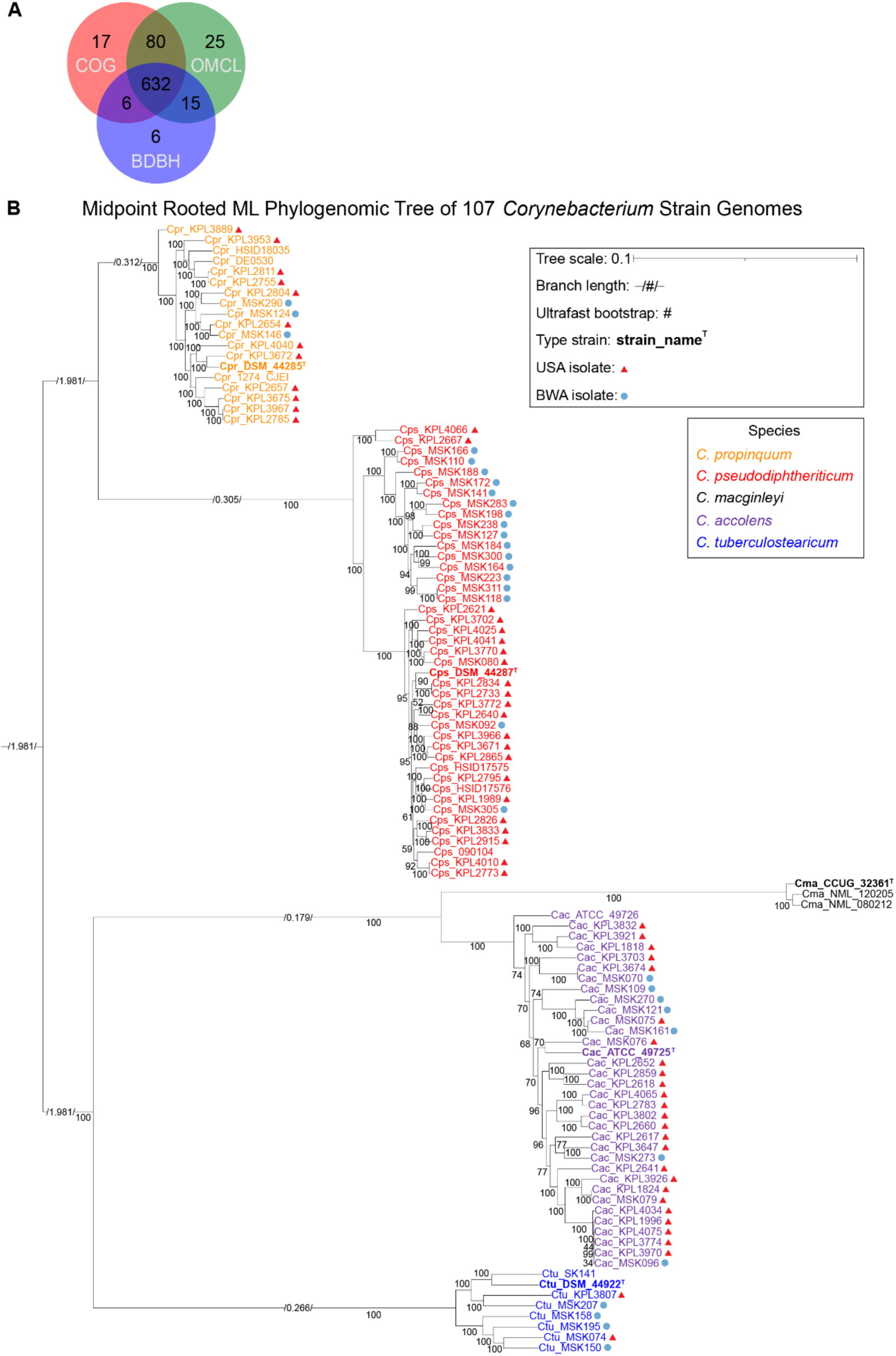
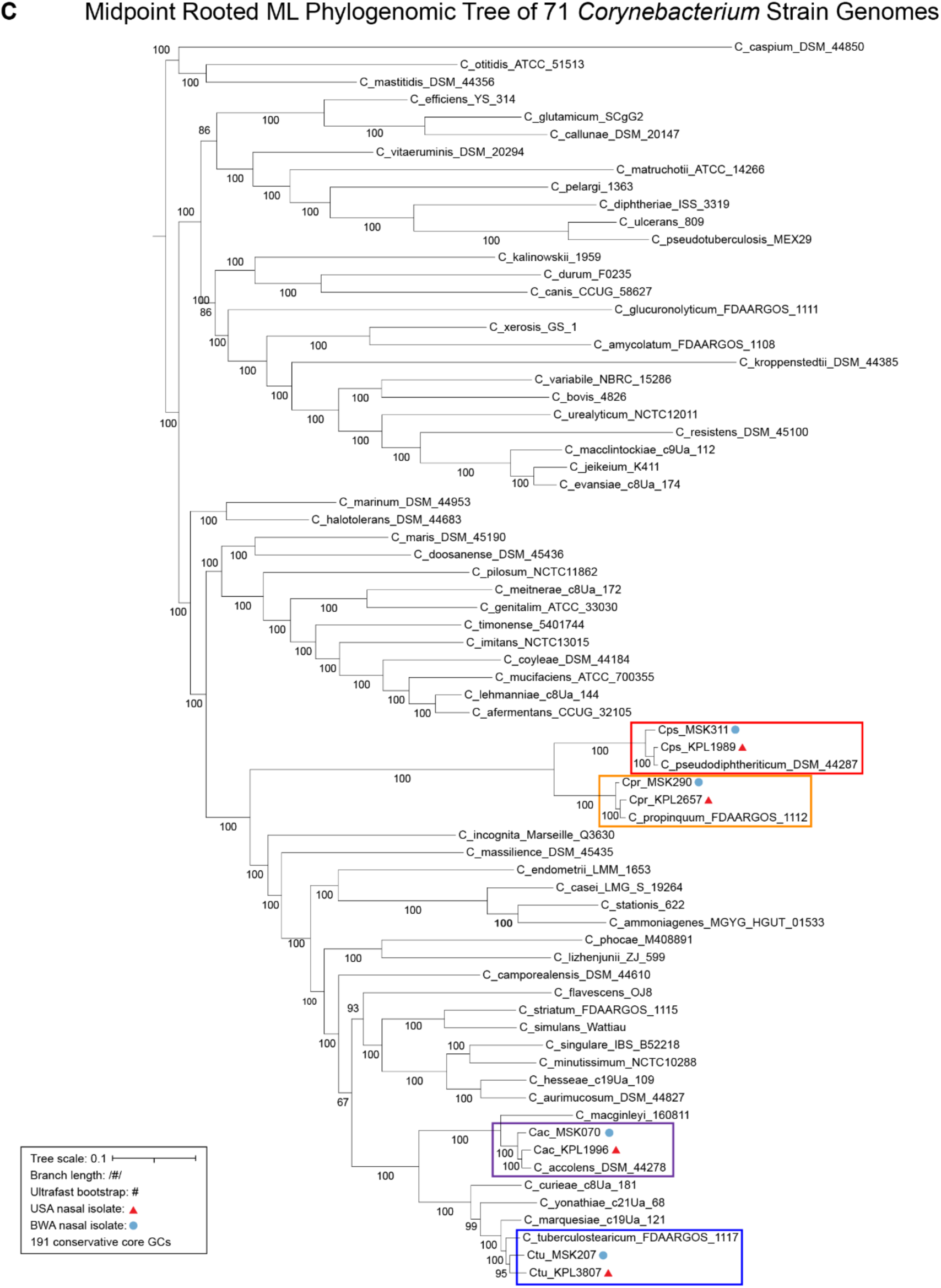
Phylogenomic trees, generated from the shared conservative core gene clusters, illustrate the evolutionary relationships among four human-nasal-associated *Corynebacterium* species and their relationship to other species within this genus. (**A**) Venn diagram showing 632 single-copy GCs are shared between 107 *Corynebacterium* strain genomes based on the consensus of BDBH, COG triangle, and OMCL algorithms using GET_HOMOLOGUES v24082022. (**B**) A maximum-likelihood phylogenomic tree constructed from 632 concatenated shared single-copy GCs from 107 *Corynebacterium* strain genomes. Species are indicated by name and color-code (see key) with type strains highlighted in bold. We used IQ-Tree v2.1.3 with model finder (BIC value 11679476.6345), edge-linked-proportional partition model, and 1,000 ultrafast rapid bootstraps (values on tree). Branch lengths are indicated (see key). The phylogeny contains 31 and 56 nasal-isolated strain genomes from Botswana (blue circles) and the USA (red triangles). Many branches are highly supported with ultrafast bootstrap values ≥ 95. (**C**) A maximum-likelihood phylogenomic tree of 63 *Corynebacterium* species illustrates the relationship of the four nasal species to other members of the genus. *C. propinquum*, *C. pseudodiphtheriticum*, *C. accolens*, and *C. tuberculostearicum* are each highlighted by a colored box. This is based on 191 shared conservative core GCs from 71 *Corynebacterium* strain genomes. Representative nasal strain genomes from Botswana (blue circle) and the USA (red triangle) are shown for each of the four species.

**Figure S2.**
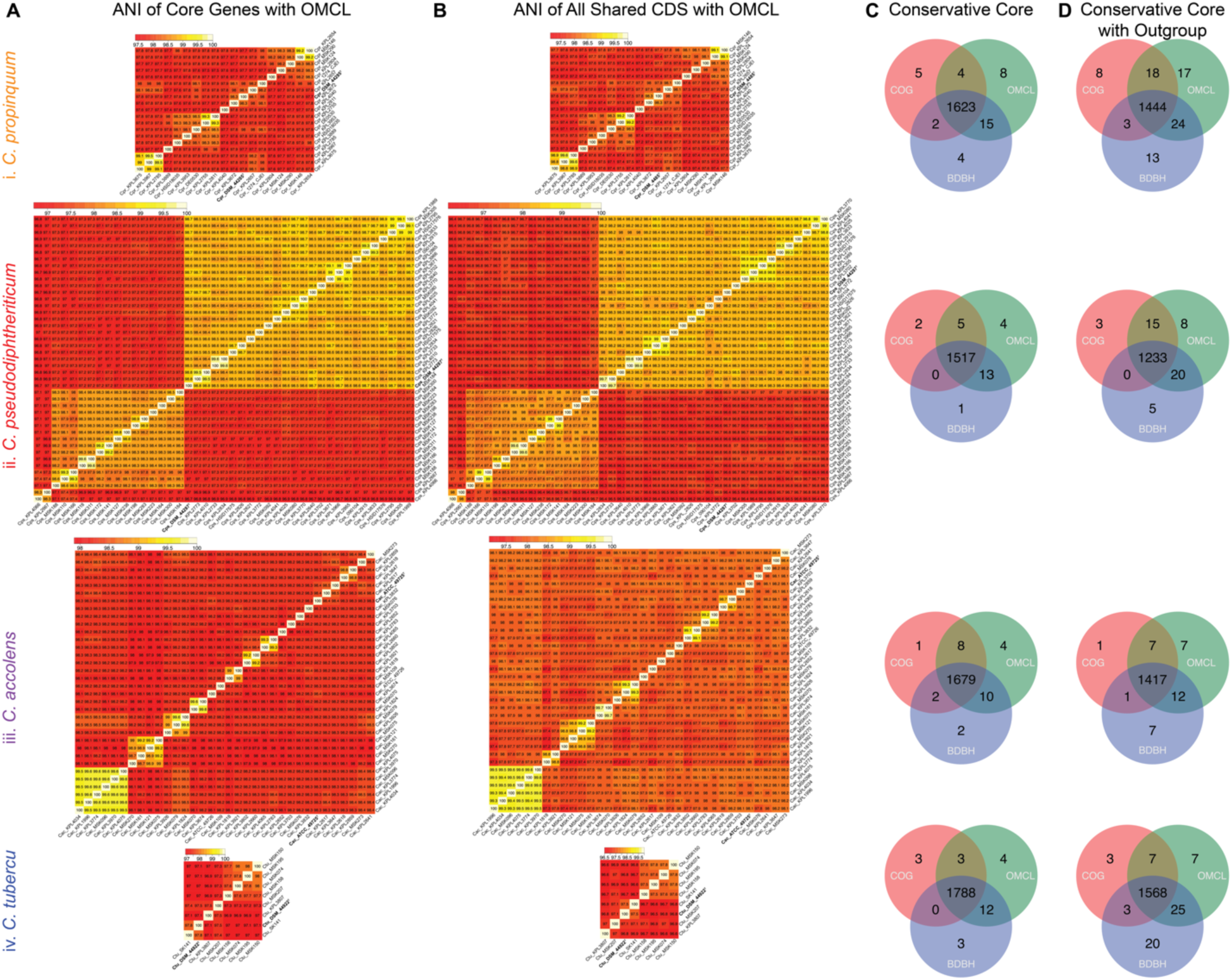
Species-level pairwise average nucleotide identities and conservative core GCs. (**A**) The shared core genes among all paired strain genomes within each species (**i-iv**) have ≥ 95% ANI (average nucleotide identity) to the type strain using the OMCL algorithm. (**B**) The ANI pairwise comparisons for all shared CDS regions are slightly less than core ANI values but still ≥ 95% among strain genomes within each species. The order of strains in A and B differ due to changes in pairwise identities because more SNPs were included in B. (**C**) The shared conservative core single-copy GCs determined by the consensus of BDBH, COG triangle, and OMCL for each of the *Corynebacterium* species. (**D**) The conservative core CGs for each species plus the type strain of its closest relative in the five-species phylogenomic tree (**Fig. S1B**). Between 112 and 240 GCs were absent from the conservative core of the outgroup for each species, with the fewest absent in the outgroup for *C. pseudodiphtheriticum* and the most absent in the outgroup for *C. tuberculostearicum*.

**Figure S3.**
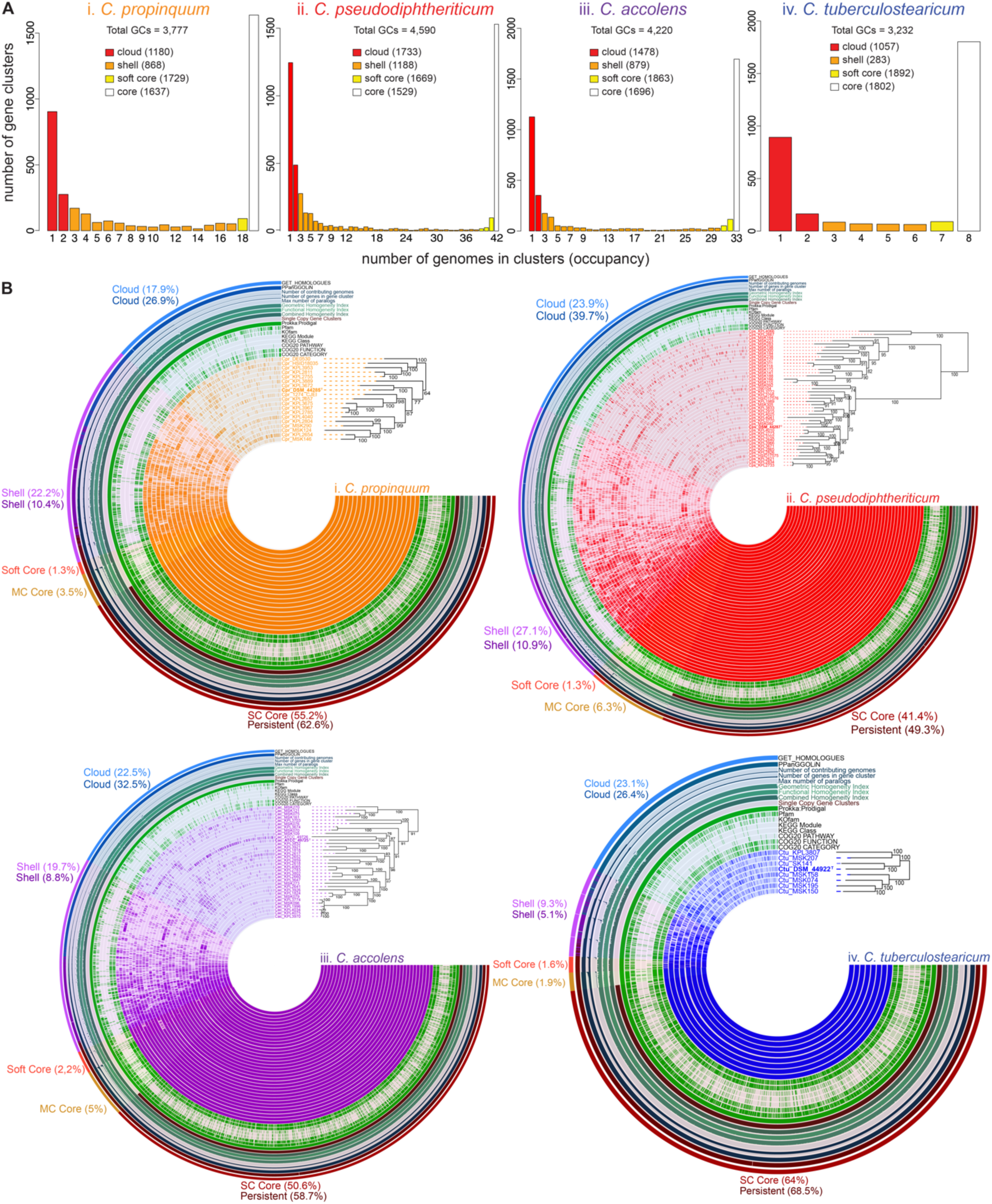
Three different analysis platforms estimated a similar pangenome for each individual *Corynebacterium* species. (**A**) Representation of the pangenome estimated by GET_HOMOLOGUES with COG triangle and OMCL of (**i**) *C. propinquum*, 19 genomes; (**ii)** *C. pseudodiphtheriticum*, 42 genomes; (**iii**) *C. accolens*, 33 genomes; and (**iv**) *C. tuberculostearicum,* 8 genomes. (**B**) Using pangenomes clustered with anvi’o, we constructed a concentric dendrogram for each species (**i-vi**) ordered to match the species-specific phylogenies in Figure 1. The GCs present in the core/persistent genome consistently accounted for over half of the pangenome in each of the nasal *Corynebacterium* species. The outermost concentric ring indicates anvi’o-defined GCs (with percentages) in the single copy core (dark red), multicopy core (beige), soft core (red), shell (light purple), and cloud (light blue) based on GET_HOMOLOGUES definition of each partition with anvi’o allowing further division of the core into single copy and multicopy core. The next to outermost ring indicates anvi’o-defined GCs (with percentages) assigned by PPanGGOLiN to persistent (dark maroon), shell (purple), and cloud (blue). The average core+soft core or persistent, shell, and cloud sizes across the four species using GET_HOMOLOGUES definitions were 58.58%, 19.58%, and 21.85%, respectively, compared to 56.63%, 10.13%, and 31.38% as assigned by PPanGGOLiN. The differences between the two algorithms were 1.95% for core, 9.45% for shell, and 9.53% for cloud. GET_HOMOLOGUES defines the pangenome compartments as GCs shared across a certain number of genomes, whereas PPanGGOLiN uses both presence and analysis for genomic neighborhoods to infer GCs in the persistent genome, which likely explains these differences.

**Figure S4.**
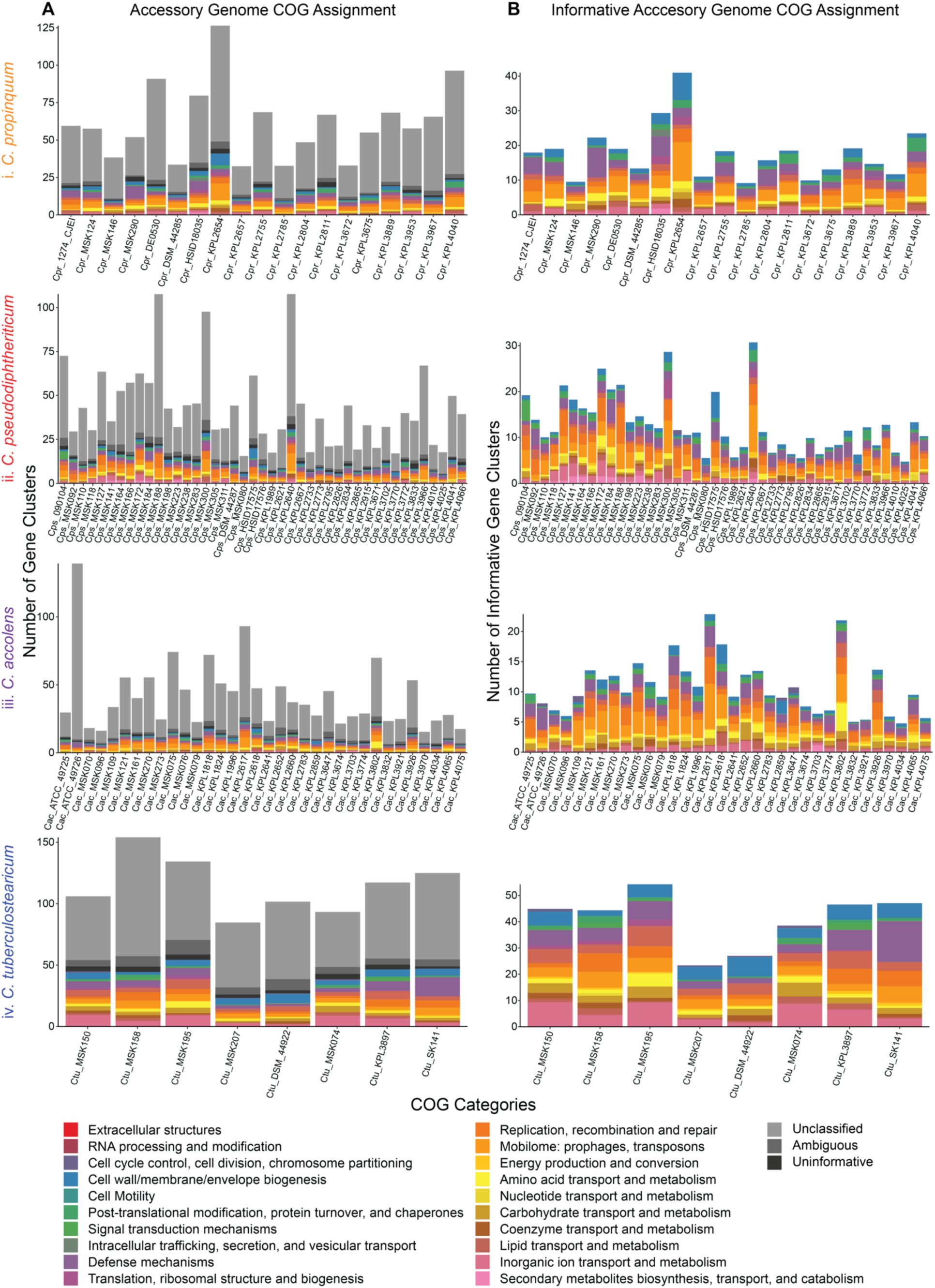
The size of the accessory genome varies between strains within each nasal *Corynebacterium* species. We used anvi’o to identify COG functional annotations and PPanGGOLiN to assign GCs to the persistent vs. accessory genome. (**A**) For each species (**i-vi**), most of the GCs in each strain’s accessory genome had either no annotation (“unclassified”), an ambiguous COG categorization (“ambiguous”), or belonged to an uninformative S or R COG category (“uninformative”). The calculated average percentages for unclassified, ambiguous, and uninformative categories per species were (**i**) 65.2%, 3.5%, and 2.4% for *C. propinquum*; (**ii**) 62.8%, 4.4%, and 2.9% for *C. pseudodiphtheriticum*; (**iii**) 67.6%, 4.7%, and 2.0% for *C. accolens*; and (**iv**) 55.1%, 6.2%, and 3.1% for *C. tuberculostearicum.* The accessory genome of *C. accolens* had the highest percentage of COGS belonging to these three categories and *C. tuberculostearicum* had the lowest. Averages of the informative assigned COG annotations (colored categories) in the accessory genome were calculated for each species as follows: (**i**) 28.9% for *C*. *propinquum*, (**ii**) 29.9% for *C. pseudodiphtheriticum*, (**iii**) 25.7% for *C. accolens*, and (**iv**) 35.6% for *C. tuberculostearicum*. (**B**) There was a relatively proportional distribution of accessory informative functional GCs among each set of genomes. The size of the accessory genome varied between strains from each of the four species. (**Ai**) For *C. propinquum,* KPL2654 and KPL4040 had the largest accessory sizes, whereas KPL2785 and KPL3672 had the smallest. (**Aii**) For *C. pseudodiphtheriticum,* KPL2640 and MSK188 had the largest accessory sizes, whereas MSK080 and KPL3070 had the smallest. (**Aiii**) For *C. accolens,* ATCC_49726 and KPL2617 had the largest accessory sizes, whereas MSK096 and KPL3970 had the smallest. (**Aiv**) For *C. tuberculostearicum* MSK158 and MSK195 had the largest accessory sizes, whereas MSK207 and MSK074 had the smallest.

**Figure S5.**
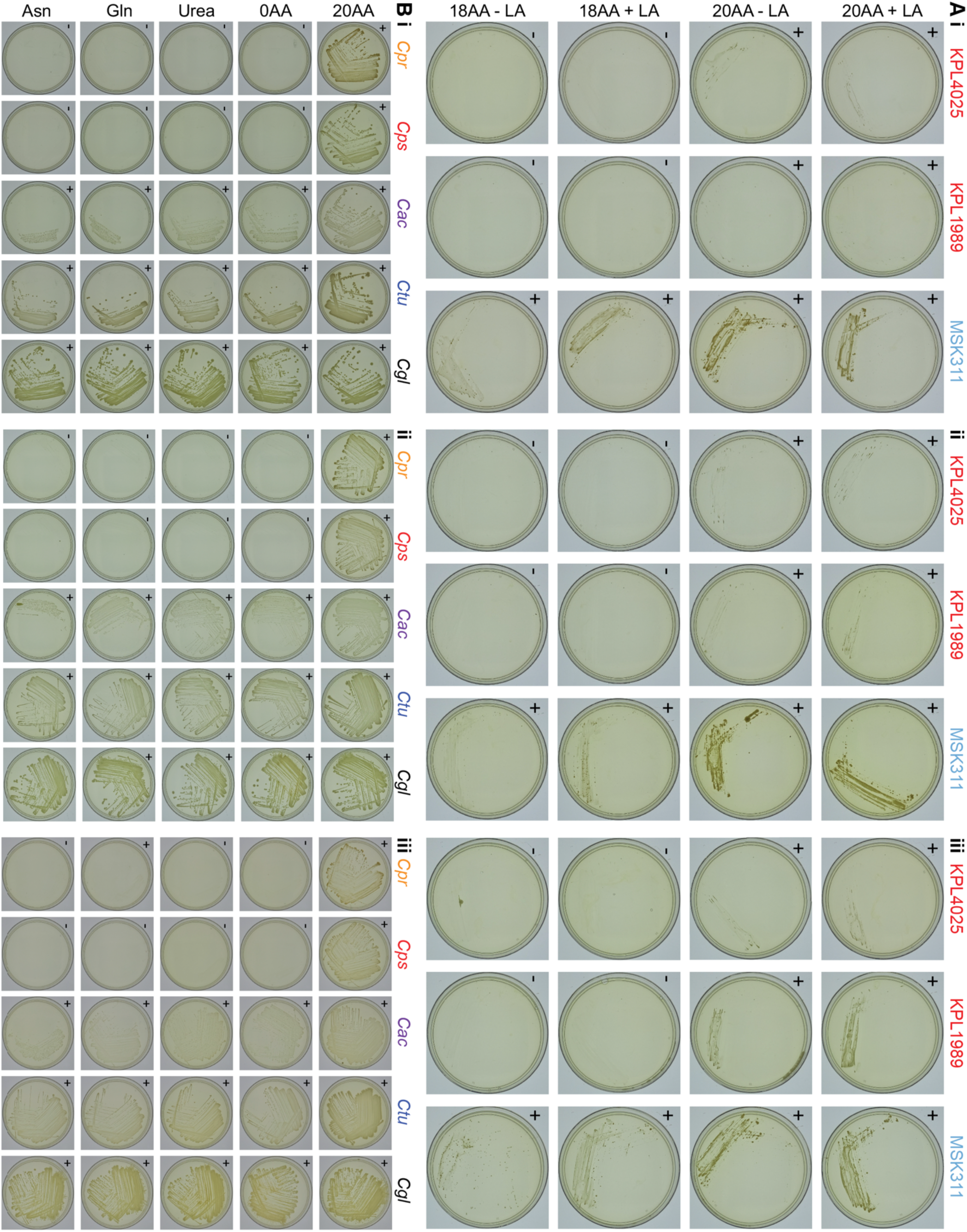
Results of additional growth experiments from Figures 6 (A) and 7 (B). **(A)** The Botswanan *C. pseudodiphtheriticum* strain MSK311, which encodes the *fpr2cysIXHDNYZ* operon, grew on chemically defined agarose medium with sulfate as the only source of sulfur (KS-CDM 18AA - LA), whereas the USA strains KPL1989 and KPL4025, which lack this operon, did not. (**B**) *C. accolens* (*Cac*) and *C. tuberculostearicum* (*Ctu*), as well as *C. glutamicum* (*Cgl*), grew on MOPS-buffered CDM agarose medium in the absence of all 20 amino acids, whereas *C*. *propinquum* (*Cpr*) and *C*. *pseudodiphtheriticum* (*Cps*) did not. All images captured after 8 days of growth at 34°C with 5% CO_2_ with a humidification pan. (+) = growth, (-) = no growth.

## REFERENCES

1. Biesbroek G, Bosch AA, Wang X, Keijser BJ, Veenhoven RH, Sanders EA, Bogaert D. 2014. The impact of breastfeeding on nasopharyngeal microbial communities in infants. Am J Respir Crit Care Med 190:298–308.

2. Biesbroek G, Tsivtsivadze E, Sanders EA, Montijn R, Veenhoven RH, Keijser BJ, Bogaert D. 2014. Early respiratory microbiota composition determines bacterial succession patterns and respiratory health in children. Am J Respir Crit Care Med 190:1283–92.

3. Mika M, Mack I, Korten I, Qi W, Aebi S, Frey U, Latzin P, Hilty M. 2015. Dynamics of the nasal microbiota in infancy: a prospective cohort study. J Allergy Clin Immunol 135:905–912 e11.

4. Teo SM, Mok D, Pham K, Kusel M, Serralha M, Troy N, Holt BJ, Hales BJ, Walker ML, Hollams E, Bochkov YA, Grindle K, Johnston SL, Gern JE, Sly PD, Holt PG, Holt KE, Inouye M. 2015. The infant nasopharyngeal microbiome impacts severity of lower respiratory infection and risk of asthma development. Cell Host Microbe 17:704–15.

5. Peterson SW, Knox NC, Golding GR, Tyler SD, Tyler AD, Mabon P, Embree JE, Fleming F, Fanella S, Van Domselaar G, Mulvey MR, Graham MR. 2016. A Study of the Infant Nasal Microbiome Development over the First Year of Life and in Relation to Their Primary Adult Caregivers Using cpn60 Universal Target (UT) as a Phylogenetic Marker. PLoS One 11:e0152493.

6. Bosch AATM, Levin E, Van Houten MA, Hasrat R, Kalkman G, Biesbroek G, De Steenhuijsen Piters WAA, De Groot P-KCM, Pernet P, Keijser BJF, Sanders EAM, Bogaert D. 2016. Development of Upper Respiratory Tract Microbiota in Infancy is Affected by Mode of Delivery. EBioMedicine 9:336–345.

7. Bosch A, de Steenhuijsen Piters WAA, van Houten MA, Chu M, Biesbroek G, Kool J, Pernet P, de Groot PCM, Eijkemans MJC, Keijser BJF, Sanders EAM, Bogaert D. 2017. Maturation of the Infant Respiratory Microbiota, Environmental Drivers, and Health Consequences. A Prospective Cohort Study. Am J Respir Crit Care Med 196:1582–1590.

8. Shilts MH, Rosas-Salazar C, Tovchigrechko A, Larkin EK, Torralba M, Akopov A, Halpin R, Peebles RS, Moore ML, Anderson LJ, Nelson KE, Hartert TV, Das SR. 2016. Minimally Invasive Sampling Method Identifies Differences in Taxonomic Richness of Nasal Microbiomes in Young Infants Associated with Mode of Delivery. Microb Ecol 71:233–42.

9. Chu DM, Ma J, Prince AL, Antony KM, Seferovic MD, Aagaard KM. 2017. Maturation of the infant microbiome community structure and function across multiple body sites and in relation to mode of delivery. Nat Med 23:314–326.

10. Salter SJ, Turner C, Watthanaworawit W, de Goffau MC, Wagner J, Parkhill J, Bentley SD, Goldblatt D, Nosten F, Turner P. 2017. A longitudinal study of the infant nasopharyngeal microbiota: The effects of age, illness and antibiotic use in a cohort of South East Asian children. PLoS Negl Trop Dis 11:e0005975.

11. Ta LDH, Yap GC, Tay CJX, Lim ASM, Huang CH, Chu CW, De Sessions PF, Shek LP, Goh A, Van Bever HPS, Teoh OH, Soh JY, Thomas B, Ramamurthy MB, Goh DYT, Lay C, Soh SE, Chan YH, Saw SM, Kwek K, Chong YS, Godfrey KM, Hibberd ML, Lee BW. 2018. Establishment of the nasal microbiota in the first 18 months of life: Correlation with early-onset rhinitis and wheezing. J Allergy Clin Immunol 142:86–95.

12. Toivonen L, Hasegawa K, Waris M, Ajami NJ, Petrosino JF, Camargo CA, Jr., Peltola V. 2019. Early nasal microbiota and acute respiratory infections during the first years of life. Thorax 74:592–599.

13. Kelly MS, Plunkett C, Yu Y, Aquino JN, Patel SM, Hurst JH, Young RR, Smieja M, Steenhoff AP, Arscott-Mills T, Feemster KA, Boiditswe S, Leburu T, Mazhani T, Patel MZ, Rawls JF, Jawahar J, Shah SS, Polage CR, Cunningham CK, Seed PC. 2022. Non-diphtheriae Corynebacterium species are associated with decreased risk of pneumococcal colonization during infancy. ISME J 16:655–665.

14. Yan M, Pamp SJ, Fukuyama J, Hwang PH, Cho DY, Holmes S, Relman DA. 2013. Nasal microenvironments and interspecific interactions influence nasal microbiota complexity and *S. aureus* carriage. Cell host & microbe 14:631–40.

15. Ramakrishnan VR, Feazel LM, Gitomer SA, Ir D, Robertson CE, Frank DN. 2013. The microbiome of the middle meatus in healthy adults. PLoS One 8:e85507.

16. Biswas K, Hoggard M, Jain R, Taylor MW, Douglas RG. 2015. The nasal microbiota in health and disease: variation within and between subjects. Front Microbiol 9:134.

17. Kaspar U, Kriegeskorte A, Schubert T, Peters G, Rudack C, Pieper DH, Wos-Oxley M, Becker K. 2016. The culturome of the human nose habitats reveals individual bacterial fingerprint patterns. Environ Microbiol 18:2130–42.

18. Wos-Oxley ML, Chaves-Moreno D, Jauregui R, Oxley AP, Kaspar U, Plumeier I, Kahl S, Rudack C, Becker K, Pieper DH. 2016. Exploring the bacterial assemblages along the human nasal passage. Environ Microbiol 18:2259–71.

19. De Boeck I, Wittouck S, Wuyts S, Oerlemans EFM, van den Broek MFL, Vandenheuvel D, Vanderveken O, Lebeer S. 2017. Comparing the Healthy Nose and Nasopharynx Microbiota Reveals Continuity As Well As Niche-Specificity. Front Microbiol 8:2372.

20. Laufer AS, Metlay JP, Gent JF, Fennie KP, Kong Y, Pettigrew MM. 2011. Microbial communities of the upper respiratory tract and otitis media in children. mBio 2:e00245–10.

21. Pettigrew MM, Laufer AS, Gent JF, Kong Y, Fennie KP, Metlay JP. 2012. Upper respiratory tract microbial communities, acute otitis media pathogens, and antibiotic use in healthy and sick children. Appl Environ Microbiol 78:6262–70.

22. Bomar L, Brugger SD, Yost BH, Davies SS, Lemon KP. 2016. *Corynebacterium accolens* Releases Antipneumococcal Free Fatty Acids from Human Nostril and Skin Surface Triacylglycerols. mBio 7:e01725–15.

23. Kelly MS, Surette MG, Smieja M, Pernica JM, Rossi L, Luinstra K, Steenhoff AP, Feemster KA, Goldfarb DM, Arscott-Mills T, Boiditswe S, Rulaganyang I, Muthoga C, Gaofiwe L, Mazhani T, Rawls JF, Cunningham CK, Shah SS, Seed PC. 2017. The Nasopharyngeal Microbiota of Children With Respiratory Infections in Botswana. Pediatr Infect Dis J 36:e211–e218.

24. Hasegawa K, Linnemann RW, Mansbach JM, Ajami NJ, Espinola JA, Petrosino JF, Piedra PA, Stevenson MD, Sullivan AF, Thompson AD, Camargo CA, Jr. 2017. Nasal Airway Microbiota Profile and Severe Bronchiolitis in Infants: A Case-control Study. Pediatr Infect Dis J 36:1044–1051.

25. Kelly MS, Surette MG, Smieja M, Rossi L, Luinstra K, Steenhoff AP, Goldfarb DM, Pernica JM, Arscott-Mills T, Boiditswe S, Mazhani T, Rawls JF, Cunningham CK, Shah SS, Feemster KA, Seed PC. 2018. Pneumococcal Colonization and the Nasopharyngeal Microbiota of Children in Botswana. Pediatr Infect Dis J 37:1176–1183.

26. Lappan R, Imbrogno K, Sikazwe C, Anderson D, Mok D, Coates H, Vijayasekaran S, Bumbak P, Blyth CC, Jamieson SE, Peacock CS. 2018. A microbiome case-control study of recurrent acute otitis media identified potentially protective bacterial genera. BMC Microbiol 18:13.

27. Man WH, Clerc M, de Steenhuijsen Piters WAA, van Houten MA, Chu M, Kool J, Keijser BJF, Sanders EAM, Bogaert D. 2019. Loss of Microbial Topography between Oral and Nasopharyngeal Microbiota and Development of Respiratory Infections Early in Life. Am J Respir Crit Care Med 15:760–770.

28. Xu L, Earl J, Pichichero ME. 2021. Nasopharyngeal microbiome composition associated with *Streptococcus pneumoniae* colonization suggests a protective role of *Corynebacterium* in young children. PLoS One 16:e0257207.

29. Coleman A, Bialasiewicz S, Marsh RL, Grahn Hakansson E, Cottrell K, Wood A, Jayasundara N, Ware RS, Zaugg J, Sidjabat HE, Adams J, Ferguson J, Brown M, Roos K, Cervin A. 2021. Upper Respiratory Microbiota in Relation to Ear and Nose Health Among Australian Aboriginal and Torres Strait Islander Children. J Pediatric Infect Dis Soc 10:468–476.

30. Coleman A, Zaugg J, Wood A, Cottrell K, Hakansson EG, Adams J, Brown M, Cervin A, Bialasiewicz S. 2021. Upper Respiratory Tract Microbiome of Australian Aboriginal and Torres Strait Islander Children in Ear and Nose Health and Disease. Microbiol Spectr 9:e0036721.

31. Escapa IF, Chen T, Huang Y, Gajare P, Dewhirst FE, Lemon KP. 2018. New Insights into Human Nostril Microbiome from the Expanded Human Oral Microbiome Database (eHOMD): a Resource for the Microbiome of the Human Aerodigestive Tract. mSystems 3:e00187–18.

32. Liu CM, Price LB, Hungate BA, Abraham AG, Larsen LA, Christensen K, Stegger M, Skov R, Andersen PS. 2015. *Staphylococcus aureus* and the ecology of the nasal microbiome. Sci Adv 1:e1400216.

33. Teutsch B, Berger A, Marosevic D, Schonberger K, Lam TT, Hubert K, Beer S, Wienert P, Ackermann N, Claus H, Drayss M, Thiel K, van der Linden M, Vogel U, Sing A. 2017. *Corynebacterium* species nasopharyngeal carriage in asymptomatic individuals aged >/= 65 years in Germany. Infection 45:607–611.

34. Salamzade R, Swaney MH, Kalan LR. 2022. Comparative Genomic and Metagenomic Investigations of the *Corynebacterium tuberculostearicum* Species Complex Reveals Potential Mechanisms Underlying Associations To Skin Health and Disease. Microbiol Spectr doi:10.1128/spectrum.03578-22:e0357822.

35. Oh J, Byrd AL, Park M, Program NCS, Kong HH, Segre JA. 2016. Temporal Stability of the Human Skin Microbiome. Cell 165:854–66.

36. Salamzade R, Cheong JZA, Sandstrom S, Swaney MH, Stubbendieck RM, Starr NL, Currie CR, Singh AM, Kalan LR. 2023. Evolutionary investigations of the biosynthetic diversity in the skin microbiome using lsaBGC. Microb Genom 9.

37. Ahmed N, Joglekar P, Deming C, Program NCS, Lemon KP, Kong HH, Segre JA, Conlan S. 2023. Genomic characterization of the C. tuberculostearicum species complex, a prominent member of the human skin microbiome. mSystems 8:e0063223.

38. Brugger SD, Eslami SM, Pettigrew MM, Escapa IF, Henke MT, Kong Y, Lemon KP. 2020. Dolosigranulum pigrum Cooperation and Competition in Human Nasal Microbiota. mSphere 5:e00852–20.

39. Stubbendieck RM, May DS, Chevrette MG, Temkin MI, Wendt-Pienkowski E, Cagnazzo J, Carlson CM, Gern JE, Currie CR. 2019. Competition among Nasal Bacteria Suggests a Role for Siderophore-Mediated Interactions in Shaping the Human Nasal Microbiota. Appl Environ Microbiol 85.

40. Ramsey MM, Freire MO, Gabrilska RA, Rumbaugh KP, Lemon KP. 2016. *Staphylococcus aureus* Shifts toward Commensalism in Response to *Corynebacterium* Species. Frontiers in Microbiology 7:1230.

41. Hardy BL, Dickey SW, Plaut RD, Riggins DP, Stibitz S, Otto M, Merrell DS. 2019. *Corynebacterium pseudodiphtheriticum* Exploits *Staphylococcus aureus* Virulence Components in a Novel Polymicrobial Defense Strategy. mBio 10.

42. Zhao Y, Bitzer A, Power JJ, Belikova D, Torres Salazar BO, Adolf LA, Gerlach D, Krismer B, Heilbronner S. 2024. Nasal commensals reduce Staphylococcus aureus proliferation by restricting siderophore availability. ISME J 18.

43. Uehara Y, Nakama H, Agematsu K, Uchida M, Kawakami Y, Abdul Fattah AS, Maruchi N. 2000. Bacterial interference among nasal inhabitants: eradication of *Staphylococcus aureus* from nasal cavities by artificial implantation of *Corynebacterium* sp. J Hosp Infect 44:127–33.

44. Lina G, Boutite F, Tristan A, Bes M, Etienne J, Vandenesch F. 2003. Bacterial competition for human nasal cavity colonization: role of Staphylococcal agr alleles. Appl Environ Microbiol 69:18–23.

45. Wos-Oxley ML, Plumeier I, von Eiff C, Taudien S, Platzer M, Vilchez-Vargas R, Becker K, Pieper DH. 2010. A poke into the diversity and associations within human anterior nare microbial communities. ISME J 4:839–51.

46. Johnson RC, Ellis MW, Lanier JB, Schlett CD, Cui T, Merrell DS. 2015. Correlation between nasal microbiome composition and remote purulent skin and soft tissue infections. Infection and immunity 83:802–11.

47. Blum FC, Whitmire JM, Bennett JW, Carey PM, Ellis MW, English CE, Law NN, Tribble DR, Millar EV, Merrell DS. 2022. Nasal microbiota evolution within the congregate setting imposed by military training. Sci Rep 12:11492.

48. Kiryukhina NV, Melnikov VG, Suvorov AV, Morozova YA, Ilyin VK. 2013. Use of *Corynebacterium pseudodiphtheriticum* for elimination of *Staphylococcus aureus* from the nasal cavity in volunteers exposed to abnormal microclimate and altered gaseous environment. Probiotics and antimicrobial proteins 5:233–8.

49. Kluytmans J, van Belkum A, Verbrugh H. 1997. Nasal carriage of *Staphylococcus aureus*: epidemiology, underlying mechanisms, and associated risks. Clin Microbiol Rev 10:505–20.

50. von Eiff C, Becker K, Machka K, Stammer H, Peters G, Group FtS. 2001. Nasal carriage as a source of *Staphylococcus aureus* bacteremia. N Engl J Med 344:11–6.

51. Wertheim HF, Vos MC, Ott A, van Belkum A, Voss A, Kluytmans JA, van Keulen PH, Vandenbroucke-Grauls CM, Meester MH, Verbrugh HA. 2004. Risk and outcome of nosocomial *Staphylococcus aureus* bacteraemia in nasal carriers versus non-carriers. Lancet 364:703–5.

52. Young BC, Wu CH, Gordon NC, Cole K, Price JR, Liu E, Sheppard AE, Perera S, Charlesworth J, Golubchik T, Iqbal Z, Bowden R, Massey RC, Paul J, Crook DW, Peto TE, Walker AS, Llewelyn MJ, Wyllie DH, Wilson DJ. 2017. Severe infections emerge from commensal bacteria by adaptive evolution. Elife 6.

53. Kanmani P, Clua P, Vizoso-Pinto MG, Rodriguez C, Alvarez S, Melnikov V, Takahashi H, Kitazawa H, Villena J. 2017. Respiratory Commensal Bacteria *Corynebacterium pseudodiphtheriticum* Improves Resistance of Infant Mice to Respiratory Syncytial Virus and *Streptococcus pneumoniae* Superinfection. Front Microbiol 8:1613.

54. Horn KJ, Jaberi Vivar AC, Arenas V, Andani S, Janoff EN, Clark SE. 2021. *Corynebacterium* Species Inhibit *Streptococcus pneumoniae* Colonization and Infection of the Mouse Airway. Front Microbiol 12:804935.

55. Menberu MA, Liu S, Cooksley C, Hayes AJ, Psaltis AJ, Wormald PJ, Vreugde S. 2021. *Corynebacterium accolens* Has Antimicrobial Activity against *Staphylococcus aureus* and Methicillin-Resistant *S. aureus* Pathogens Isolated from the Sinonasal Niche of Chronic Rhinosinusitis Patients. Pathogens 10.

56. Neubauer M, Šourek J, Rýc M, Boháček J, Mára M, Mňuková J. 1991. *Corynebacterium accolens* sp. nov., a Gram-Positive Rod Exhibiting Satellitism, from Clinical Material. Systematic and Applied Microbiology 14:46–51.

57. Riegel P, de Briel D, Prevost G, Jehl F, Monteil H. 1993. Proposal of *Corynebacterium propinquum* sp. nov. for *Corynebacterium* group ANF-3 strains. FEMS Microbiol Lett 113:229–234.

58. Karlyshev AV, Melnikov VG. 2013. Draft Genome Sequence of *Corynebacterium pseudodiphtheriticum* Strain 090104 "Sokolov". Genome Announc 1.

59. Bernier AM, Bernard K. 2018. Whole-Genome Sequences of *Corynebacterium macginleyi* CCUG 32361(T) and Clinical Isolates NML 080212 and NML 120205. Microbiol Resour Announc 7.

60. Zhang C, Song W, Ma HR, Peng X, Anderson DJ, Fowler VG, Jr., Thaden JT, Xiao M, You L. 2020. Temporal encoding of bacterial identity and traits in growth dynamics. Proc Natl Acad Sci U S A 117:20202–20210.

61. Swaney MH, Sandstrom S, Kalan LR. 2022. Cobamide Sharing Is Predicted in the Human Skin Microbiome. mSystems 7:e0067722.

62. Flores Ramos S, Brugger SD, Escapa IF, Skeete CA, Cotton SL, Eslami SM, Gao W, Bomar L, Tran TH, Jones DS, Minot S, Roberts RJ, Johnston CD, Lemon KP. 2021. Genomic Stability and Genetic Defense Systems in *Dolosigranulum pigrum*, a Candidate Beneficial Bacterium from the Human Microbiome. mSystems 6:e0042521.

63. McInerney JO, McNally A, O’Connell MJ. 2017. Why prokaryotes have pangenomes. Nat Microbiol 2:17040.

64. Eren AM, Kiefl E, Shaiber A, Veseli I, Miller SE, Schechter MS, Fink I, Pan JN, Yousef M, Fogarty EC, Trigodet F, Watson AR, Esen OC, Moore RM, Clayssen Q, Lee MD, Kivenson V, Graham ED, Merrill BD, Karkman A, Blankenberg D, Eppley JM, Sjodin A, Scott JJ, Vazquez-Campos X, McKay LJ, McDaniel EA, Stevens SLR, Anderson RE, Fuessel J, Fernandez-Guerra A, Maignien L, Delmont TO, Willis AD. 2021. Community-led, integrated, reproducible multi-omics with anvi’o. Nat Microbiol 6:3–6.

65. Oliveira PH, Touchon M, Cury J, Rocha EPC. 2017. The chromosomal organization of horizontal gene transfer in bacteria. Nat Commun 8:841.

66. Iranzo J, Wolf YI, Koonin EV, Sela I. 2019. Gene gain and loss push prokaryotes beyond the homologous recombination barrier and accelerate genome sequence divergence. Nat Commun 10:5376.

67. Gautreau G, Bazin A, Gachet M, Planel R, Burlot L, Dubois M, Perrin A, Medigue C, Calteau A, Cruveiller S, Matias C, Ambroise C, Rocha EPC, Vallenet D. 2020. PPanGGOLiN: Depicting microbial diversity via a partitioned pangenome graph. PLoS Comput Biol 16:e1007732.

68. Kalinowski J, Bathe B, Bartels D, Bischoff N, Bott M, Burkovski A, Dusch N, Eggeling L, Eikmanns BJ, Gaigalat L, Goesmann A, Hartmann M, Huthmacher K, Kramer R, Linke B, McHardy AC, Meyer F, Mockel B, Pfefferle W, Puhler A, Rey DA, Ruckert C, Rupp O, Sahm H, Wendisch VF, Wiegrabe I, Tauch A. 2003. The complete *Corynebacterium glutamicum* ATCC 13032 genome sequence and its impact on the production of L-aspartate-derived amino acids and vitamins. J Biotechnol 104:5–25.

69. Wolf S, Becker J, Tsuge Y, Kawaguchi H, Kondo A, Marienhagen J, Bott M, Wendisch VF, Wittmann C. 2021. Advances in metabolic engineering of Corynebacterium glutamicum to produce high-value active ingredients for food, feed, human health, and well-being. Essays Biochem 65:197–212.

70. Oliveira A, Oliveira LC, Aburjaile F, Benevides L, Tiwari S, Jamal SB, Silva A, Figueiredo HCP, Ghosh P, Portela RW, De Carvalho Azevedo VA, Wattam AR. 2017. Insight of Genus *Corynebacterium*: Ascertaining the Role of Pathogenic and Non-pathogenic Species. Front Microbiol 8:1937.

71. Burkovski A. 2013. Cell envelope of corynebacteria: structure and influence on pathogenicity. ISRN Microbiol 2013:935736.

72. Cronan JE. 2016. Assembly of Lipoic Acid on Its Cognate Enzymes: an Extraordinary and Essential Biosynthetic Pathway. Microbiol Mol Biol Rev 80:429–50.

73. Tramonti A, Nardella C, di Salvo ML, Barile A, D’Alessio F, de Crecy-Lagard V, Contestabile R. 2021. Knowns and Unknowns of Vitamin B(6) Metabolism in Escherichia coli. EcoSal Plus 9.

74. Shelton AN, Seth EC, Mok KC, Han AW, Jackson SN, Haft DR, Taga ME. 2019. Uneven distribution of cobamide biosynthesis and dependence in bacteria predicted by comparative genomics. ISME J 13:789–804.

75. Bruggemann H, Henne A, Hoster F, Liesegang H, Wiezer A, Strittmatter A, Hujer S, Durre P, Gottschalk G. 2004. The complete genome sequence of *Propionibacterium acnes*, a commensal of human skin. Science 305:671–3.

76. Tettelin H, Nelson KE, Paulsen IT, Eisen JA, Read TD, Peterson S, Heidelberg J, DeBoy RT, Haft DH, Dodson RJ, Durkin AS, Gwinn M, Kolonay JF, Nelson WC, Peterson JD, Umayam LA, White O, Salzberg SL, Lewis MR, Radune D, Holtzapple E, Khouri H, Wolf AM, Utterback TR, Hansen CL, McDonald LA, Feldblyum TV, Angiuoli S, Dickinson T, Hickey EK, Holt IE, Loftus BJ, Yang F, Smith HO, Venter JC, Dougherty BA, Morrison DA, Hollingshead SK, Fraser CM. 2001. Complete genome sequence of a virulent isolate of *Streptococcus pneumoniae*. Science 293:498–506.

77. Diep BA, Gill SR, Chang RF, Phan TH, Chen JH, Davidson MG, Lin F, Lin J, Carleton HA, Mongodin EF, Sensabaugh GF, Perdreau-Remington F. 2006. Complete genome sequence of USA300, an epidemic clone of community-acquired meticillin-resistant *Staphylococcus aureus*. Lancet 367:731–9.

78. Gill SR, Fouts DE, Archer GL, Mongodin EF, Deboy RT, Ravel J, Paulsen IT, Kolonay JF, Brinkac L, Beanan M, Dodson RJ, Daugherty SC, Madupu R, Angiuoli SV, Durkin AS, Haft DH, Vamathevan J, Khouri H, Utterback T, Lee C, Dimitrov G, Jiang L, Qin H, Weidman J, Tran K, Kang K, Hance IR, Nelson KE, Fraser CM. 2005. Insights on evolution of virulence and resistance from the complete genome analysis of an early methicillin-resistant *Staphylococcus aureus* strain and a biofilm-producing methicillin-resistant *Staphylococcus epidermidis* strain. J Bacteriol 187:2426–38.

79. Seibold G, Dempf S, Schreiner J, Eikmanns BJ. 2007. Glycogen formation in Corynebacterium glutamicum and role of ADP-glucose pyrophosphorylase. Microbiology (Reading) 153:1275–1285.

80. Seibold GM, Eikmanns BJ. 2007. The *glgX* gene product of *Corynebacterium glutamicum* is required for glycogen degradation and for fast adaptation to hyperosmotic stress. Microbiology (Reading) 153:2212–2220.

81. Tsai YC, Conlan S, Deming C, Program NCS, Segre JA, Kong HH, Korlach J, Oh J. 2016. Resolving the Complexity of Human Skin Metagenomes Using Single-Molecule Sequencing. mBio 7:e01948–15.

82. Tauch A, Schneider J, Szczepanowski R, Tilker A, Viehoever P, Gartemann KH, Arnold W, Blom J, Brinkrolf K, Brune I, Gotker S, Weisshaar B, Goesmann A, Droge M, Puhler A. 2008. Ultrafast pyrosequencing of *Corynebacterium kroppenstedtii* DSM44385 revealed insights into the physiology of a lipophilic corynebacterium that lacks mycolic acids. J Biotechnol 136:22–30.

83. Sichtig H, Minogue T, Yan Y, Stefan C, Hall A, Tallon L, Sadzewicz L, Nadendla S, Klimke W, Hatcher E, Shumway M, Aldea DL, Allen J, Koehler J, Slezak T, Lovell S, Schoepp R, Scherf U. 2019. FDA-ARGOS is a database with public quality-controlled reference genomes for diagnostic use and regulatory science. Nat Commun 10:3313.

84. Marienhagen J, Kennerknecht N, Sahm H, Eggeling L. 2005. Functional analysis of all aminotransferase proteins inferred from the genome sequence of Corynebacterium glutamicum. J Bacteriol 187:7639–46.

85. McHardy AC, Tauch A, Ruckert C, Puhler A, Kalinowski J. 2003. Genome-based analysis of biosynthetic aminotransferase genes of Corynebacterium glutamicum. J Biotechnol 104:229–40.

86. Koper K, Han SW, Pastor DC, Yoshikuni Y, Maeda HA. 2022. Evolutionary origin and functional diversification of aminotransferases. J Biol Chem 298:102122.

87. Gelfand DH, Steinberg RA. 1977. Escherichia coli mutants deficient in the aspartate and aromatic amino acid aminotransferases. J Bacteriol 130:429–40.

88. Lensmire JM, Hammer ND. 2019. Nutrient sulfur acquisition strategies employed by bacterial pathogens. Curr Opin Microbiol 47:52–58.

89. Ruckert C, Koch DJ, Rey DA, Albersmeier A, Mormann S, Puhler A, Kalinowski J. 2005. Functional genomics and expression analysis of the *Corynebacterium glutamicum fpr2-cysIXHDNYZ* gene cluster involved in assimilatory sulphate reduction. BMC Genomics 6:121.

90. Williams SJ, Senaratne RH, Mougous JD, Riley LW, Bertozzi CR. 2002. 5’-adenosinephosphosulfate lies at a metabolic branch point in mycobacteria. J Biol Chem 277:32606–15.

91. Berndt C, Lillig CH, Wollenberg M, Bill E, Mansilla MC, de Mendoza D, Seidler A, Schwenn JD. 2004. Characterization and reconstitution of a 4Fe-4S adenylyl sulfate/phosphoadenylyl sulfate reductase from Bacillus subtilis. J Biol Chem 279:7850–5.

92. Simic P, Willuhn J, Sahm H, Eggeling L. 2002. Identification of glyA (encoding serine hydroxymethyltransferase) and its use together with the exporter ThrE to increase L-threonine accumulation by Corynebacterium glutamicum. Appl Environ Microbiol 68:3321–7.

93. Tesch M, de Graaf AA, Sahm H. 1999. In vivo fluxes in the ammonium-assimilatory pathways in corynebacterium glutamicum studied by 15N nuclear magnetic resonance. Appl Environ Microbiol 65:1099–109.

94. Beckers G, Nolden L, Burkovski A. 2001. Glutamate synthase of Corynebacterium glutamicum is not essential for glutamate synthesis and is regulated by the nitrogen status. Microbiology (Reading) 147:2961–70.

95. Son HF, Kim KJ. 2016. Structural Insights into a Novel Class of Aspartate Aminotransferase from *Corynebacterium glutamicum*. PLoS One 11:e0158402.

96. Hirasawa T, Wachi M, Nagai K. 2000. A mutation in the Corynebacterium glutamicum ltsA gene causes susceptibility to lysozyme, temperature-sensitive growth, and L-glutamate production. J Bacteriol 182:2696–701.

97. Javid B, Sorrentino F, Toosky M, Zheng W, Pinkham JT, Jain N, Pan M, Deighan P, Rubin EJ. 2014. Mycobacterial mistranslation is necessary and sufficient for rifampicin phenotypic resistance. Proc Natl Acad Sci U S A 111:1132–7.

98. Agapova A, Serafini A, Petridis M, Hunt DM, Garza-Garcia A, Sohaskey CD, de Carvalho LPS. 2019. Flexible nitrogen utilisation by the metabolic generalist pathogen Mycobacterium tuberculosis. Elife 8.

99. Marienhagen J, Eggeling L. 2008. Metabolic function of Corynebacterium glutamicum aminotransferases AlaT and AvtA and impact on L-valine production. Appl Environ Microbiol 74:7457–62.

100. Copeland E, Leonard K, Carney R, Kong J, Forer M, Naidoo Y, Oliver BGG, Seymour JR, Woodcock S, Burke CM, Stow NW. 2018. Chronic Rhinosinusitis: Potential Role of Microbial Dysbiosis and Recommendations for Sampling Sites. Front Cell Infect Microbiol 8:57.

101. Renz A, Widerspick L, Drager A. 2021. First Genome-Scale Metabolic Model of Dolosigranulum pigrum Confirms Multiple Auxotrophies. Metabolites 11.

102. Adolf LA, Heilbronner S. 2022. Nutritional Interactions between Bacterial Species Colonising the Human Nasal Cavity: Current Knowledge and Future Prospects. Metabolites 12.

103. Jang J, Forbes VE, Sadowsky MJ. 2022. Probable role of *Cutibacterium acnes* in the gut of the polychaete *Capitella teleta*. Sci Total Environ 809:151127.

104. Howden BP, Giulieri SG, Wong Fok Lung T, Baines SL, Sharkey LK, Lee JYH, Hachani A, Monk IR, Stinear TP. 2023. *Staphylococcus aureus* host interactions and adaptation. Nat Rev Microbiol doi:10.1038/s41579-023-00852-y:1-16.

105. Blasche S, Kim Y, Patil KR. 2017. Draft Genome Sequence of *Corynebacterium kefirresidentii* SB, Isolated from Kefir. Genome Announc 5.

106. Chen T, Yu WH, Izard J, Baranova OV, Lakshmanan A, Dewhirst FE. 2010. The Human Oral Microbiome Database: a web accessible resource for investigating oral microbe taxonomic and genomic information. Database (Oxford) 2010:baq013.

107. Saheb Kashaf S, Proctor DM, Deming C, Saary P, Holzer M, Program NCS, Taylor ME, Kong HH, Segre JA, Almeida A, Finn RD. 2022. Integrating cultivation and metagenomics for a multi-kingdom view of skin microbiome diversity and functions. Nat Microbiol 7:169–179.

108. Nurk S, Bankevich A, Antipov D, Gurevich AA, Korobeynikov A, Lapidus A, Prjibelski AD, Pyshkin A, Sirotkin A, Sirotkin Y, Stepanauskas R, Clingenpeel SR, Woyke T, McLean JS, Lasken R, Tesler G, Alekseyev MA, Pevzner PA. 2013. Assembling single-cell genomes and mini-metagenomes from chimeric MDA products. J Comput Biol 20:714–37.

109. Walker BJ, Abeel T, Shea T, Priest M, Abouelliel A, Sakthikumar S, Cuomo CA, Zeng Q, Wortman J, Young SK, Earl AM. 2014. Pilon: an integrated tool for comprehensive microbial variant detection and genome assembly improvement. PLoS One 9:e112963.

110. Bolger AM, Lohse M, Usadel B. 2014. Trimmomatic: a flexible trimmer for Illumina sequence data. Bioinformatics 30:2114–20.

111. Bankevich A, Nurk S, Antipov D, Gurevich AA, Dvorkin M, Kulikov AS, Lesin VM, Nikolenko SI, Pham S, Prjibelski AD, Pyshkin AV, Sirotkin AV, Vyahhi N, Tesler G, Alekseyev MA, Pevzner PA. 2012. SPAdes: A New Genome Assembly Algorithm and Its Applications to Single-Cell Sequencing. Journal of Computational Biology 19:455–477.

112. Parks DH, Imelfort M, Skennerton CT, Hugenholtz P, Tyson GW. 2015. CheckM: assessing the quality of microbial genomes recovered from isolates, single cells, and metagenomes. Genome Res 25:1043–55.

113. Human Microbiome Jumpstart Reference Strains C, Nelson KE, Weinstock GM, Highlander SK, Worley KC, Creasy HH, Wortman JR, Rusch DB, Mitreva M, Sodergren E, Chinwalla AT, Feldgarden M, Gevers D, Haas BJ, Madupu R, Ward DV, Birren BW, Gibbs RA, Methe B, Petrosino JF, Strausberg RL, Sutton GG, White OR, Wilson RK, Durkin S, Giglio MG, Gujja S, Howarth C, Kodira CD, Kyrpides N, Mehta T, Muzny DM, Pearson M, Pepin K, Pati A, Qin X, Yandava C, Zeng Q, Zhang L, Berlin AM, Chen L, Hepburn TA, Johnson J, McCorrison J, Miller J, Minx P, Nusbaum C, Russ C, Sykes SM, Tomlinson CM, et al. 2010. A catalog of reference genomes from the human microbiome. Science 328:994–9.

114. Perrin A, Rocha EPC. 2021. PanACoTA: a modular tool for massive microbial comparative genomics. NAR Genom Bioinform 3:lqaa106.

115. Seemann T. 2014. Prokka: rapid prokaryotic genome annotation. Bioinformatics 30:2068–9.

116. Hyatt D, Chen G-L, Locascio PF, Land ML, Larimer FW, Hauser LJ. 2010. Prodigal: prokaryotic gene recognition and translation initiation site identification. BMC Bioinformatics 11:119.

117. Contreras-Moreira B, Vinuesa P. 2013. GET_HOMOLOGUES, a versatile software package for scalable and robust microbial pangenome analysis. Appl Environ Microbiol 79:7696–701.

118. Vinuesa P, Contreras-Moreira B. 2015. Robust identification of orthologues and paralogues for microbial pan-genomics using GET_HOMOLOGUES: a case study of pIncA/C plasmids. Methods Mol Biol 1231:203–32.

119. Kristensen DM, Kannan L, Coleman MK, Wolf YI, Sorokin A, Koonin EV, Mushegian A. 2010. A low-polynomial algorithm for assembling clusters of orthologous groups from intergenomic symmetric best matches. Bioinformatics 26:1481–1487.

120. Li L. 2003. OrthoMCL: Identification of Ortholog Groups for Eukaryotic Genomes. Genome Research 13:2178–2189.

121. Altschul S. 1997. Gapped BLAST and PSI-BLAST: a new generation of protein database search programs. Nucleic Acids Research 25:3389–3402.

122. Vinuesa P, Ochoa-Sanchez LE, Contreras-Moreira B. 2018. GET_PHYLOMARKERS, a Software Package to Select Optimal Orthologous Clusters for Phylogenomics and Inferring Pan-Genome Phylogenies, Used for a Critical Geno-Taxonomic Revision of the Genus Stenotrophomonas. Front Microbiol 9:771.

123. Minh BQ, Schmidt HA, Chernomor O, Schrempf D, Woodhams MD, von Haeseler A, Lanfear R. 2020. IQ-TREE 2: New Models and Efficient Methods for Phylogenetic Inference in the Genomic Era. Mol Biol Evol 37:1530–1534.

124. Chernomor O, von Haeseler A, Minh BQ. 2016. Terrace Aware Data Structure for Phylogenomic Inference from Supermatrices. Syst Biol 65:997–1008.

125. Hoang DT, Chernomor O, von Haeseler A, Minh BQ, Vinh LS. 2018. UFBoot2: Improving the Ultrafast Bootstrap Approximation. Mol Biol Evol 35:518–522.

126. Kalyaanamoorthy S, Minh BQ, Wong TKF, Von Haeseler A, Jermiin LS. 2017. ModelFinder: fast model selection for accurate phylogenetic estimates. Nature Methods 14:587–589.

127. Letunic I, Bork P. 2021. Interactive Tree Of Life (iTOL) v5: an online tool for phylogenetic tree display and annotation. Nucleic Acids Res 49:W293–W296.

128. Delmont TO, Eren AM. 2018. Linking pangenomes and metagenomes: the *Prochlorococcus* metapangenome. PeerJ 6:e4320.

129. Buchfink B, Xie C, Huson DH. 2015. Fast and sensitive protein alignment using DIAMOND. Nat Methods 12:59–60.

130. Tatusov RL, Natale DA, Garkavtsev IV, Tatusova TA, Shankavaram UT, Rao BS, Kiryutin B, Galperin MY, Fedorova ND, Koonin EV. 2001. The COG database: new developments in phylogenetic classification of proteins from complete genomes. Nucleic Acids Res 29:22–8.

131. Galperin MY, Wolf YI, Makarova KS, Vera Alvarez R, Landsman D, Koonin EV. 2021. COG database update: focus on microbial diversity, model organisms, and widespread pathogens. Nucleic Acids Res 49:D274–D281.

132. Kanehisa M, Goto S. 2000. KEGG: kyoto encyclopedia of genes and genomes. Nucleic Acids Res 28:27–30.

133. Kanehisa M, Furumichi M, Sato Y, Kawashima M, Ishiguro-Watanabe M. 2023. KEGG for taxonomy-based analysis of pathways and genomes. Nucleic Acids Res 51:D587–D592.

134. Bateman A, Birney E, Durbin R, Eddy SR, Howe KL, Sonnhammer EL. 2000. The Pfam protein families database. Nucleic Acids Res 28:263–6.

135. Eddy SR. 2011. Accelerated Profile HMM Searches. PLoS Comput Biol 7:e1002195.

136. Altschul SF, Gish W, Miller W, Myers EW, Lipman DJ. 1990. Basic local alignment search tool. J Mol Biol 215:403–10.

137. Edgar RC. 2004. MUSCLE: multiple sequence alignment with high accuracy and high throughput. Nucleic Acids Res 32:1792–7.

138. Benedict MN, Henriksen JR, Metcalf WW, Whitaker RJ, Price ND. 2014. ITEP: an integrated toolkit for exploration of microbial pan-genomes. BMC Genomics 15:8.

139. van Dongen S, Abreu-Goodger C. 2012. Using MCL to Extract Clusters from Networks. *In* van Helden J, Toussaint A, Thieffry D (ed), Bacterial Molecular Networks Methods in Molecular Biology (Methods and Protocols), vol 804. Springer, New York, NY.

140. Veseli I, Chen YT, Schechter MS, Vanni C, Fogarty EC, Watson AR, Jabri B, Blekhman R, Willis AD, Yu MK, Fernandez-Guerra A, Fussel J, Eren AM. 2023. Microbes with higher metabolic independence are enriched in human gut microbiomes under stress. bioRxiv doi:10.1101/2023.05.10.540289.

141. Shaiber A, Willis AD, Delmont TO, Roux S, Chen LX, Schmid AC, Yousef M, Watson AR, Lolans K, Esen OC, Lee STM, Downey N, Morrison HG, Dewhirst FE, Mark Welch JL, Eren AM. 2020. Functional and genetic markers of niche partitioning among enigmatic members of the human oral microbiome. Genome Biol 21:292.

142. Bates D, Mächler M, Bolker B, Walker S. 2015. Fitting Linear Mixed-Effects Models Using lme4. Journal of Statistical Software 67:1–48.

143. Quinlan AR, Hall IM. 2010. BEDTools: a flexible suite of utilities for comparing genomic features. Bioinformatics 26:841–2.

144. Quinlan AR. 2014. BEDTools: The Swiss-Army Tool for Genome Feature Analysis. Curr Protoc Bioinformatics 47:11 12 1–34.

145. Katoh K, Misawa K, Kuma K, Miyata T. 2002. MAFFT: a novel method for rapid multiple sequence alignment based on fast Fourier transform. Nucleic Acids Res 30:3059–66.

146. Katoh K, Standley DM. 2013. MAFFT Multiple Sequence Alignment Software Version 7: Improvements in Performance and Usability. Molecular Biology and Evolution 30:772–780.

147. Larsson A. 2014. AliView: a fast and lightweight alignment viewer and editor for large datasets. Bioinformatics 30:3276–8.

148. Koch DJ, Ruckert C, Rey DA, Mix A, Puhler A, Kalinowski J. 2005. Role of the ssu and seu genes of Corynebacterium glutamicum ATCC 13032 in utilization of sulfonates and sulfonate esters as sulfur sources. Appl Environ Microbiol 71:6104–14.

149. Rehm N, Georgi T, Hiery E, Degner U, Schmiedl A, Burkovski A, Bott M. 2010. L-Glutamine as a nitrogen source for Corynebacterium glutamicum: derepression of the AmtR regulon and implications for nitrogen sensing. Microbiology (Reading) 156:3180–3193.

150. Buerger J, Rehm N, Grebenstein L, Burkovski A. 2016. Glutamine metabolism of Corynebacterium glutamicum: role of the glutaminase GlsK. FEMS Microbiol Lett 363.

151. Wang Q, Jiang A, Tang J, Gao H, Zhang X, Yang T, Xu Z, Xu M, Rao Z. 2021. Enhanced production of L-arginine by improving carbamoyl phosphate supply in metabolically engineered Corynebacterium crenatum. Appl Microbiol Biotechnol 105:3265–3276.

