## Supplementary material for "Metabolic capabilities are highly conserved among human nasal-associated *Corynebacterium* species in pangenomic analyses": Figure S1

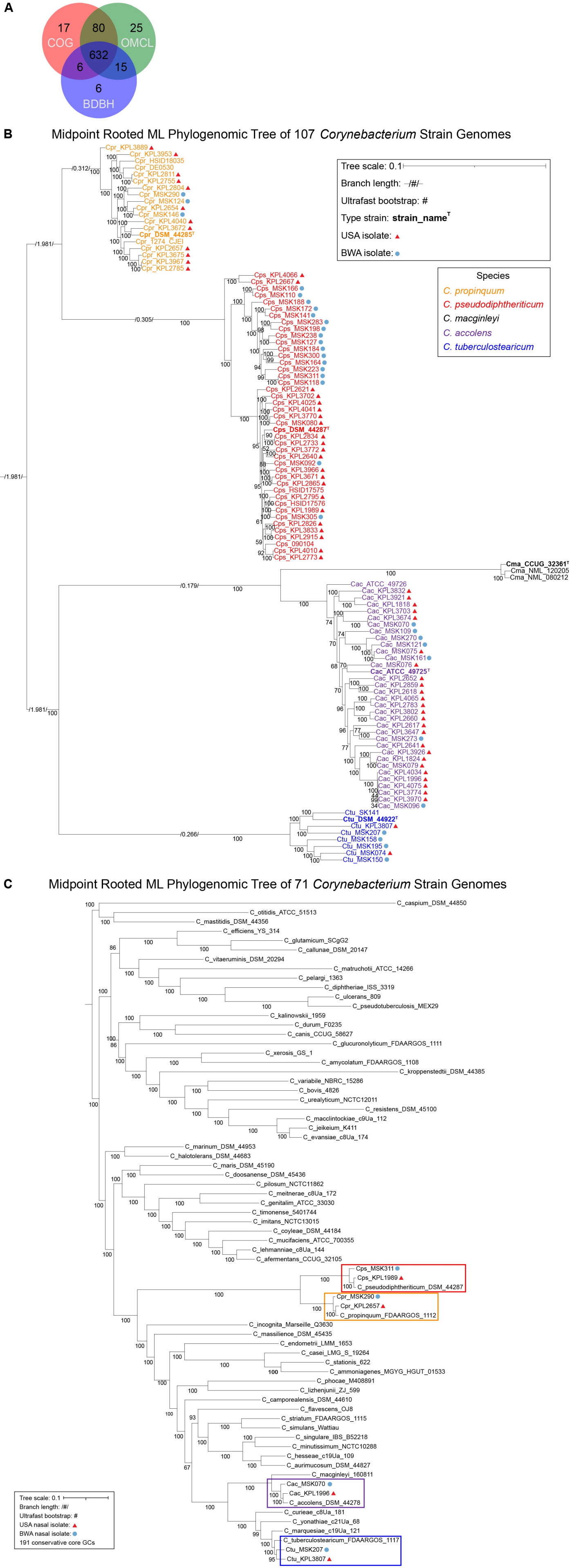

A
COG
OMCL
BDBH
17
80
25
632
6
15
6
B
Midpoint Rooted ML Phylogenomic Tree of 107 Corynebacterium Strain Genomes
Tree scale: 0.1
Branch length: /#/
Ultrafast bootstrap: #
Type strain: strain_name^T
USA isolate:
BWA isolate:
Species
C. propinquum
C. pseudodiphtheriticum
C. macginleyi
C. accolens
C. tuberculostearicum
Cpr_KPL3889
Cpr_KPL3953
Cpr_HSID18035
Cpr_DE0530
Cpr_KPL2811
Cpr_KPL2755
Cpr_KPL2804
Cpr_MSK290
Cpr_MSK124
Cpr_KPL2654
Cpr_MSK146
Cpr_KPL4040
Cpr_KPL3672
Cpr_DSM_44285^T
Cpr_1274_CJEI
Cpr_KPL2657
Cpr_KPL3675
Cpr_KPL3967
Cpr_KPL2785
Cps_KPL4066
Cps_KPL2667
Cps_MSK166
Cps_MSK110
Cps_MSK188
Cps_MSK172
Cps_MSK141
Cps_MSK283
Cps_MSK198
Cps_MSK238
Cps_MSK127
Cps_MSK184
Cps_MSK300
Cps_MSK164
Cps_MSK223
Cps_MSK311
Cps_MSK118
Cps_KPL2621
Cps_KPL3702
Cps_KPL4025
Cps_KPL4041
Cps_KPL3770
Cps_MSK080
Cps_DSM_44287^T
Cps_KPL2834
Cps_KPL2733
Cps_KPL3772
Cps_KPL2640
Cps_MSK092
Cps_KPL3966
Cps_KPL3671
Cps_KPL2865
Cps_HSID17575
Cps_KPL2795
Cps_HSID17576
Cps_KPL1989
Cps_MSK305
Cps_KPL2826
Cps_KPL3833
Cps_KPL2915
Cps_090104
Cps_KPL4010
Cps_KPL2773
Cma_CCUG_32361^T
Cma_NML_120205
Cma_NML_080212
Cac_ATCC_49726
Cac_KPL3832
Cac_KPL3921
Cac_KPL1818
Cac_KPL3703
Cac_KPL3674
Cac_MSK070
Cac_MSK109
Cac_MSK270
Cac_MSK121
Cac_MSK075
Cac_MSK161
Cac_MSK076
Cac_ATCC_49725^T
Cac_KPL2652
Cac_KPL2859
Cac_KPL2618
Cac_KPL4065
Cac_KPL2783
Cac_KPL3802
Cac_KPL2660
Cac_KPL2617
Cac_KPL3647
Cac_MSK273
Cac_KPL2641
Cac_KPL3926
Cac_KPL1824
Cac_MSK079
Cac_KPL4034
Cac_KPL1996
Cac_KPL4075
Cac_KPL3774
Cac_KPL3970
Cac_MSK096
Ctu_SK141
Ctu_DSM_44922^T
Ctu_KPL3807
Ctu_MSK207
Ctu_MSK158
Ctu_MSK195
Ctu_MSK074
Ctu_MSK150
C
Midpoint Rooted ML Phylogenomic Tree of 71 Corynebacterium Strain Genomes
C_caspium_DSM_44850
C_otitidis_ATCC_51513
C_mastitidis_DSM_44356
C_efficiens_YS_314
C_glutamicum_SCgG2
C_callunae_DSM_20147
C_vitaeruminis_DSM_20294
C_matruchotii_ATCC_14266
C_pelargi_1363
C_diphtheriae_ISS_3319
C_ulcerans_809
C_pseudotuberculosis_MEX29
C_kalinowskii_1959
C_durum_F0235
C_canis_CCUG_58627
C_glucuronolyticum_FDAARGOS_1111
C_xerosis_GS_1
C_amycolatum_FDAARGOS_1108
C_kroppenstedtii_DSM_44385
C_variabile_NBRC_15286
C_bovis_4826
C_urealyticum_NCTC12011
C_resistens_DSM_45100
C_macclintockiae_c9Ua_112
C_jeikeium_K411
C_evansiae_c8Ua_174
C_marinum_DSM_44953
C_halotolerans_DSM_44683
C_maris_DSM_45190
C_doosanense_DSM_45436
C_pilosum_NCTC11862
C_meitnerae_c8Ua_172
C_genitalim_ATCC_33030
C_timonense_5401744
C_imitans_NCTC13015
C_coyleae_DSM_44184
C_mucifaciens_ATCC_700355
C_lehmanniae_c8Ua_144
C_afermentans_CCUG_32105
Cps_MSK311
Cps_KPL1989
C_pseudodiphtheriticum_DSM_44287
Cpr_MSK290
Cpr_KPL2657
C_propinquum_FDAARGOS_1112
C_incognita_Marseille_Q3630
C_massilience_DSM_45435
C_endometrii_LMM_1653
C_casei_LMG_S_19264
C_stationis_622
C_ammoniagenes_MGYG_HGUT_01533
C_phocae_M408891
C_lizhenjunii_ZJ_599
C_camporealensis_DSM_44610
C_flavescens_OJ8
C_striatum_FDAARGOS_1115
C_simulans_Wattiau
C_singulare_IBS_B52218
C_minutissimum_NCTC10288
C_hesseae_c19Ua_109
C_aurimucosum_DSM_44827
C_macginleyi_160811
Cac_MSK070
Cac_KPL1996
C_accolens_DSM_44278
C_curieae_c8Ua_181
C_yonathiae_c21Ua_68
C_marquesiae_c19Ua_121
C_tuberculostearicum_FDAARGOS_1117
Ctu_MSK207
Ctu_KPL3807
Tree scale: 0.1
Branch length: /#/
Ultrafast bootstrap: #
USA nasal isolate:
BWA nasal isolate:
191 conservative core GCs

**Figure S1. Phylogenomic trees, generated from the shared conservative core gene clusters, illustrate the evolutionary relationships among four human-nasal-associated *Corynebacterium* species and their relationship to other species within this genus.** (**A**) Venn diagram showing 632 single-copy GCs are shared between 107 *Corynebacterium* strain genomes based on the consensus of BDBH, COG triangle, and OMCL algorithms using GET_HOMOLOGUES v24082022. (**B**) A maximum-likelihood phylogenomic tree constructed from 632 concatenated shared single-copy GCs from 107 *Corynebacterium* strain genomes. Species are indicated by name and color-code (see key) with type strains highlighted in bold. We used IQ-Tree v2.1.3 with model finder (BIC value 11679476.6345), edge-linked-proportional partition model, and 1,000 ultrafast rapid bootstraps (values on tree). Branch lengths are indicated (see key). The phylogeny contains 31 and 56 nasal-isolated strain genomes from Botswana (blue circles) and the USA (red triangles). Many branches are highly supported with ultrafast bootstrap values ≥ 95. (**C**) A maximum-likelihood phylogenomic tree of 63 *Corynebacterium* species illustrates the relationship of the four nasal species to other members of the genus. *C. propinquum*, *C. pseudodiphtheriticum*, *C. accolens*, and *C. tuberculostearicum* are each highlighted by a colored box. This is based on 191 shared conservative core GCs from 71 *Corynebacterium* strain genomes. Representative nasal strain genomes from Botswana (blue circle) and the USA (red triangle) are shown for each of the four species.
