## Supplementary material for "Metabolic capabilities are highly conserved among human nasal-associated *Corynebacterium* species in pangenomic analyses": Figure S2

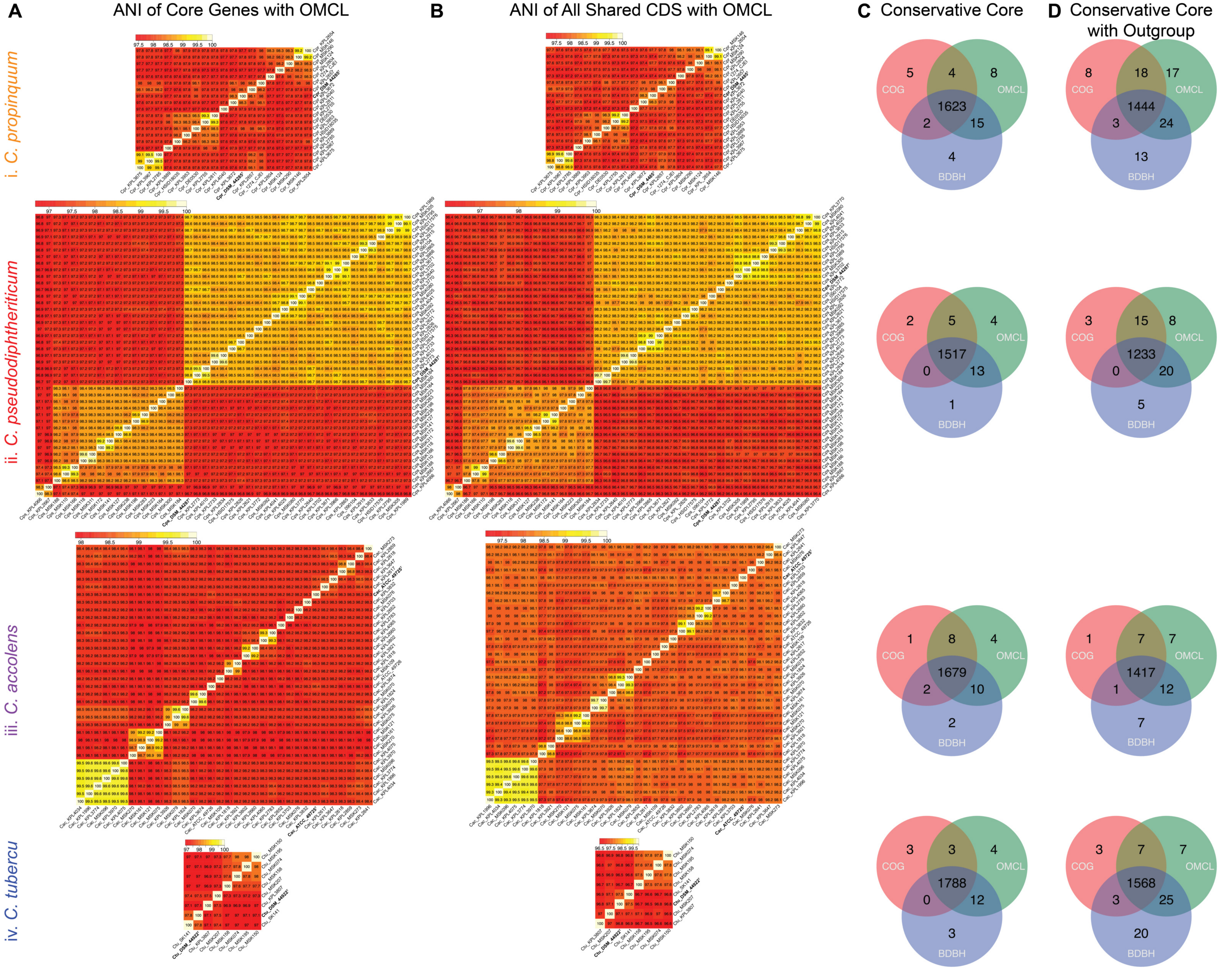
A
ANI of Core Genes with OMCL
B
ANI of All Shared CDS with OMCL
C Conservative Core
D Conservative Core with Outgroup
i. C. propinquum
ii. C. pseudodiphtheriticum
iii. C. accolens
iv. C. tubercu
COG
OMCL
BDBH
5
4
8
1623
2
15
4
8
18
17
1444
3
24
13
2
5
4
1517
0
13
1
3
15
8
1233
0
20
5
1
8
4
1679
2
10
2
1
7
7
1417
1
12
7
3
3
4
1788
0
12
3
3
7
7
1568
3
25
20

**Figure S2. Species-level pairwise average nucleotide identities and conservative core GCs. (A**) The shared core genes among all paired strain genomes within each species (**i-iv**) have ≥ 95% ANI (average nucleotide identity) to the type strain using the OMCL algorithm. (**B**) The ANI pairwise comparisons for all shared CDS regions are slightly less than core ANI values but still ≥ 95% among strain genomes within each species. The order of strains in A and B differ due to changes in pairwise identities because more SNPs were included in B. (**C**) The shared conservative core single-copy GCs determined by the consensus of BDBH, COG triangle, and OMCL for each of the *Corynebacterium* species. (**D**) The conservative core CGs for each species plus the type strain of its closest relative in the five-species phylogenomic tree (**Fig. S1B**). Between 112 and 240 GCs were absent from the conservative core of the outgroup for each species, with the fewest absent in the outgroup for *C. pseudodiphtheriticum* and the most absent in the outgroup for *C. tuberculostearicum*.
