## Supplementary material for "Metabolic capabilities are highly conserved among human nasal-associated *Corynebacterium* species in pangenomic analyses": Figure S3

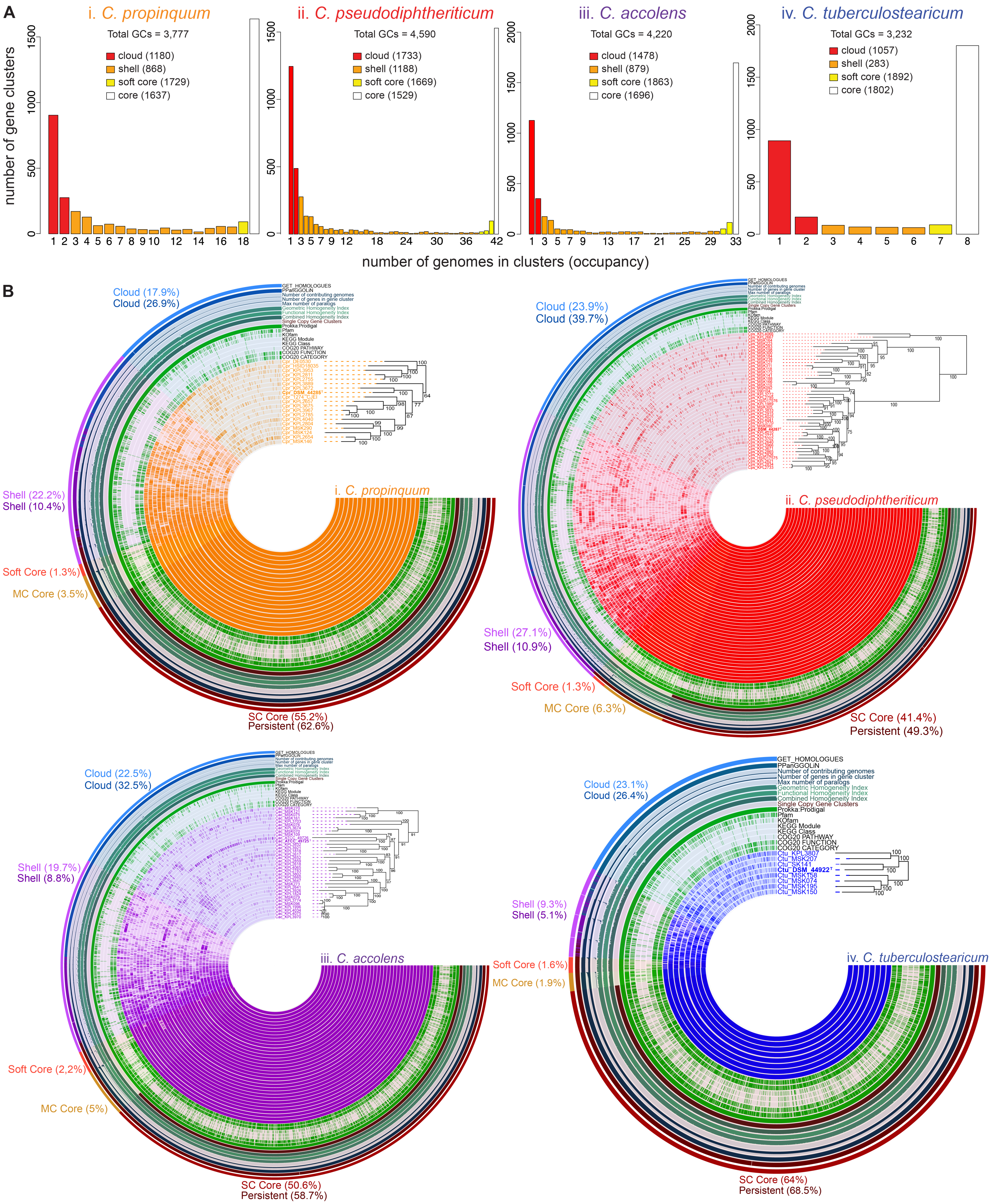
A
i. C. propinquum
Total GCs = 3,777
cloud (1180)
shell (868)
soft core (1729)
core (1637)
ii. C. pseudodiphtheriticum
Total GCs = 4,590
cloud (1733)
shell (1188)
soft core (1669)
core (1529)
iii. C. accolens
Total GCs = 4,220
cloud (1478)
shell (879)
soft core (1863)
core (1696)
iv. C. tuberculostearicum
Total GCs = 3,232
cloud (1057)
shell (283)
soft core (1892)
core (1802)
number of gene clusters
number of genomes in clusters (occupancy)
B
Cloud (17.9%)
Cloud (26.9%)
Shell (22.2%)
Shell (10.4%)
Soft Core (1.3%)
MC Core (3.5%)
SC Core (55.2%)
Persistent (62.6%)
i. C. propinquum
Cloud (23.9%)
Cloud (39.7%)
Shell (27.1%)
Shell (10.9%)
Soft Core (1.3%)
MC Core (6.3%)
SC Core (41.4%)
Persistent (49.3%)
ii. C. pseudodiphtheriticum
Cloud (22.5%)
Cloud (32.5%)
Shell (19.7%)
Shell (8.8%)
Soft Core (2,2%)
MC Core (5%)
SC Core (50.6%)
Persistent (58.7%)
iii. C. accolens
Cloud (23.1%)
Cloud (26.4%)
Shell (9.3%)
Shell (5.1%)
Soft Core (1.6%)
MC Core (1.9%)
SC Core (64%)
Persistent (68.5%)
iv. C. tuberculostearicum
GET_HOMOLOGUES
PPanGGOLiN
Number of contributing genomes
Number of genes in gene cluster
Max number of paralogs
Geometric Homogeneity Index
Functional Homogeneity Index
Combined Homogeneity Index
Single Copy Gene Clusters
Prokka:Prodigal
Pfam
KOfam
KEGG Module
KEGG Class
COG20 PATHWAY
COG20 FUNCTION
COG20 CATEGORY
Ctu_KPL3807
Ctu_MSK207
Ctu_SK141
Ctu_DSM_44922ᵀ
Ctu_MSK158
Ctu_MSK074
Ctu_MSK195
Ctu_MSK150

**Figure S3. Three different analysis platforms estimated a similar pangenome for each individual *Corynebacterium* species.** (**A**) Representation of the pangenome estimated by GET_HOMOLOGUES with COG triangle and OMCL of (**i**) *C. propinquum*, 19 genomes; (**ii)** *C. pseudodiphtheriticum*, 42 genomes; (**iii**) *C. accolens*, 33 genomes; and (**iv**) *C. tuberculostearicum,* 8 genomes. (**B**) Using pangenomes clustered with anvi'o, we constructed a concentric dendrogram for each species (**i-vi**) ordered to match the species-specific phylogenies in Figure 1. The GCs present in the core/persistent genome consistently accounted for over half of the pangenome in each of the nasal *Corynebacterium* species. The outermost concentric ring indicates anvi'o-defined GCs (with percentages) in the single copy core (dark red), multicopy core (beige), soft core (red), shell (light purple), and cloud (light blue) based on GET_HOMOLOGUES definition of each partition with anvi'o allowing further division of the core into single copy and multicopy core. The next to outermost ring indicates anvi'o-defined GCs (with percentages) assigned by PPanGGOLiN to persistent (dark maroon), shell (purple), and cloud (blue). The average core+soft core or persistent, shell, and cloud sizes across the four species using GET_HOMOLOGUES definitions were 58.58%, 19.58%, and 21.85%, respectively, compared to 56.63%, 10.13%, and 31.38% as assigned by PPanGGOLiN. The differences between the two algorithms were 1.95% for core, 9.45% for shell, and 9.53% for cloud. GET_HOMOLOGUES defines the pangenome compartments as GCs shared across a certain number of genomes, whereas PPanGGOLiN uses both presence and analysis for genomic neighborhoods to infer GCs in the persistent genome, which likely explains these differences.
