## Supplementary material for "Metabolic capabilities are highly conserved among human nasal-associated *Corynebacterium* species in pangenomic analyses": Figure S4

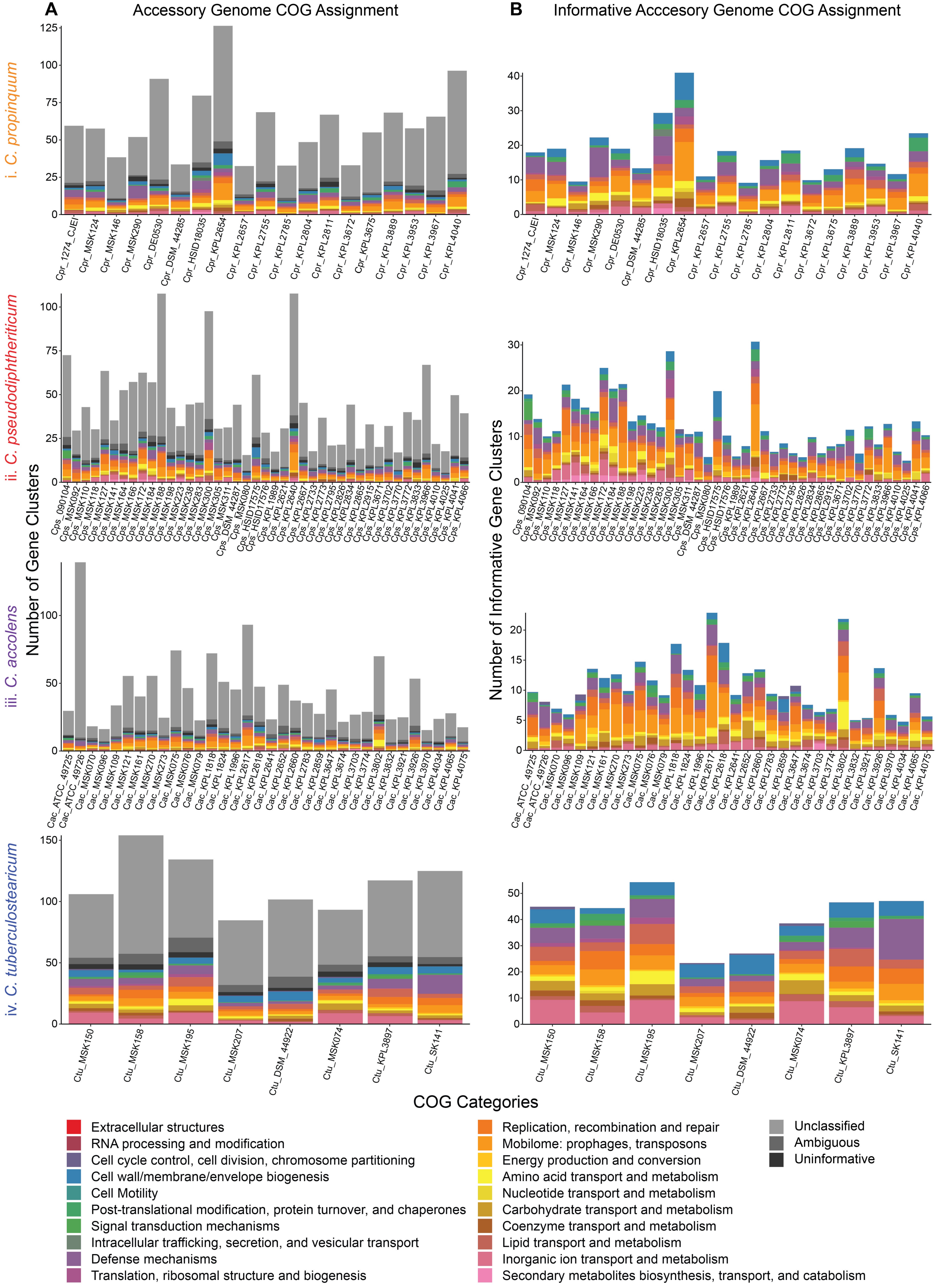

A
Accessory Genome COG Assignment
B
Informative Acccesory Genome COG Assignment
Number of Gene Clusters
Number of Informative Gene Clusters
i. C. propinquum
Cpr_1274_CJEI
Cpr_MSK124
Cpr_MSK146
Cpr_MSK290
Cpr_DE0530
Cpr_DSM_44285
Cpr_HSID18035
Cpr_KPL2654
Cpr_KPL2657
Cpr_KPL2755
Cpr_KPL2785
Cpr_KPL2804
Cpr_KPL2811
Cpr_KPL3672
Cpr_KPL3675
Cpr_KPL3889
Cpr_KPL3953
Cpr_KPL3967
Cpr_KPL4040
ii. C. pseudodiphtheriticum
Cps_090104
Cps_MSK092
Cps_MSK110
Cps_MSK118
Cps_MSK127
Cps_MSK141
Cps_MSK164
Cps_MSK166
Cps_MSK172
Cps_MSK184
Cps_MSK188
Cps_MSK198
Cps_MSK223
Cps_MSK238
Cps_MSK283
Cps_MSK300
Cps_MSK305
Cps_MSK311
Cps_DSM_44287
Cps_MSK080
Cps_HSID17575
Cps_HSID17576
Cps_KPL1989
Cps_KPL2621
Cps_KPL2640
Cps_KPL2667
Cps_KPL2733
Cps_KPL2773
Cps_KPL2795
Cps_KPL2826
Cps_KPL2834
Cps_KPL2865
Cps_KPL2915
Cps_KPL3671
Cps_KPL3702
Cps_KPL3770
Cps_KPL3772
Cps_KPL3833
Cps_KPL3966
Cps_KPL4010
Cps_KPL4025
Cps_KPL4041
Cps_KPL4066
iii. C. accolens
Cac_ATCC_49725
Cac_ATCC_49726
Cac_MSK070
Cac_MSK096
Cac_MSK109
Cac_MSK121
Cac_MSK161
Cac_MSK270
Cac_MSK273
Cac_MSK075
Cac_MSK076
Cac_MSK079
Cac_KPL1818
Cac_KPL1824
Cac_KPL1996
Cac_KPL2617
Cac_KPL2618
Cac_KPL2641
Cac_KPL2652
Cac_KPL2660
Cac_KPL2783
Cac_KPL2859
Cac_KPL3647
Cac_KPL3674
Cac_KPL3703
Cac_KPL3774
Cac_KPL3802
Cac_KPL3832
Cac_KPL3921
Cac_KPL3926
Cac_KPL3970
Cac_KPL4034
Cac_KPL4065
Cac_KPL4075
iv. C. tuberculostearicum
Ctu_MSK150
Ctu_MSK158
Ctu_MSK195
Ctu_MSK207
Ctu_DSM_44922
Ctu_MSK074
Ctu_KPL3897
Ctu_SK141
COG Categories
Extracellular structures
RNA processing and modification
Cell cycle control, cell division, chromosome partitioning
Cell wall/membrane/envelope biogenesis
Cell Motility
Post-translational modification, protein turnover, and chaperones
Signal transduction mechanisms
Intracellular trafficking, secretion, and vesicular transport
Defense mechanisms
Translation, ribosomal structure and biogenesis
Replication, recombination and repair
Mobilome: prophages, transposons
Energy production and conversion
Amino acid transport and metabolism
Nucleotide transport and metabolism
Carbohydrate transport and metabolism
Coenzyme transport and metabolism
Lipid transport and metabolism
Inorganic ion transport and metabolism
Secondary metabolites biosynthesis, transport, and catabolism
Unclassified
Ambiguous
Uninformative
