## Supplementary material for "Metabolic capabilities are highly conserved among human nasal-associated *Corynebacterium* species in pangenomic analyses": Figure S5

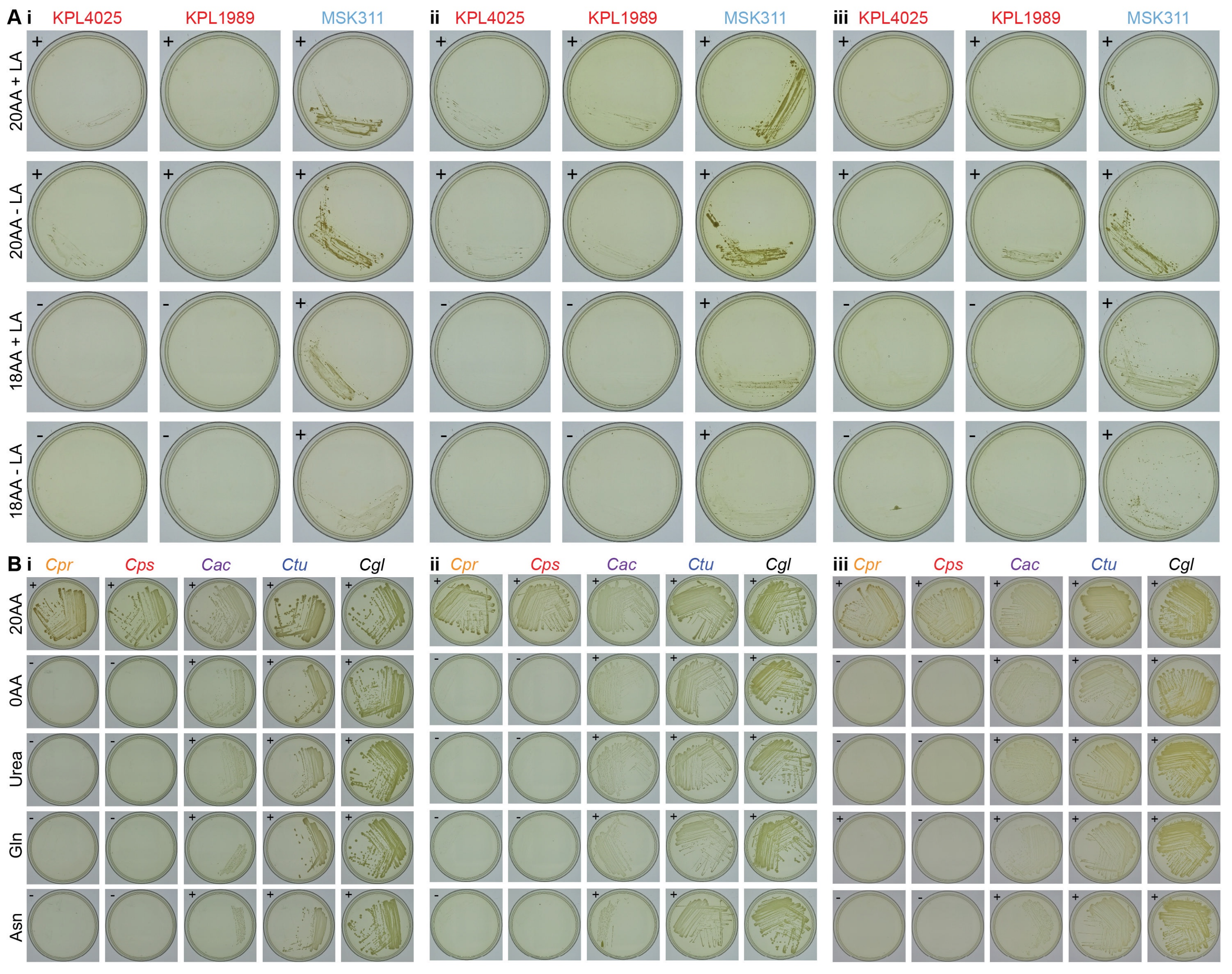

A i
KPL4025
KPL1989
MSK311
ii
KPL4025
KPL1989
MSK311
iii
KPL4025
KPL1989
MSK311
20AA + LA
20AA - LA
18AA + LA
18AA - LA
B i
Cpr
Cps
Cac
Ctu
Cgl
ii
Cpr
Cps
Cac
Ctu
Cgl
iii
Cpr
Cps
Cac
Ctu
Cgl
20AA
0AA
Urea
Gln
Asn

**Figure S5. Results of additional growth experiments from Figures 6 (A) and 7 (B).** (**A**) The Botswanan *C. pseudodiphtheriticum* strain MSK311, which encodes the *fpr2cysIXHDNYZ* operon, grew on chemically defined agarose medium with sulfate as the only source of sulfur (KS-CDM 18AA - LA), whereas the USA strains KPL1989 and KPL4025, which lack this operon, did not. (**B**) *C. accolens* (*Cac*) and *C. tuberculostearicum* (*Ctu*), as well as *C. glutamicum* (*Cgl*), grew on MOPS-buffered CDM agarose medium in the absence of all 20 amino acids, whereas *C*. *propinquum* (*Cpr*) and *C*. *pseudodiphtheriticum* (*Cps*) did not. All images captured after 8 days of growth at 34°C with 5% $CO_2$ with a humidification pan. (+) = growth, (-) = no growth.
